# Spatiotemporal analysis of glioma heterogeneity reveals Col1A1 as an actionable target to disrupt tumor mesenchymal differentiation, invasion and malignancy

**DOI:** 10.1101/2020.12.01.404970

**Authors:** Andrea Comba, Syed M. Faisal, Patrick J. Dunn, Anna E. Argento, Todd C. Hollon, Wajd N. Al-Holou, Maria Luisa Varela, Daniel B. Zamler, Gunnar L Quass, Pierre F. Apostolides, Clifford Abel, Christine E. Brown, Phillip E. Kish, Alon Kahana, Celina G. Kleer, Sebastien Motsch, Maria G Castro, Pedro R. Lowenstein

## Abstract

Intra-tumoral heterogeneity and diffuse infiltration are hallmarks of glioblastoma that challenge treatment efficacy. However, the mechanisms that set up both tumor heterogeneity and invasion remain poorly understood. Herein, we present a comprehensive spatiotemporal study that aligns distinctive intra-tumoral histopathological structures, oncostreams, with dynamic properties and a unique, actionable, spatial transcriptomic signature. Oncostreams are dynamic multicellular fascicles of spindle-like and aligned cells with mesenchymal properties, detected using *ex vivo* explants and *in vivo* intravital imaging. Their density correlates with tumor aggressiveness in genetically engineered mouse glioma models, and high-grade human gliomas. Oncostreams facilitate the intra-tumoral distribution of tumoral and non-tumoral cells, and the invasion of the normal brain. These fascicles are defined by a specific molecular signature that regulates their organization and function. Oncostreams structure and function depend on overexpression of COL1A1. COL1A1 is a central gene in the dynamic organization of glioma mesenchymal transformation, and a powerful regulator of glioma malignant behavior. Inhibition of COL1A1 eliminated oncostreams, reprogramed the malignant histopathological phenotype, reduced expression of the mesenchymal associated genes, induced changes in the tumor microenvironment and prolonged animal survival. Oncostreams represent a novel pathological marker of potential value for diagnosis, prognosis, and treatment.

## INTRODUCTION

High grade gliomas (HGG) are the most prevalent and malignant brain tumors. They grow rapidly, invade surrounding normal brain, and recur within 12 months. Median survival is 18-20 months, in spite of current standard of care^1, 2^. Despite some notable successful outcomes from the large cancer sequencing programs, which identified driver genes in a number of cancers, effective therapeutically actionable breakthroughs have not yet been identified in HGG^3–7^.

HGG are highly heterogeneous at the histological, cellular, and molecular level. Heterogeneity of HGG is illustrated in addition by characteristic pathological structures such as pseudopalisades, microvascular proliferation, and areas of hypoxia and necrosis^2, 8^. The molecular characterization of glioma heterogeneity identified three main molecular signatures: proneural, mesenchymal, and classical^4, 9^. However, later studies demonstrated that all three transcriptomic signatures are expressed within individual tumors^5, 10, 11^. Rather than outright glioma subtypes, the consensus proposes that individual tumors are enriched in particular molecular subtypes. Thus, studies have correlated histological features with genetic alterations and transcriptional expression patterns. For example, highly aggressive histological features such as hypoxic, perinecrotic and microvascular proliferative zones have been associated with the mesenchymal molecular signature and worse prognosis^8^. However, the molecular classification has only minor clinical impact. Thus, alternative classification schemes using a pathway-based classification are currently being considered^12^. How these new classifications will deal with tumor heterogeneity remains to be explored. Moreover, different microenvironmental, metabolic, and therapeutic factors drive transitions of the GBM transcriptomic signature, particularly transitions to mesenchymal states. It is important to note that glioblastoma plasticity explains the high degree of tumor heterogeneity and prompts the selection of new clones at recurrence or therapy resistance^13–16^. It has been established that intra-tumoral heterogeneity is represented by four main cellular states, the progenitor, astrocyte, oligodendrocyte, and mesenchymal like-state, which represent tumor plasticity and are affected by the tumor microenvironment^15^.

Tumoral mesenchymal transformation is a hallmark of gliomas^13, 17, 18^. A mesenchymal phenotype is defined by cells with spindle-like, fibroblast-like morphology associated with alterations in their dynamic cellular organization leading to an increase in cell migration and invasion^19, 20^. The mesenchymal phenotype is controlled by particular transcription factors and downstream genes related to the extracellular matrix (ECM), cell adhesion, migration, and tumor angiogenesis^18, 21, 22^.

However, the cellular and molecular mechanisms that regulate mesenchymal transformation in gliomas, especially concerning the mesenchymal features of invasive cells, has remained elusive. Integrating morphological features, spatially resolved transcriptomics, and cellular dynamics resulting from mesenchymal transformation, growth, and invasion are thus of great relevance to our understanding of glioma progression^13, 20^.

Cell migration is essential to continued cancer growth and invasion. Morphological and biochemical changes that occur during mesenchymal transformation allow glioma cells to move throughout the tumor microenvironment and invade the adjacent normal brain. Tumor cells also migrate along blood vessels, white matter tracks, and the subpial surface. Within the tumor microenvironment, aligned extracellular matrix fibers help guide the movement of highly motile mesenchymal-like cancer cells ^23–25^.

Our study reveals that malignant gliomas, both high grade human gliomas and mouse glioma models, display regular distinctive anatomical multicellular fascicles of aligned and elongated, spindle-like cell. We suggest they are areas of mesenchymal transformation. For the sake of simplicity in their description throughout the manuscript, we have named these areas ‘oncostreams’.

Using time lapse laser scanning confocal imaging *ex vivo*, and multiphoton microscopy *in vivo* of high grade glioma explants we demonstrated that oncostreams are organized collective dynamic structures; they are present at the tumor core and at areas of tumor invasion of the normal brain. Collective motion is a form of collective behavior where individual units’ (cells) movement is regulated by local intercellular interactions (i.e., attraction/repulsion) resulting in large scale coordinated cellular migration ^26–30^. Collective motion plays an essential role in embryogenesis and wound healing^27, 31–33^. Emergent organized collective motion patterns could help explain so far poorly understood tumoral behaviors such as invasion, metastasis, and especially recurrence^27, 31^. Studies of tumor motility have concentrated on the behavior of glioma cells at the tumor invasive border^27, 34, 35^. Potential motility at the glioma core has not been studied in much detail so far. This study challenges the conventional belief that cells in the central core are non-motile and indicate that the glioma core displays collective migratory patterns. This would suggest that the capacity of gliomas to invade and grow, results from phenomena occurring at the tumor invasive border, and from the overall capacity of gliomas to organize collective motion throughout the tumor mass, from the tumor core to the tumor invasive border.

To study the molecular mechanisms underlying oncostream organization and function we used laser capture microdissection (LCM) followed by RNA-sequencing and bioinformatics analysis. We discovered that oncostreams are defined by a mesenchymal transformation signature enriched in extracellular matrix related proteins, and which suggest that Collagen1A1 (COL1A1) is a key determinant of oncostream organization. Inhibition of COL1A1 within glioma cells lead to oncostream loss and reshaping of the highly aggressive phenotype of HGG. These data indicate that COL1A1 is likely to constitute the tumor microenvironment scaffold, and to serve to organize areas of collective motion in gliomas.

COL1A1 has been shown previously to be a major component of the extracellular matrix in different cancers, including glioma, and has been reported to promote tumor growth and invasion^36, 37^. Alternatively, some data suggest that collagen fibers could be passive barriers to resist tumor cell infiltration or provide biophysical and biochemical support for cell migration. Some studies reported that density of COL1A1 inversely correlates with glioma patient’s prognosis. However, other studies showed that either increased or decreased deposition of collagen could be associated with increased tumor malignancy^38–40^. Therefore, it is important to further determine the role of COL1A1 in glioma invasion and continued growth.

This study provides a comprehensive study of the histological, morphological, and dynamic properties of glioma tumors. In addition, we uncover a novel characterization of the molecular mechanisms that define intra-tumoral mesenchymal transformation in gliomas and discuss their therapeutic implications. Oncostreams are anatomically and molecularly distinctive, regulate glioma growth and invasion, display collective motion, and are regulated by the extracellular matrix, specially by COL1A1. Inhibiting COL1A1 within glioma cells is a potential therapeutic strategy to mitigate glioma mesenchymal transformation, intra-tumoral heterogeneity, and thus, reduce deadly glioma invasion and continued growth.

## RESULTS

### Intra-tumoral multicellular fascicles of elongated and aligned cells in gliomas: oncostreams

High grade gliomas (HGG) are characterized by anatomical, cellular and molecular heterogeneity which determines, in part, tumor aggressiveness and reduces treatment efficacy^5, 7, 11^. Histopathological analysis of mouse and human gliomas revealed the presence of frequent distinct multicellular fascicles of elongated (spindle-like) and aligned cells (≈5-30 cells wide) distributed throughout the tumors. These structures resemble areas of mesenchymal transformation which we describe as “oncostreams*”* (Fig. 1A-B).

**Fig. 1.**
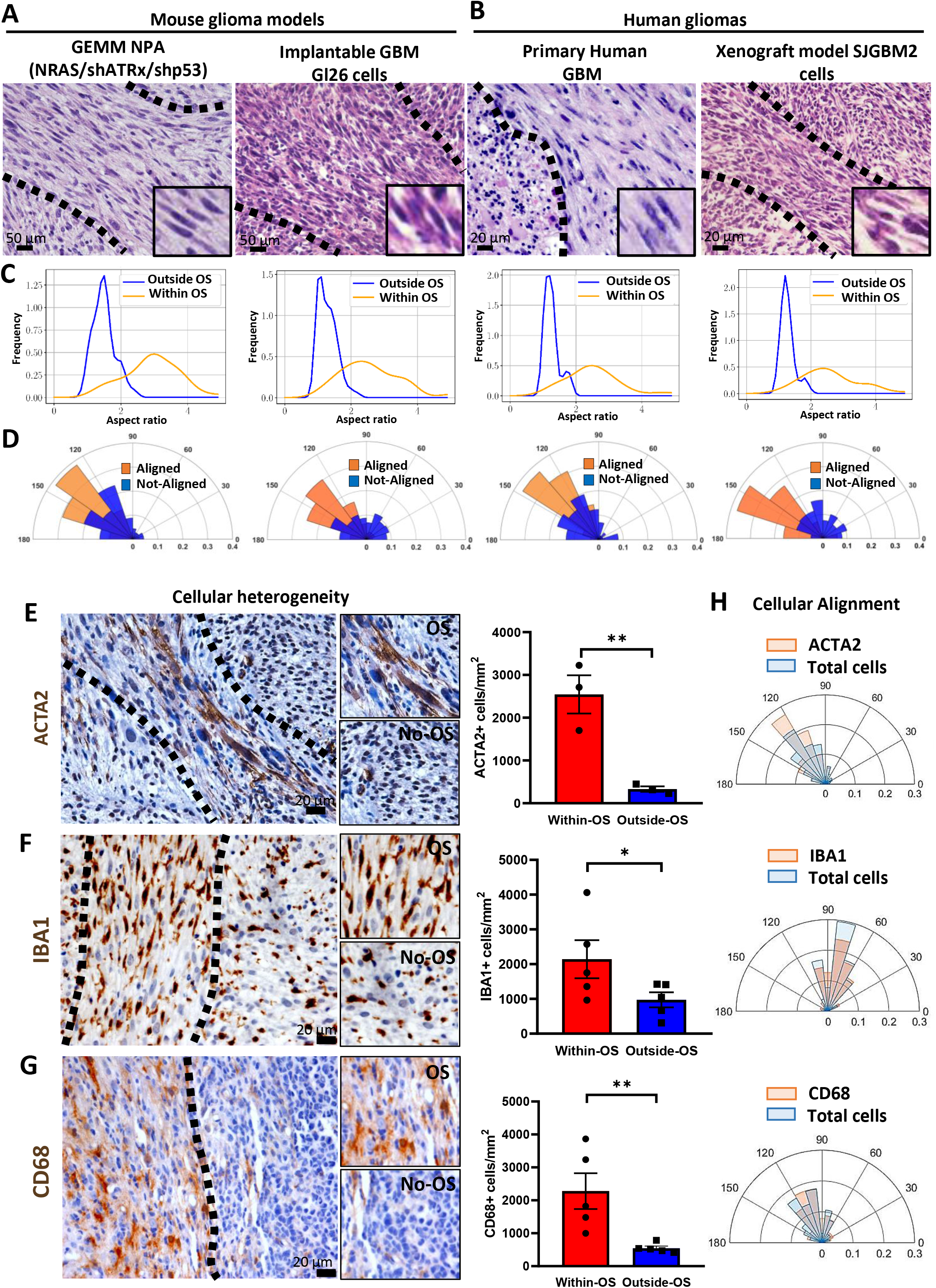
Oncostreams are multicellular fascicles present in mouse and human gliomas. **A)** Representative 5 µm H&E microtome sections from gliomas showing that fascicles of spindle-like glioma cells (oncostreams, outlined by the dotted line) are present in a GEMM of gliomas NPA (NRAS/shATRx/shp53) and the GL26 intracranial implantable model of glioma. Scale bars: 50 μm. **B)** Representative H&E microtome sections of human glioma and human xenografts showing the presence of oncostreams. Scale bar: 20 μm. **C-D)** Histograms showing the cellular shape analysis (aspect ratio) (**C**) and angle orientation (alignment) for the corresponding images (**D**) show areas of oncostreams (OS) formed by elongated and aligned cells and areas with no oncostreams (No-OS) as rounded and not-aligned cells. **E-G)** Immunostaining shows that tumor cells, mesenchymal cells (ACTA2+), microglia/macrophages (IBA1+ and CD68+), are aligned within, the main orientation axis of oncostreams. Bar graphs show the quantification of ACTA+, n=3 (**E**), IBA1+, n=5 (**F**) and CD68+, n=5 (**G**) cells within oncostreams areas in NPA tumors. 6-13 areas of oncostreams per tumor section per animal were imaged. Scale bar: 20 μm. Error bars represent ± SEM; unpaired t-test analysis, *p<0.05, **p<0.001. **H)** Angle orientation shows the alignment of ACTA+, IBA1+ and CD68+ cells within oncostreams for the corresponding images.

To study the presence and morphological characterization of oncostreams, we examined histological sections from various mouse glioma models as well as human glioma specimens (Fig. 1 A-B). We determined the existence of oncostreams in genetically engineered mouse models (GEMM) of glioma including NPA (Nras, shP53, shATRx) and NPD (Nras, shP53, PDGFβ) and other implantable models (GL26) (Fig. 1A **and Supplementary Fig. 1A-B**). Moreover, human glioma samples from primary resections and a xenograft glioma model, SJGBM2, established the presence of these multicellular structures in human tissue (Fig. 1B **and Supplementary Fig. 1C**). Morphological analysis determined that cells within histological areas corresponding to oncostreams have an aspect ratio of 2.63±0.19 (elongated or spindle-like cells) compared to the surrounding tissue where cells have an aspect ratio of 1.37±0.12 (round cells), both in mouse and human gliomas as shown in Fig. 1C **and Supplementary Fig. 1D.** We also determined that elongated cells within oncostreams are nematically aligned with each other, whereas outside of oncostreams, cell orientations are not aligned (Fig. 1D **and Supplementary Fig. 1E).**

To gain insight into the cellular features of oncostreams we asked if they are homogeneous or heterogeneous multicellular structures. We observed that in GEMM of gliomas, oncostreams are formed by GFP+ tumor cells, and are enriched in other tumor microenvironment cells such as ACTA2+ mesenchymal cells, Iba1+ and CD68+ tumor associated microglia/macrophages cells, Nestin+ cells and GFAP+ glial derived cells (Fig. 1, E-G**, and Supplementary Fig. 2 A-D**). The quantification of mesenchymal cells (ACTA2+), and tumor associated microglia/macrophages (TAM) cells (CD68+ and Iba1+) showed a significant enrichment of these populations within oncostreams compared to the surrounding areas (Fig. 1E-G). Moreover, non-tumoral cells within oncostreams were positively aligned along the main axes of oncostreams, and with tumor cells in mouse gliomas (Fig. 1H). This suggests that oncostreams are mesenchymal-like structures which interact with TAM and mesenchymal cells.

To test if oncostreams form along existing brain structures, we evaluated their co-localization with white matter tracts. Although, occasional positive immune-reactivity (Neurofilament-L) was present within some areas of the tumors, oncostream fascicles were not preferentially organized along brain axonal pathways **(Supplementary Fig. 1F)**. These data indicate that oncostreams are fascicles of spindle-like aligned cells within glioma tumors, which contain tumor and non-tumor cells.

### Oncostream density positively correlates with tumor aggressiveness and poor prognosis in mouse and human gliomas

Oncostreams are unique histological features that contribute to intra-tumoral heterogeneity suggesting a potential role in glioma progression and malignancy. To understand whether the presence of oncostreams correlates with tumor aggressiveness and clinical outcomes, we generated genetically engineered tumors of different malignant behaviors using the Sleeping Beauty Transposon system. These models reproduce the malignant histopathological features of gliomas as demonstrated in previous studies^41–44^. We induced tumors harboring two different genotypes: (1) Activation of RTK/RAS/PI3K pathway, in combination with p53 and ATRX downregulation (**NPA**), and, (2) RTK/RAS/PI3K activation, p53 downregulation, ATRX downregulation, and mutant IDH1-R132 expression (**NPAI**) (Fig. 2A). IDH1-wild-type tumors (**NPA**) display a highly malignant phenotype and worse survival prognosis (Mediam survival (MS): 70 days), compared with tumors harboring the IDH1-R132R mutation, **NPAI,** (MS: 213 days) (Fig. 2B). This outcome reproduces human disease, as patients with IDH1-mutant tumors also have prolonged median survival^1, 44, 45^. Tumor histopathological analysis showed a positive correlation between the density of oncostreams and tumor malignancy (Fig. 2C-D). NPA (IDH1-WT) tumors exhibited larger areas of oncostreams within a highly infiltrative and heterogeneous glioma characterized by abundant necrosis, microvascular proliferation, pseudopalisades and cellular heterogeneity as described before ^43, 44^. Conversely, NPAI (IDH1-Mut) tumors display a very low density of oncostreams and a homogenous histology mainly comprised of round cells, low amounts of necrosis, no microvascular proliferation, absence of pseudopalisades and less invasive borders (Fig. 2C **and Supplementary Fig. 5).**

**Fig. 2.**
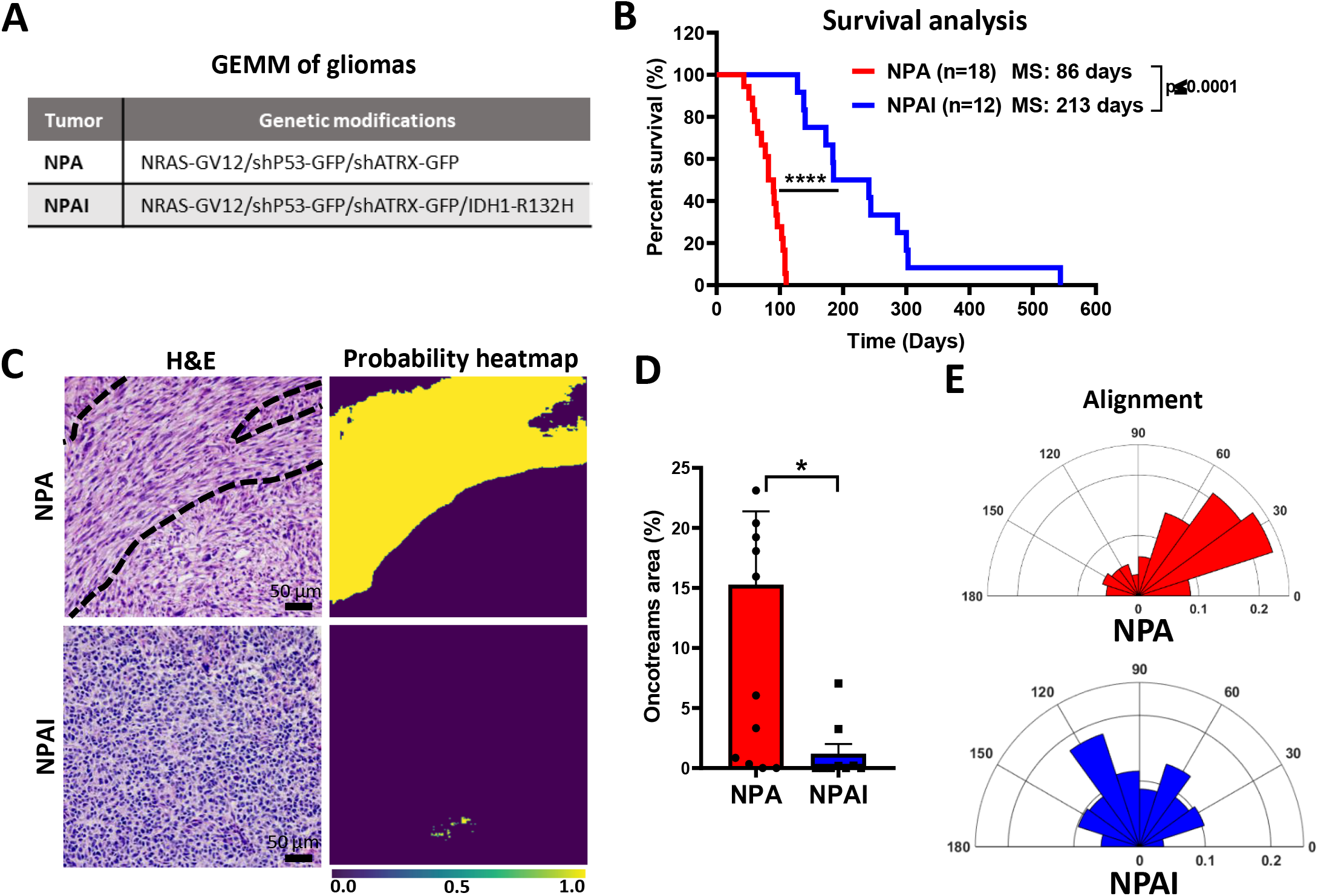
Oncostreams density positively correlates with tumor aggressiveness in GEMM of gliomas. **A)** Genetic makeup of NPA and NPAI tumors. **B)** Kaplan–Meier survival curves of NPA and NPAI mouse gliomas show that animals bearing IDH1-R132H mutant tumors (NPAI) have prolonged median survival (MS): **NPA** (MS: 86 days; n: 18) versus **NPAI** (MS: 213 days; n:12). Log-rank (Mantel-Cox) test; ****p<0.0001. **C-D)** Deep learning method for oncostream detection in H&E stained mouse glioma sections: **C)** Representative images of oncostreams manually segmented on H&E stained sections of NPA gliomas and NPAI tumors. The output of our trained model for each image is shown below (probability heat maps), for tissues containing oncostreams (NPA), and without oncostreams (NPAI), scale bar = 50 μm. **D)** 10-14 random fields per tumor section per animal were imaged, n=9 NPA and n=12 NPAI, and quantified using deep learning analysis. Error bars represent ± SEM; unpaired t-test analysis, *p<0.05. **E)** Angle histogram plots show aligned cells in NPA tumors vs non-aligned cells in NPAI tumors for the representative images showed in figure.

Further, to objectively identify and quantify tumor areas covered by oncostreams, we trained a fully convolutional neural network (fCNN) (**Supplementary Fig. 3** and 4A). Our deep learning analysis found that oncostreams occupied 15.28 ± 6.10% of the area in NPA tumors compared with only 1.18 ± 0.81 % in NPAI tumors (Fig. 2C and D**, and Fig. Supplementary 5A and B**). Cellular alignment analysis validated the presence or absence of oncostreams (Fig. 2E).

To determine whether oncostreams are linked to glioma aggressiveness in human patients, we evaluated a large cohort of TCGA glioma diagnostic tissue slides from the Genomic Data Commons Portal of the National Cancer Institute. We visually examined 100 TCGA-glioblastoma multiforme tissue sections (WHO Grade IV) and 120 TCGA-low grade glioma tissues (WHO Grade II and III) using the portal’s slide image viewer (**Supplementary Table 1**). Oncostreams were present in 47% of TCGA-GBM grade IV tumors tissue, in 8.6 % of TCGA-LGG grade III, and were absent from TCGA-LGG grade II (Fig. 3A-C **and Supplementary Table 2**), consistent with tumor aggressiveness (http://gliovis.bioinfo.cnio.es)^46^. We then determined the presence of oncostreams across known molecular subtypes of HHG (Grade IV)^4^. We found oncostream fascicles in 59.4% of Mesenchymal (MES), 53.6% of Classical (CL) subtypes and only 26.7% of Proneural (PN) (**Fig. Supplementary 6A**). Finally, we evaluated oncostreams presence related to IDH status and 1p 19q co-deletion in LGG (Grade III). Oncostreams were present in 16.6% of IDH-WT subtype, 5% of IDHmut-non-codel and absent from IDHmut-codel subtype (**Fig. Supplementary 6B)**. These analyses suggest that oncostream presence is higher in Mesenchymal and Classical subtypes and correlates with IDH-WT status, and thus with a poor prognosis.

**Fig. 3.**
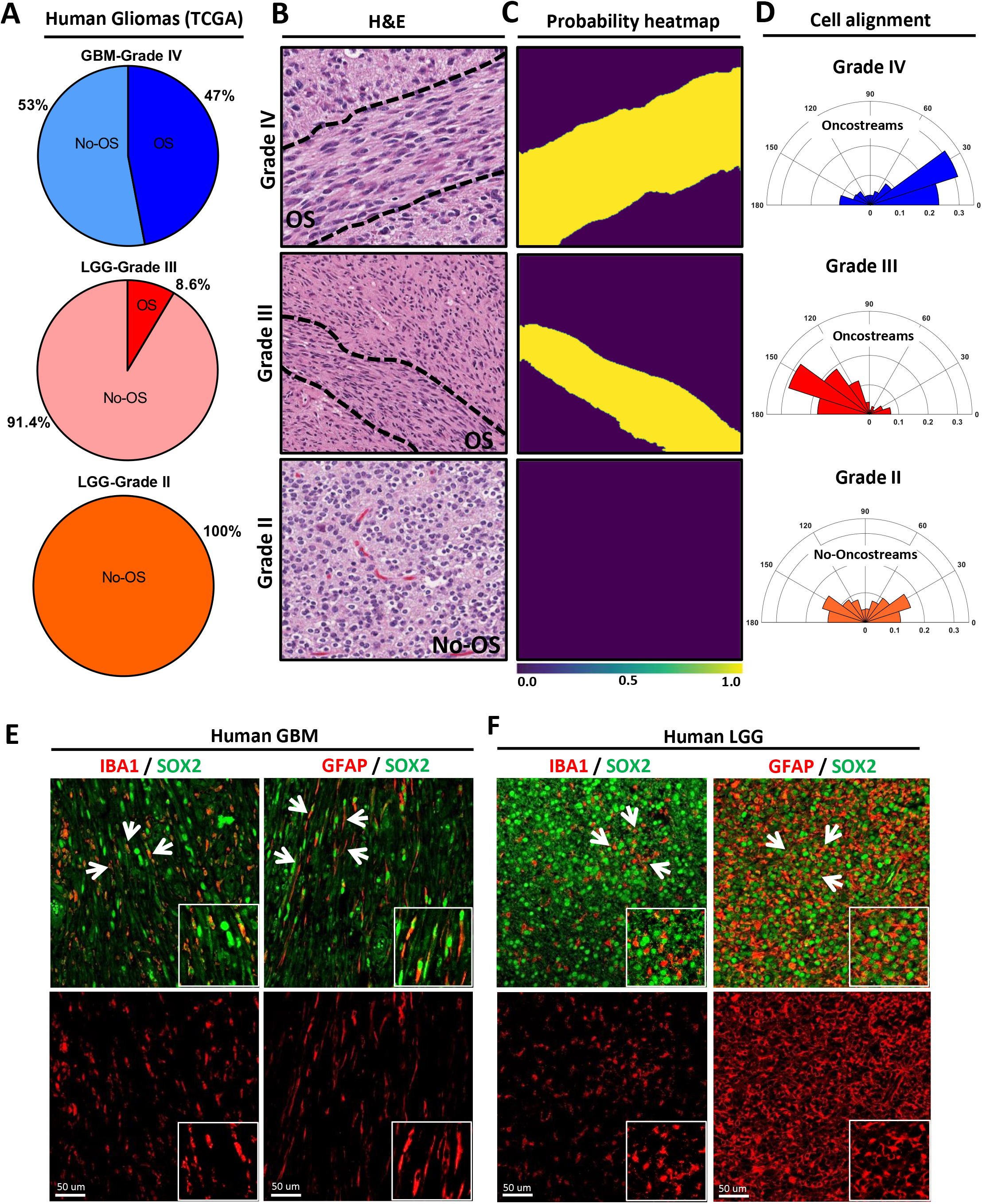
The density of oncostreams positively correlates with tumor aggressiveness in human gliomas. **A)** TCGA tumors were analyzed from different grade: GBM-Grade IV (100 tumors). LGG-Grade III (70 tumors) and LGG-Grade II (50 tumors). Pie charts show percentage of tumors displaying oncostreams in relation to tumor grade. Oncostreams are present in 47% of GBM grade IV tumors, 8.6 % of LGG grade III, and are absent from LGG grade II. **B)** Manual identification of oncostreams in H&E images are shown for human gliomas with WHO grades IV, III, II from TCGA. **C)** Deep learning analysis for human gliomas. Our algorithm was able to detect oncostreams in grade IV and III gliomas but not in grade II gliomas. **D)** Angle histogram plots show the alignment of cells in H&E histology sections of Grade IV and Grade III gliomas’ oncostreams and random alignment in grade II glioma sections lacking oncostreams. Angle histogram correspond to the representative images. **E-F)** Immuno-fluorescence staining of SOX2+ tumor cells (green), glial fibrillary acidic protein (GFAP+) cells (red), and microglia/macrophage (IBA1+) cells (red) in high-grade human glioblastoma (GBM) (WHO Grade IV), IDH-WT **(E)** and in low-grade glioma (LGG) (WHO Grade III), IDH-mutant **(F)**, showing oncostreams heterogeneity and cellular alignment of these cells in human high-grade gliomas but not in low grade gliomas (arrows). Scale bars: 50 µm.

To validate the histological identification, we examined H&E images using our deep learning algorithm (**Fig. Supplementary 4B**). We observed a strong concordance (>84%) between machine learning and the manual histological identification of oncostreams (**Table Supplementary 3)**. Oncostream presence and their segmentation by deep learning is illustrated in Fig. 3C **and Supplementary Fig. 7** and **8**. Additionally, alignment analysis of glioma cells confirmed the existence of fascicles of elongated, mesenchymal-like cells in human gliomas (Fig. 3D). Thus, our deep learning algorithm validates our histological identification of oncostreams and confirms that the density of oncostream fascicles positively correlates with glioma aggressiveness.

The analysis of cellular heterogeneity showed that non-tumoral cells such as Iba1+ macrophages/microglia and GFAP+ glial derived cells were positively aligned within oncostreams tumoral cells (SOX+) in human HGG (Fig. 3E). Conversely, we detected that low grade gliomas (LGG) exhibited homogenous round cells, GFAP+ and Iba1+ cells throughout the tumor with no defined orientation or alignment (Fig. 3F).

### Oncostreams are defined by a distinctive spatial transcriptome signature

To determine whether oncostreams fascicles are characterized by a specific gene expression profile, we performed a spatially-resolved transcriptomic analysis using laser capture microdissection (LCM) coupled to RNA sequencing (RNA-Seq). Oncostreams were dissected according to their morphological characteristics defined above. Surrounding areas of homogenous rounded cells were selected as non-oncostreams areas (control) (Fig. 4A). RNA-Seq analysis detected a set of 43 differentially expressed (DE) genes; 16 genes were upregulated and 27 downregulated within oncostreams (Fig. 4 B-C **and Table Supplementary 4**).

**Fig. 4.**
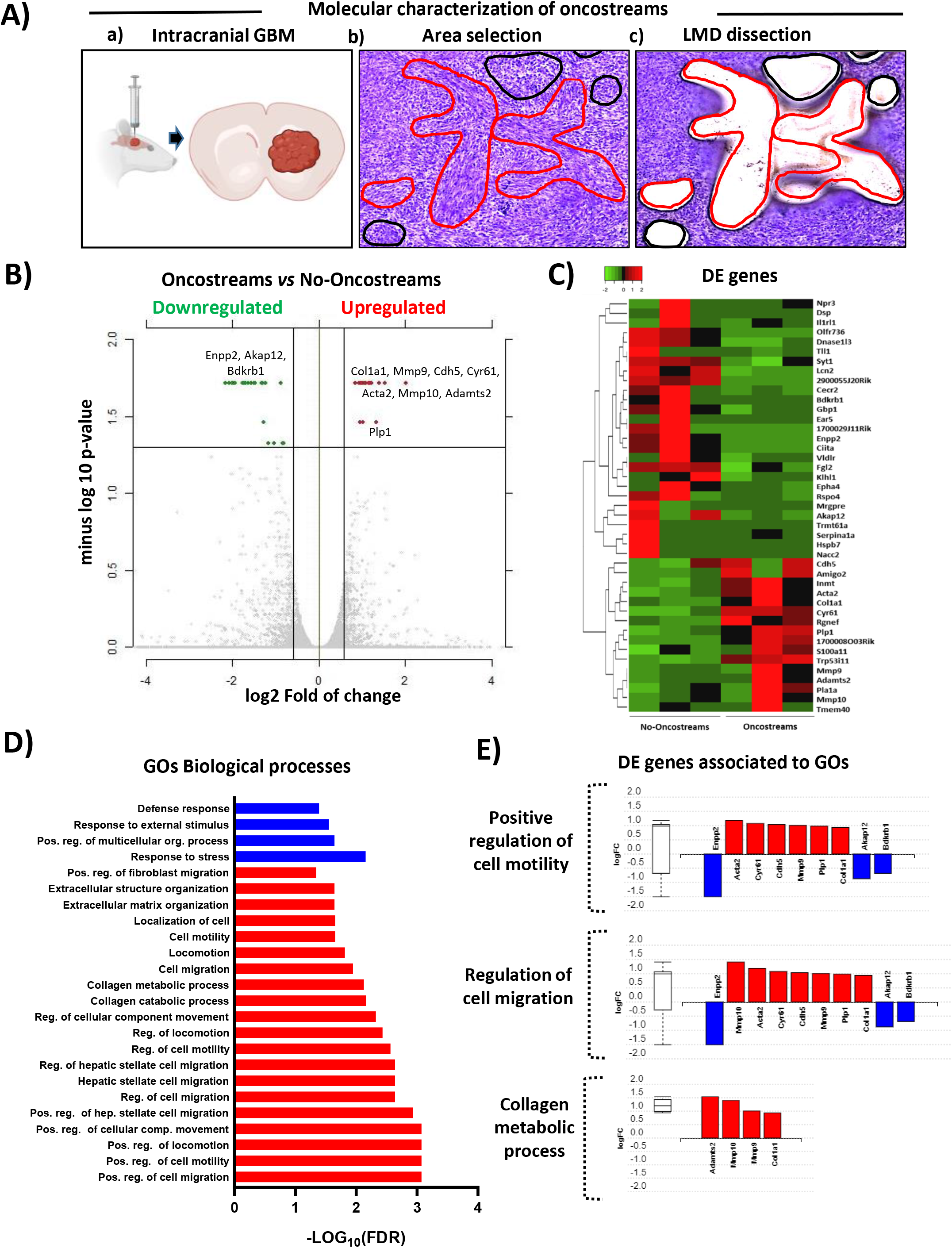
Oncostreams are defined by a unique gene expression signature related to mesenchymal transformation and migration. **A)** (a) Schematic representation of spatial transcriptomic analysis of glioma oncostreams using Laser Capture Microdissection (LCM). Glioma tumors were generated by intracranial implantation of NPA tumor cells in C57BL6 mice. (b-c) Oncostream areas (red outline) were identified and dissected from surrounding glioma tissue (black outline) in mouse glioma samples using a LCM microscope. **B)** A volcano plot displays differentially expressed (DE) genes from oncostream *vs* no-oncostream areas. DE genes were selected based on a fold change of ≥ 1.5 and a q-value (false discovery rate (FDR) corrected p-value) of ≤ 0.05. Upregulated genes (red dots) and downregulated genes (green dots) are shown. Relevant genes related to mesenchymal migration are labeled on the graph. **C)** Heat map illustrates gene expression patterns for oncostream vs no-oncostream areas in NPA glioma tumors (n=3 biological replicates/group). Differentially upregulated genes (16) are represented in red and downregulated genes (n=27) are represented in green (q-value ≤ 0.05 and fold change ≥ ± 1.5). **D)** Functional enrichment analysis of overrepresented GO terms (biological processes) obtained when comparing oncostream vs no-oncostream DE genes. p-value corrected for multiple comparisons using the FDR method. Cutoff FDR<0.05. Blue: Downregulated GOs. Red: upregulated GOs. **E)** Bar graphs show DE genes annotated to the most relevant enriched GOs biological process: “Positive regulation of cell motility”, “Regulation of cell migration” and “Collagen metabolic process.”

Functional enrichment analysis of DE genes, performed using the I-PathwayGuide platform (Advaita Corporation, MI, USA), showed that False Discovery Rate (FDR) corrected gene ontology (GOs) terms were associated with migration and extracellular matrix biological process. GOs such as “positive regulation of motility” “positive regulation of cell migration”, “collagen catabolic processes” and “extracellular matrix organization” were the most over-represented biological processes (Fig. 4D **and Table Supplementary 5**). The upregulated DE genes within the relevant GOs include: COL1A1, MMP9, MMP10, ACTA2, ADAMTS2, CDH5, CYR61, PLP1 and those downregulated were ENPP2, AKAP12, BDKRB1 (Fig. 4E **and Fig. Supplementary 9**). Significant DE genes shared by related GOs are shown in **Supplementary Fig. 9**. These data indicate that oncostreams can be identified by a specific gene expression set and suggest a distinct role for oncostreams as intra-tumoral mesenchymal-like migratory assemblies within glioma tumors.

### COL1A1 contributes to oncostream organization in high-grade gliomas

Histopathologically, oncostreams are spindle-like multicellular fascicles with a defined DE gene expression signature enriched in mesenchymal genes. The GO ontology analyses suggests a central role of collagen catabolic process and extracellular matrix organization in oncostreams function. To understand the molecular mechanisms that regulate oncostream organization and function, we identified critical genes using network analysis. Network interactions revealed that COL1A1 is a hub gene, one of the most highly connected nodes, representing a potential regulator of the network’s signaling pathways and biological functions (Fig 5A **and Fig. Supplementary 10A**). We found that the most relevant COL1A1 related pathways include: Focal Adhesion, Extracellular Matrix Organization and Integrin Signaling pathways (**Fig. Supplementary 10B-C and Table Supplementary 7 and 8**).

**Fig. 5.**
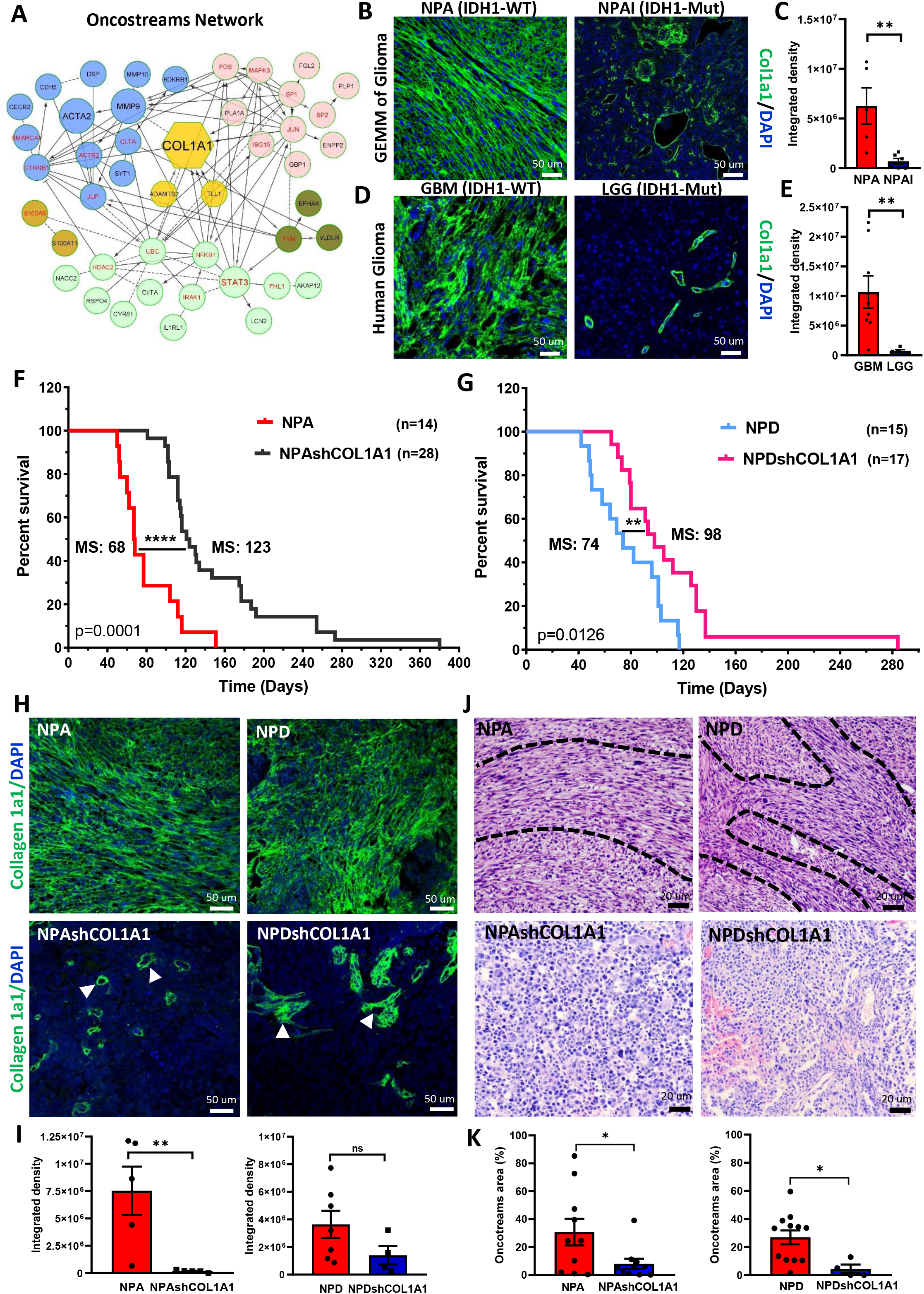
COL1A1 is a central hub in oncostream organization and glioma malignancy. **A)** Network analysis of the DE genes comparing oncostreams versus no-oncostreams DE genes. Genes with a higher degree of connectivity are highlighted with larger nodes. Clusters of nodes with the same color illustrate modules of highly interacting of genes in the network. **B)** Immunofluorescence analysis of COL1A1 expression in GEMM of glioma tissues comparing NPA (IDH1-WT) vs NPAI (IDH1mut). Representative confocal images display COL1A1 expression in green (Alexa 488) and nuclei in blue (DAPI). Scale bar: 50 μm. **C)** Bar graphs represent COL1A1 quantification in terms of fluorescence integrated density. NPA n=5 and NPAI n=6 animals for each experimental condition were used for the analysis. Ten fields of each tumor section were selected at random. Error bars represent ±SEM. t-test, **p<0.01. **D)** Immunofluorescence analysis of COL1A1 expression in human GBM and LGG tumors. COL1A1 expression in green (Alexa 488) and nuclei in blue (DAPI). Scale bar: 50 μm. **E)** Bar graphs represent COL1A1 quantification as fluorescence integrated density. 5 (LGG) and 8 (GBM) tumor samples were used for the analysis. Ten fields of each tumor section were selected at random. Error bars represent ±SEM. t-test, **p<0.01. **F-G)** GEMM of glioma with COL1A1 inhibition. **F)** Kaplan–Meier survival curve comparing NPA (MS: 68 days; n: 14) vs NPAshCOL1A1 (MS: 123 days; n: 28) **G)** Kaplan–Meier survival curve comparing NPD (MS: 74 days; n=15) versus NPDshCOL1A1 (MS: 98 days; n=17). Log-rank (Mantel-Cox) test. **** p<0.0001, **p<0.0126. **H)** Immunofluorescence analysis of COL1A1 expression in GEMM of glioma controls (NPA and NPD) and Col1A1 downregulation (NPAshCOL1A1 and NPDshCOL1A1). Representative confocal images of COL1A1 expression in green (Alexa 488) and nuclei in blue (DAPI). Arrows indicate COL1A1 enriched perivascular cells. Scale bar: 50 μm. **I)** Bar graphs represent COL1A1 quantification in terms of fluorescence integrated density. 5-7 tumor samples for each experimental condition were used for the analysis. Ten fields of each tumor section were selected at random. Error bars represent ±SEM. t-test, **p<0.01. **J)** Representative images of the histopathological identification of oncostreams in H&E tissue sections comparing the COL1A1 knockdown tumors with their respective controls. Scale bars: 50 μm. **K)** Quantitative analysis of oncostream areas using deep learning analysis. 4-12 tumor samples for each experimental condition were used for the analysis. Error bars represent ± SEM; unpaired t-test analysis, *p<0.05.

To analyze the role of COL1A1 in oncostream organization, we analyzed COL1A1 expression by immunofluorescence analysis. The COL1A1 gene encodes for the alpha-1 chain of type I collagen fibers. We observed that collagen fibers were aligned within oncostreams and overexpressed in more aggressive NPA (IDH1-WT) gliomas compared with NPAI (IDH1-Mut) tumors. COL1A1 expression was significantly lower and only found surrounding blood vessels in NPAI (IDH-Mut) tumors (Fig. 5B-C). Correspondingly, human GBM glioma tumors (IDH1-WT) with high oncostream densities showed prominent alignment of collagen fibers along these fascicles and higher COL1A1 expression compared to LGG (IDH1-Mut) (Fig. 5D and E).

Moreover, TCGA-glioma data indicate that COL1A1 has differentially higher expression in GBM histological Grade IV. LGG IDH-WT tumors display higher expression of COL1A1 than IDH1-Mutant. Within the GBM molecular subtype classification^4, 9^, the Mesenchymal group shows higher expression of COL1A1 than the Proneural and Classical groups (**Fig. Supplementary 11A**); the COL1A1 gene is clearly associated with the mesenchymal subtype. Analysis of patient survival related to COL1A1 expression showed that mesenchymal GBM subtype displayed a significantly shorter survival (MS: 10.4 months) for COL1A1 high tumors, compared to COL1A1 low (MS: 17.9 months) tumors. Classical and Proneural subtypes did not show survival differences associated to COL1A1 expression (**Fig. Supplementary 11B**). Thus, oncostreams represent intra-tumoral mesenchymal-like structures organized along collagen fibers.

### COL1A1 depletion leads to oncostream loss, tumor microenvironment (TME) remodeling and increases in median survival

To evaluate the functional role of COL1A1 in oncostream formation we generated a COL1A1-deficient genetically engineered mouse glioma model. We generated COL1A1 wildtype, and COL1A1 knock-down tumors with different genetic backgrounds (**Fig Supplementary 12A-C**). COL1A1 downregulation increased median survival (MS) (Fig. 5F and G). The knockdown of COL1A1 in NPA tumors (NPAshCOL1A1) increased survival to MS: 123 days, compared to NPA control tumors (MS: 68 days) (Fig. 5F). Similarly, COL1A1 knockdown in NPD tumors harboring PDGFβ ligand upregulation (NPDshCOL1A1), also exhibited an increased median survival (MS: 98 days) compared to the NPD controls (MS: 74 days) (Fig. 5G).

To further analyze the effects of COL1A1 downregulation, we evaluated the histopathological features of glioma tumors, quantified oncostream density using deep learning analysis and evaluated COL1A1 expression within glioma tissues (Fig. 5 H-I). We observed that NPA tumors with COL1A1 downregulation showed a significant reduction of COL1A1 immunoreactivity within tumors; it was only maintained in small areas surrounding blood vessels (Fig. 5J-K). COL1A1 inhibition led to oncostream loss and reprogramming of the histopathological tumoral characteristics as evidenced by homogenous round cell morphology, resembling low grade tumors (Fig. 5J-K). Downregulation of COL1A1 in NPD tumors appeared less effective, with large areas of remaining COL1A1 (**Fig. %H-I**). Nonetheless, COL1A1 was downregulated within tumor cells and oncostream dismantling was significant compared to NPD control. Some oncostream areas remained associated with blood vessels which displayed significant amounts of COL1A1 (Fig. 5J-K).

We analyzed the effect of COL1A1 depletion on the intrinsic properties of tumoral cells. In vitro studies showed that COL1A1-knockdown cells exhibited a significantly decreased cell proliferation and cell migration compared to controls (**Supplementary Fig. 13 A-D**). Also, we observed that intracranial implantation of COL1A1-knockdown cells resulted in decreased tumor growth and progression when compared to controls (**Fig. Supplementary 13E**). *In vivo*, genetically engineered COL1A1 knockdown tumors displayed decreased cell proliferation (PCNA+ cells) (Fig. 6A-B), increased apoptosis via activation of Cleaved-Caspase 3, and downregulation of the anti-apoptotic protein Survivin **(Supplementary Fig. 14A-C**).

**Fig. 6.**
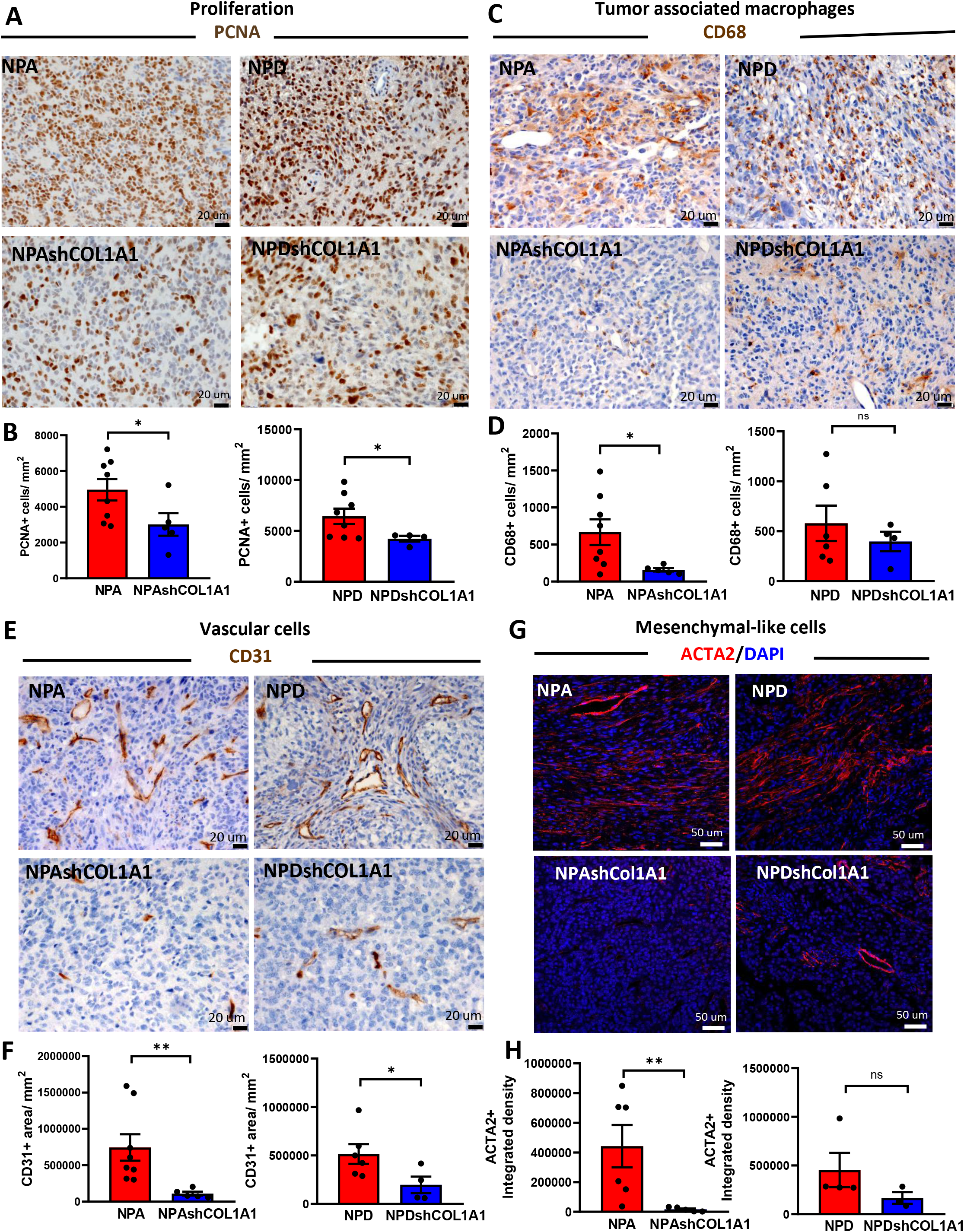
Knockdown of COL1A1 within glioma cells modifies the tumor microenvironment. Immunohistochemical analysis (**A**, **C** and **E**) of GEMM of glioma controls (**NPA** and **NPD**) and COL1A1 downregulation (**NPAshCOL1A1** and **NPDshCOL1A1**). **A)** Representative images of PCNA expression. Scale bar: 20 μm. **B)** Bar graphs represent the quantification of PCNA+ cells numbers (cells/mm^2^) using QuPath positive cell detection. Error bars represent ±SEM, (NPA: n=8, NPAshCOL1A1: n=5, NPD: n=8, NPDshCOL1A1: n=4), t-test, *p<0.05. **C)** Representative images of CD68 expression. Scale bar: 20 μm**. D)** Bar graphs represent CD68+ cell quantification (cells/mm^2^) using QuPath positive cell detection. Error bars represent ±SEM, (NPA: n=8, NPAshCOL1A1: n=5, NPD: n=6, NPDshCOL1A1: n=4), t-test, *p<0.05, ns: no significant. **E)** Representative images of CD31 expression. Scale bar: 20 μm**. F)** Bar graphs represent CD31+ cells quantification (cells/mm^2^) using QuPath positive cells detection. Error bars represent ±SEM, (NPA: n=8, NPAshCOL1A1: n=5, NPD: n=6, NPDshCOL1A1: n=4), t-test, **p<0.01, *p<0.05. **G)** Immunofluorescence analysis of GEMM of glioma controls (**NPA** and **NPD**) and COL1A1 downregulation (**NPAshCOL1A1** and **NPDshCOL1A1**). Representative images of ACTA2 expression in red (Alexa 555) and nuclei in blue (DAPI). Scale bar: 50 μm. **H)** Bar graphs represent ACTA2 quantification in terms of fluorescence integrated density. Error bars represent ±SEM, (NPA: n=6, NPAshCOL1A1: n=5, NPD: n=4, NPDshCOL1A1: n=3), t-test, **p<0.01, ns: no significant.

Furthermore, to determine whether COL1A1 downregulation within glioma cells modifies the glioma TME we analyzed changes in tumor associated macrophages (TAM), endothelial cells and mesenchymal cells. We found that COL1A1 knockdown tumors exhibited a decreased recruitment of CD68+ TAM (Fig. 6C-D), impaired CD31+ endothelial vascular proliferation (Fig. 6E-F) and diminished ACTA2+ perivascular mesenchymal cells (Fig. 6G-H). Moreover, inhibition of COL1A1 within glioma cells led to downregulation of fibronectin expression, a mesenchymal associated extracellular matrix protein (**Fig. Supplementary 14E-F**) that is associated with a more aggressive phenotype.

These preclinical animal models knocked down the expression of COL1A1 from the earliest stages of tumor development. Further, to evaluate the effects of the pharmacological degradation of deposited collagen fibers in highly malignant tumors we analyzed explants of brain tumor sections treated with collagenase. We observed that collagenase treatment decreased reticular fibers (general collagen staining), reduced COL1A1 expression and disassemble fibers’ alignment along tumoral cells and caused oncostreams depletion in a dose dependent manner (**Fig. Supplementary 15A-D**). These data indicate that oncostream organization and functions are regulated by COL1A1. COL1A1 knockdown within glioma cells decreased oncostream formation, reprogramed glioma mesenchymal transformation and remodeled the glioma TME, thus increasing animal survival. COL1A1 inhibition represents a novel approach for future translational development.

### Oncostreams’ mesenchymal patterns reveal intra-tumoral collective motion in gliomas

GO analysis indicates that biological processes such as positive regulation of motility/migration are enriched within oncostream fascicles. Overexpression of extracellular matrix (ECM)-associated proteins suggest a potential role of COL1A1 fibers in regulating oncostreams’ motility. To study if oncostreams represent migratory structures within glioma tumors, we established a physiologically viable explant brain tumor slice model containing a high density of oncostreams (Fig. 7A). The movement of glioma cells expressing green fluorescent protein (GFP), within the thickness of each explant, was visualized using time-lapse confocal imagining and tracked using Fiji’s plug-in Track-Mate (Fig. 7A-C).

**Fig. 7.**
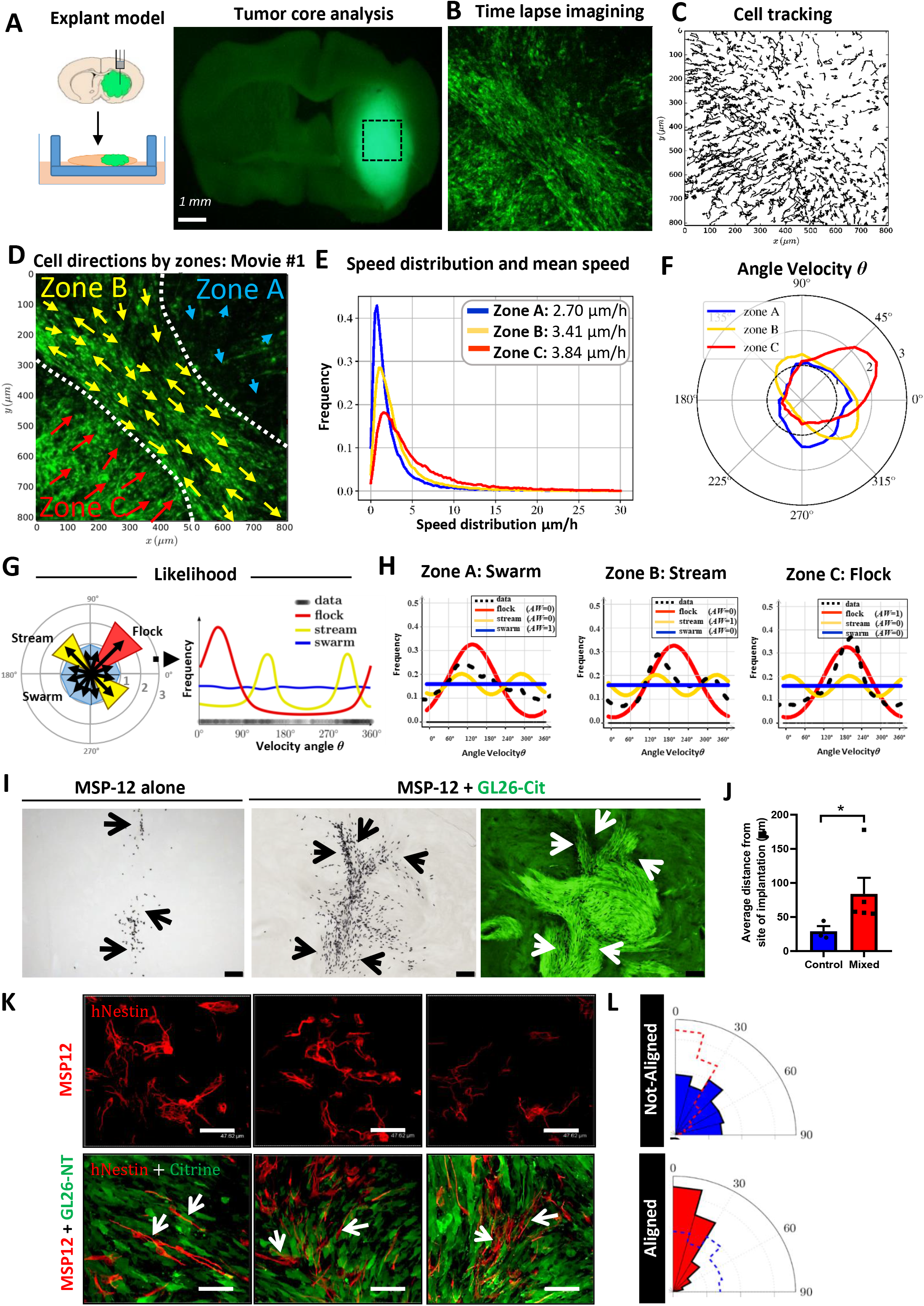
Collective dynamics of oncostreams increase cell spreading within the tumor core. **A)** Experimental setup: NPA-GFP glioma cells were intracranially implanted in C57BL6 mice. Explant slice cultures of growing tumors were used for confocal time-lapse imaging of the tumor core. **B)** Single representative time-lapse confocal image of glioma cells within the tumor core (Movie #1). **C)** Tracking analysis of individual cell paths performed using the Track-Mate plugin from Image-J. **D)** Preferred directions of cells within three zones (A-C) superimposed onto a representative time lapse-image. **E)** Speed distribution and mean speed (*µm*/*hr*) in Zones A, B and C. **F)** Distribution of angle velocity for each zone. The Angle Velocity of each cell is denoted *υ*. The plot shows the proportion of cells moving in angle direction *υ* for each zone. **G-H)** Classification of collective motion patterns: *stream*, *flock* or *swarm*. The distribution is *uni-modal* for a flock (only one peak) and *bi-model* for a stream (two peaks = 2 preferred angle velocity). For a *swarm*, the distribution is *flat* (no preferred angle velocity). In **(G)** Angle Velocity was transformed to a histogram; these data were then used to calculate the likelihood that a particular distribution of velocity angles corresponds to either a *stream*, *flock*, or *swarm*. The results are given in **(H)** for each zone. The frequency distribution of the data (shown in black) uses a *non-parametric* estimation (kernel density estimator). We tested three types of distributions, *ρ,* to describe the data-sets and give a likelihood for each case. The best fit was then determined by the Akaike weight (AW). **I)** Co-implantation of highly malignant GL26-citrine cells (green) and human MSP-12 glioma stem cells (ratio 1:30), and MSP-12 cells alone (control – left image). Immunohistochemistry of human nuclei (black) denote MSP-12 cells. Arrows show the distribution of MSP-12 cells within the brain or the tumor. Scale bar: 100 μm. **J)** Quantification of the distance of MSP-12 from the site of implantation. n=3 for control and n=5 for co-implantation (MSP-12+GL26). Error bars ± SEM; t-test, *p<0.05. **K)** Immunofluorescence images of human-nestin (red) labeling MSP-12 cells, and GL26-citrine cells. Note that MSP-12 cells have a multipolar morphological structure when alone, but a bipolar, elongated structure when aligned to GL26-citrine cells. Scale bar: 47.62 μm. **L)** Angle histogram plots quantify the alignment of MSP-12 within oncostreams, and the random alignment of MSP-12 cells when implanted alone (with dashed overlays of the other condition’s alignment).

Migration analyses show complex glioma cell dynamics throughout the tumor core. The glioma tumor core displays groups of cells (within particular zones) with similar nematic orientation and displaying complex movement patterns (Fig. 7D **and Fig. Supplementary 16A**) and, which represent collective motion^27, 29–31^. Angle velocity distribution indicated the existence of three patterns of collective motility shown schematically in Fig. 7D and F: in ‘Zone A’ cells don’t have a preferred direction, in ‘Zone B’ cells move in opposite directions (∼ 135° and 315°), and in ‘Zone C’ all cells move with a predominant preferred direction (∼ 45°) (Fig. 7D and F). We named these patterns ‘s*warm*’ (Zone A), ‘*stream*’ (Zone B), or ‘*flock*’ (Zone C) (Fig. 7G). They were classified by likelihood analysis: the distribution of the angle velocity is constant in a *swarm* (all angle velocity are equally probable), bi-modal in a *stream* (cells are moving in equal but opposite directions), and uni-modal in a *flock* (cells move in one direction) (Fig. 7H). These patterns were observed in all tumor slices examined (**Fig. Supplementary 18, 19 and 20**). Average cell speeds differed among the three patterns (Fig. 7E**, and Supplementary 18, 19 and 20**). In the tumor core, swarms moved faster and without orientation, followed by directionally moving flocks and streams (**Fig. Supplementary 27**). To determine which of these collective motion patterns match oncostream histological features, we analyzed H&E sections corresponding to imaged organotypic slices (**Fig. Supplementary 16B).** Cells within histological areas corresponding to *streams* and *flocks* have an aspect ratio of 2.2 and 2.7, respectively, (spindle-like cells), while those within areas corresponding to *swarms* have an aspect ratio of 1.2 (round cells) **(Fig. Supplementary 16C-D)**. Moreover, elongated cells within *streams* and *flocks* are nematically aligned with each other, whereas round cells within *swarms* are not **(Fig. Supplementary 16E).** As predicted by our *in silico* model, ^47^ these results suggest that cell shape, or eccentricity, is driving feature in the organization of collective motion patterns (**Fig. Supplementary 16F**). Therefore, taking into account cell shape and alignment, we define oncostreams as the histological expression of collective motion patterns (*streams* and *flocks*). Notice that only the dynamic analysis of collective motion can differentiate between *streams* and *flocks*. At the histological level both appear as oncostreams.

In collective motion of flocks, interactions among individual cells are sufficient to propagate order throughout a large population of starlings^48^. To define if oncostream migration patterns recall organized collective motion behavior, we analyzed the organization of the cells by performing local pair-wise correlation analysis (relative position and pair directional correlation) by tumor zones (**Fig. Supplementary 17A-C**). These analyses indicate the spatial correlation of location and alignment between individual cells. We observed that within *swarms* cells are more separated, as neighbors are located at 20-40 μm. S*treams* and *flocks* have higher cell density, and the nearest neighbors are closer, at 20-30 μm (**Fig. Supplementary 17E and S18, S19, S20**). Pair-wise directional correlation with nearby neighbors showed that cell movement is positively correlated in all patterns at distances between 10-50 *μm*, with higher correlation left-to-right for *streams* (≈0.2), left-to-right/front-to-back for *flocks* (≈0.2-0.4), and a lower correlation for *swarms* (≈0.1) (**Fig. Supplementary 17F and S18, S19, S20).** We ascertained that tumor cells within oncostreams migrate in a directional manner (“streams (↑↓)” and “flocks (↑↑)”), while non-oncostream cells move randomly without directional alignment as “swarms”. Thus, our analyses strongly indicate that within the tumor core of high-grade glioma cells are dynamically heterogeneous and display organized collective migratory behavior associated with tumor histological and genetic features.

### Oncostreams increase the intratumoral spread of tumoral and non-tumoral cells

Pair-wise correlation analysis showed that oncostream glioma cells are collectively organized. To test the underlying nature of collective oncostream motility, we analyzed adherent junction markers. Tumors with oncostreams were negative for E-cadherin, whereas N-cadherin was strongly expressed (**Fig. Supplementary 21A**), suggesting that these fascicles move in a manner akin to collective migration of mesenchymal cells of the neural crest^33, 49^. Although, no difference in N-cadherin were found within oncostreams and the surrounding areas, N-cadherin was elevated in TCGA-GBM (Grade IV) tumors compared to TCGA-LGG (Grade III and II). High levels of N-cadherin correlate with lower survival in HGG patients and mesenchymal transformation (**Fig. Supplementary 21B-C**).

On the other hand, oncostream growth and motility is unlikely to be due to glioma proliferation. BrdU staining showed no differences between oncostream and non-oncostream regions, and in the oncostreams, the mitotic plane was always perpendicular to the main axis as expected (**Fig. Supplementary 21D-E**). These results are also supported by the RNA-Seq data of dissected oncostreams, where proliferation genes were not differentially expressed (Fig. 4 A-C). Collective motion could affect the distribution of other cells within the tumor. Since oncostreams are heterogeneous, we inquired about their pro-tumoral role by potentially spreading cells throughout the tumor. We designed co-implantation experiments using human glioma stem cells (MSP-12), and highly aggressive and oncostream-forming glioma cells (GL26) co-implanted into immunosuppressed mice. Implantation of MSP-12 cells alone generated slow-growing tumors (median survival of 6-8 months). At 21 days post-implantation, MSP-12 cells remained restricted to the injection area with an average distance of 28.9±7.73 µm from the actual injection site. Surprisingly, when MSP-12 cells were co-implanted with GL26-citrine cells, MSP-12 cells spread throughout the tumor, moving along oncostreams to much longer distances (83.7±23.74 µm) from the injection site (Fig. 7I-K). Cellular cytoplasmic processes from MSP-12 cells implanted alone displayed a random distribution. However, in co-implanted tumors, such processes from MSP-12 cells are completely aligned with glioma GL26 cells within oncostreams (Fig. 7K-L **and Fig. Supplementary 21F)**. These results strongly suggest that oncostreams function as intra-tumoral highways facilitating the rapid distribution of slow-moving glioma cells and/or non-tumor cells throughout the tumor mass. These findings could help explain the dispersal and intratumoral mixing of diverse clonal populations as demonstrated in previous studies, supporting an important potential role of oncostreams in determining spatial cellular heterogeneity.

### Dynamic interactions at the tumor border: oncostreams foster glioma aggressiveness through collective invasion of the normal brain parenchyma

Furthermore, we asked whether oncostreams participate in glioma invasion. The analysis of histological sections showed that multicellular fascicles of elongated and aligned cells are found invading from the tumor border into the normal brain parenchyma (**Supplementary Fig. 22A**). Formation of streams around blood vessels was also observed (**Supplementary Fig. 22A**). These patterns of invasion are also detected using our deep learning methods (**Fig. Supplementary 22B**). We then used our glioma explant model to analyze the invasion dynamics by time-lapse confocal imaging at the tumor border (Fig. 8A **and S23**). We implanted glioma NPA GFP+ cells into tdTomato (mT/mG) mice so tumor borders could be delineated. We observed that glioma cells that extended from the tumor border to the normal brain parenchyma used different dynamic patterns, moving as isolated random cells and/or as collective migratory structures moving directionally, and resembled oncostream structures similar to those in the tumor core (Fig. 8B-G **and Fig. Supplementary 23-26**).

**Fig. 8.**
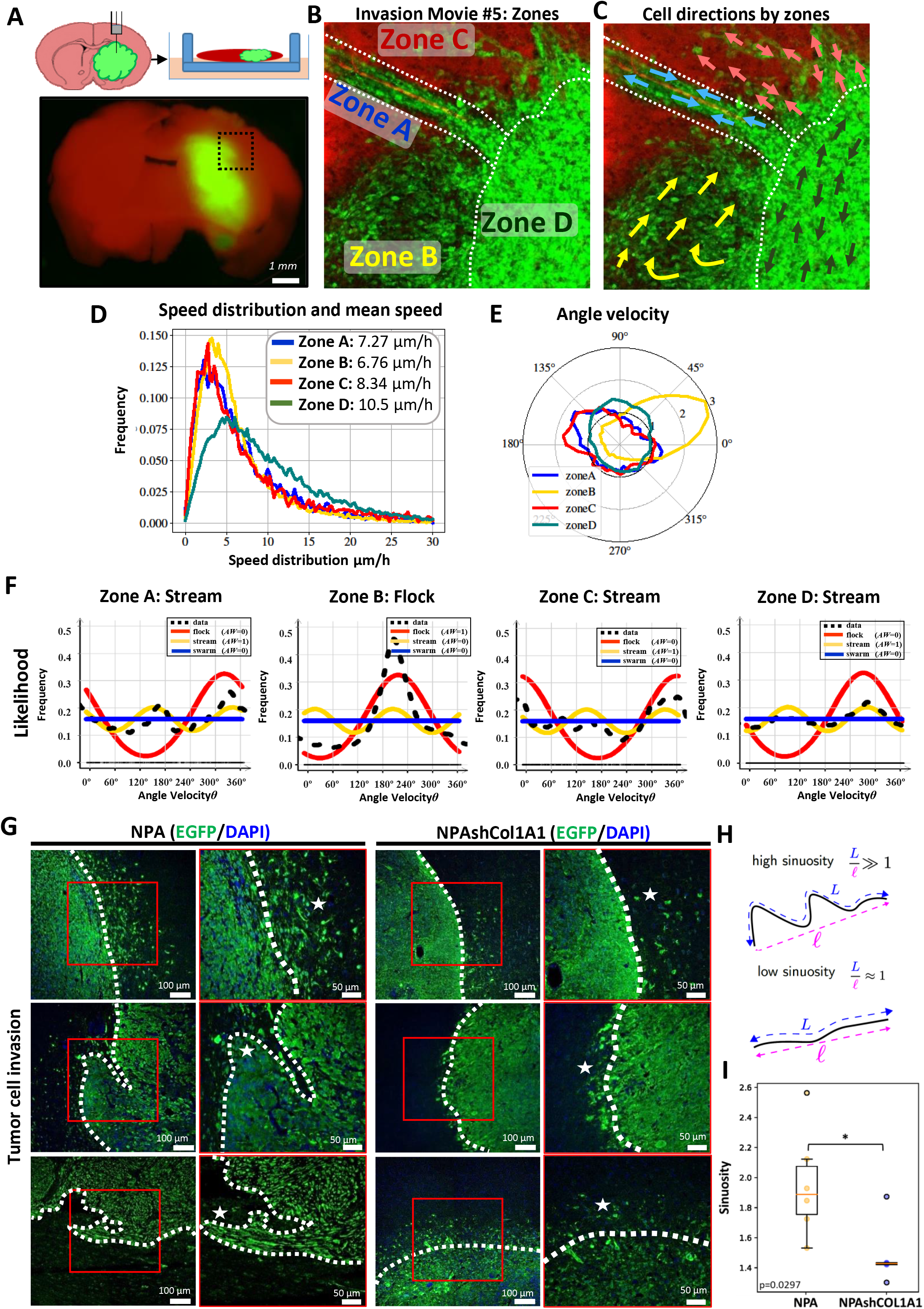
Collective invasion of COL1A1 enriched oncostreams contributes to malignant glioma behavior. **A)** Schematic representation of the experimental setup and location of imaging and quantification of tumor borders using td/mtTomato mice (Movie #5). **B)** Representative time-lapse scanning confocal image of glioma cells at the tumor border. This image was taken from border movie #1 and shows the subdivision into different zones. **C)** Preferred direction of cells within different zones superimposed onto a representative time lapse-image. **D)** Histogram of speed distribution and mean speed (*µm*/*h*) of Zones A, B, C and D. **E)** Angle Velocity distribution analysis (*υ*) performed by zones. Plot shows overall direction and magnitude of cell movement. **F)** Likelihood analysis of the dynamic patterns at the tumor border. Graph of density estimation *ρ flock* (red), *ρ stream* (yellow) and *ρ swarm* (blue). The estimation of the black line (data) uses a *non-parametric* estimation. AW: 0 or AW:1. **G)** Immunofluorescence analysis of GFP expression in GEMM of glioma controls (NPA), and NPAshCOL1A1. Representative confocal images of the tumor borders. GFP expression is shown in green (Alexa 488) and nuclei in blue (DAPI). Dotted lines show tumor borders. Stars show tumor cell invasion patterns. Notice the absence of collective invasion patterns in NPAshCOL1A1. Scale bar: 50 μm. **H**) The analysis of tumor borders was determined using the Allen-Cahn equation. Images were split into two values (−1 and +1) representing the inside and outside of the tumor to analyze the sinuosity of the borders. Illustration of the sinuosity of a curve: it is defined as the ratio between the length of the curve *L* and the distance between the two extreme points. The sinuosity is close to 1 for a straight line. **I)** Sinuosity of the border for all experiments. 4-10 images of each tumor border were obtained. NPA: n=6 and NPAshCOL1A1: n=5 tumors for each experimental condition were used for the analysis. We detected a decrease of the sinuosity in COL1A1 knockdown tumors. t-test unequal variance, *p=0.0297.

To objectively distinguish between different dynamic patterns, we determined the angle velocity distribution, and the likelihood that distributions corresponded to either a *stream*, a *flock*, or a *swarm*. We found *streams* along the perivascular niche or invading brain parenchyma without following any pre-existing brain structures, as well as cells invading as *flocks*, and *swarms* (Fig. 8E-F**, and Fig. Supplementary 23 D, G, H and S24-S26 A, C, D**). Glioma cells moving along blood vessels or directly into the brain as single cells is consistent with previous studies^10^. The correlation of position and pairwise correlation supports the existence of invading collective motion structures in NPA tumors with high expression of COL1A1 (**Fig. S22D-E and Fig. Supplementary S23J-K and-S24-S26 F-G**). We also determined the participation of collagen fibers in oncostreams invasion. Immunofluorescence analysis on explant slices showed that collagen fibers are aligned along multicellular fascicles of glioma cells invading the normal brain. These data show how collagen fibers serve as scaffolds for collective tumoral cell invasion (**Fig. Supplementary S27**).

Our data indicate the existence of a complex framework of collective motion patterns at the glioma border, that is consistent with previous descriptions^34^. Although the patterns observed at the NPA tumor border are similar to those of the tumor core, cell speed differed between the areas. Cells in the tumor core displayed significantly lower average speeds (*stream*: 4.26; *flock*: 5.95, *swarm*: 6.27 µm/hr) compared to cells at the tumor border or those invading the normal parenchyma (*stream*: 7.95; *flock*: 7.55, *swarm*: 8.01 µm/hr) (**Fig. Supplementary 28A-B**).

Then, we asked whether the knockdown of COL1A1 in NPA gliomas affects changes in the patterns of migration and invasion. Analysis of tumor cells (GFP+) at the tumor borders of GEMM of gliomas comparing NPA and NPA-shCOL1A1 showed a difference in the apparent invasion patterns. The analysis of tumor borders revealed an increase in the sinuosity of NPA tumors, a finding compatible with NPA tumors exhibiting a higher proportion of collective invasion into the normal brain when compared to NPA-shCOL1A1 tumors (Fig. 8G-I **and Supplementary Fig. S29**).

Moreover, the time lapse-confocal imaging and migration analysis of NPA-shCOL1A1 explants showed that tumor cells invade the normal brain parenchyma as isolated cells (**Supplementary Fig. 30-33**). Velocity angle, velocity vector and the likelihood analysis indicated that the overall distribution corresponded predominantly to *swarm* random patterns (**Supplementary Fig. 30-33 D, E, F)**. Further analysis of Relative Position Correlation and Pairwise correlation supports the presence of low density of cells compatible with single cell invasion patterns in NPAshCOL1A1 tumors with low expression of collagen (**Supplementary Fig. S30-33 G, H**).

We conclude that oncostreams (*streams* and *flocks*) are organized collective migratory structures enriched in COL1A1 that participate in the dynamic organization of the tumor microenvironment within the tumor core and at the tumor invasive border of high-grade gliomas, and facilitate invasion into the normal brain, impacting the malignant behavior of gliomas. Depletion of collagen1A1 eliminates oncostreams and their associated functions.

### Intravital imaging of glioma reveals the existence of oncostreams’ collective motion patterns *in vivo* and their contribution to invasion

To determine whether our previously described collective migration patterns of glioma cells *ex vivo* occur also *in vivo* we performed high resolution time lapse intravital imaging using two photon microscopy. To do so NPA glioma cells were intracranially implanted in the brains of tdTomato (mT/mG) mice at a depth of 0.8 mm (Fig 9A). To visualize cell migration we established a cranial window above the injection site (Fig 9B). After 7-15 days of tumor growth we acquired z-stack images to obtain a 3D orthogonal view of the tumor growing within the cortex. This allowed us to establish intravital imaging below the brain surface. Next, we selected a position at a depth of >100 µm and proceeded to acquire time lapse images of the tumor growing in the normal parenchyma at an interval of 5 minutes, for 8-12 hours (Fig 9C-D**, Supplementary Fig. S35A, S36A-B and S37A-B**).

**Fig. 9.**
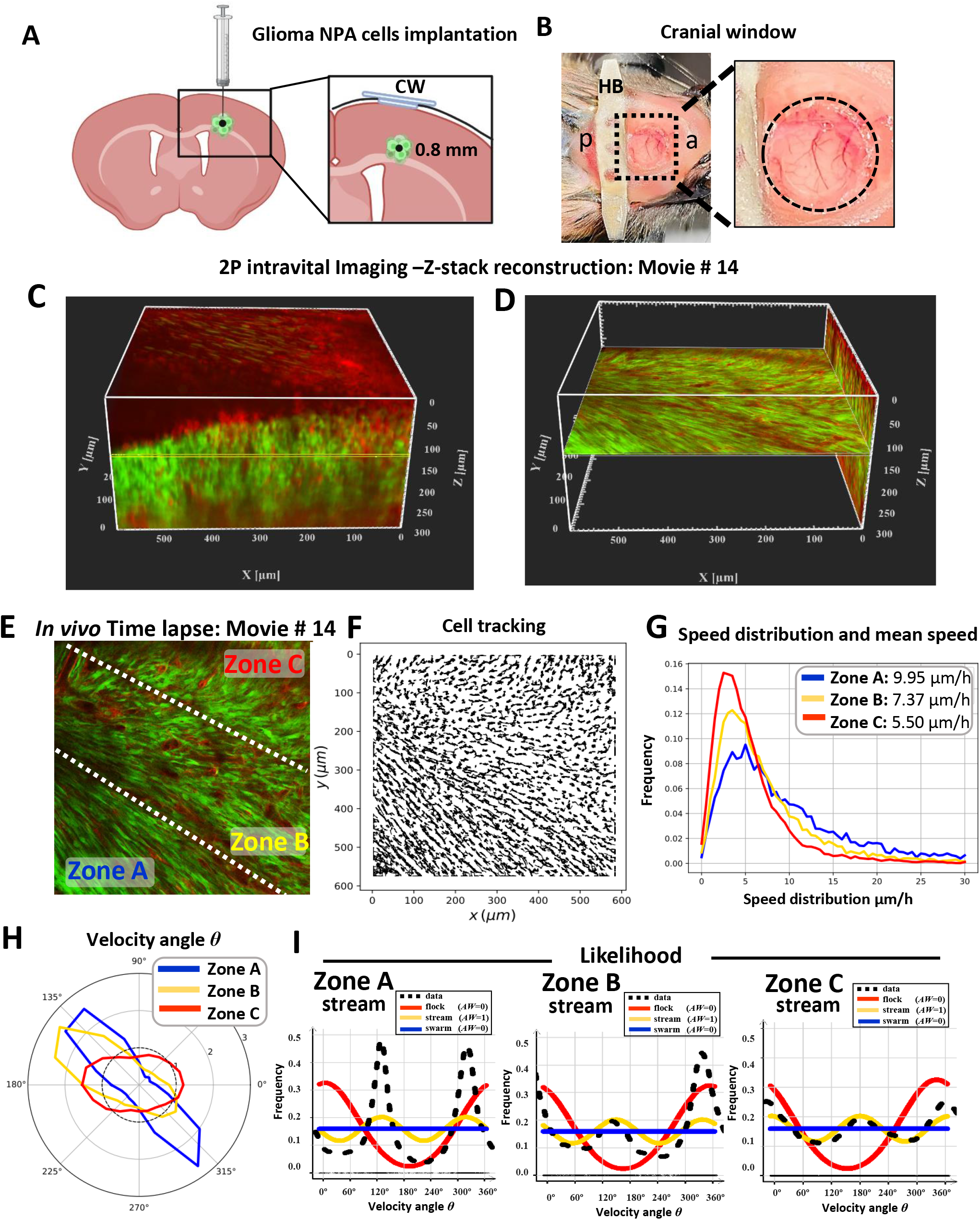
Intravital two-photon imaging reveals the collective patterns of glioma oncostream dynamics *in vivo.* **A)** Schematic representation of the cortical site of glioma cell implantation for intravital two-photon imaging. The inset shows a coronal section of the brain illustrating that glioma cells were implanted at 0.8 mm depth, with the glass cranial window (CW) positioned on top of the implantation site. **B)** Representative photograph of the head of an animal implanted with a cranial window showing the metallic head-bar (HB) positioned on the skull posterior to the cranial window. It affixed using dental cement to stabilize the imaging plane and minimize motion artifacts during time-lapse imaging. a: anterior; p: posterior. **C)** A high-resolution 3D z-stack spanning up to 300 μm depth (starting at the brain’s surface) was acquired on the multiphoton microscope, imported into the Imaris viewer and used to reconstruct this 3D image. 300 x-y frames from the brain’s surface were taken at a depth increment of 1 μm (voxel size=1) at a resolution of 1024×1024 pixels. XYZ axes of the 3D image are shown in white (596×596×300 μm), and the yellow line shows the exact imaging plane for time-lapse data acquisition *in vivo* (at 120 μm depth). Red fluorescent protein: normal brain parenchyma. Green fluorescent protein: tumor cells. **D)** This panel represents the X-Y and the Y-Z plane of the reconstructed 3D image (shown in C) using the Orthoslicer 3D function of the Imaris viewer software to illustrate the depth of the imaging plane. The X-Y plane shown at 120 μm depth illustrates the actual imaging position for movie #14 shown in **E. E)** Single representative time-lapse two-photon image of glioma cells within the tumor core *in vivo* (Movie #14) and imaged at a depth of 120 μm, showing Zones A, B, and C. **F)** Individual cell trajectories of the *in vivo* time-lapse experiment. **G)** Speed distribution and mean speed (μm/hr) for Zones A, B, and C, as indicated in **(E)**. **H**) Angle Velocity distribution for each zone’s in the *in vivo* time-lapse movie #14. The Angle Velocity of each cell is denoted θ. The plot shows the proportion of cells moving in the angle direction θ for each zone. **I)** Likelihood analysis of the dynamic patterns determined for each zone of Movie #14 obtained by intravital imaging. The frequency distribution *ρ flock* (red), *ρ stream* (yellow) and *ρ swarm* (blue) are shown. The estimation of the black line (data) uses a *non-parametric assessment* (kernel density estimator) to determine the structure of each zone. AW: 0 or AW:1.

In some cases, to determine the exact anatomical location of the tumor, following intravital imaging, we perfused-fixed the brain and performed fluorescence immuno-histochemistry analysis on paraffin embedded coronal sections which were imaged by confocal microscopy (**Supplementary Fig. S35B-C**). The location of the area imaged by intravital microscopy (>100 µm) is shown. In this figure we can identify the location of the tumor (GFP+ cells) containing parenchymal blood vessels (TdTomato+), and astrocytic processes (GFAP+) (**Supplementary Fig. S35B-C**). Thus, together with **Supplementary Fig. S35A**, we demonstrate imaging below the brain surface and within the normal parenchyma of the brain.

To analyze the movement of GFP+ glioma cells we used Fiji’s plug-in Track-Mate. In Fig. 9 the imaged area was divided into subregions to determine the existence of migration patterns (Fig. 9E,F). The analysis of cell migration *in vivo* showed that the glioma cells exhibit organized, nematically aligned cells moving collectively at a depth of 120 µm. Angle velocity distribution analysis determined the existence of ‘*stream*’ collective motion patterns in the three delimited subregions (for example in Fig. 9H), illustrating that aligned cells are moving in opposite directions. To corroborate the existence of ‘*stream’* patterns we applied our likelihood analysis. For ‘*streams’* the distribution of the angle velocity and velocity vectors displayed a bi-modal distribution (cells were moving in equal but opposite directions) (Fig. 9I), similar to that observed in the explant models. Mean speed varied from 5.50 to 9.95 µm/hour (Fig. 9 G). A different tumor imaged at a depth of 145 µm also showed the presence of ‘*streams’* (**Supplementary Fig. S36).**

To analyze the invasion of glioma cells, we imaged tumor movement at the border with normal brain (**Supplementary Fig. S35**). At an imaging depth of 140 µm, cells at the tumor border displayed ‘*stream’* collective dynamics (**Supplementary Fig. S35D, Zone A and B**). These motion patterns were determined using the angle velocity distribution analysis, and likelihood distribution analysis (**Supplementary Fig. S35 G-H**). Further analysis of Relative Position and Pairwise correlation supports the presence of high density of cells compatible with collective migration and invasion patterns (**Supplementary Fig. S34B-C, S35I-J and S36I-J**).

Further, to determine whether the knockdown of COl1A1 in NPA gliomas alters the migration patterns observed for NPA gliomas we analyzed a NPAshCOL1A1 tumor by two-photon microscopy at a depth of 110 µm (**Supplementary Fig. S37 A-B**).

The intravital migration analysis of NPAshCOL1A1 gliomas showed that glioma cells migrate without a preferred direction, and invade the normal brain parenchyma as ‘*swarms’* (**Supplementary Fig. S37 C-G**). The alignment analysis, Relative Position, and Pairwise correlation confirm the presence of not-aligned low-density cells compatible with single cell invasion patterns in gliomas with COL1A1 downregulation (**Supplementary Fig. S37 E, J, K).**

The collective motion patterns found *in vivo* resemble the collective motion patterns described in the *ex vivo* explant model. Our results show that glioma cells expressing collagen are organized in collective dynamic patterns at the tumor core and the tumor invasive border, in tumor explants and in *in vivo* intravital models of gliomas analyzed by two photon microscopy.

## DISCUSSION

Mesenchymal transformation is a hallmark of tumor heterogeneity that is associated with a more aggressive phenotype and therapeutic resistance^13, 18, 21^. Mesenchymal transformation involves fibroblast-like morphological changes associated with active migration and gain of expression of mesenchymal genes as previously described^21, 22^.

Herein we present a comprehensive study that defines the morphological, cellular, dynamic, and molecular properties of multicellular mesenchymal-like structures within gliomas. These structures are fascicles of aligned spindle-like cells found throughout the tumors and represent areas of mesenchymal transformation. We interpret these structures to be the histological expression of areas of collective motion of glioma cells. For the sake of simplicity, we have referred to these areas of mesenchymal transformation as oncostreams.

Oncostreams are areas of mesenchymal transformation and are identified histologically as fascicles of aligned and elongated cells. When examined dynamically, we found that tumor cells move by collective motion within the tumor core and at the invading border. The capacity to identify areas of collective motion in histological sections has allowed us to characterize the molecular organization of such dynamic structures. We thus describe the overall molecular mechanisms that govern the organization and function of these structures and demonstrate the causal role of individual mediators. Surprisingly, we discovered that COL1A1 is central to the structural and dynamic characteristics of oncostreams. Indeed, the loss of COL1A1 expression from tumor cells disrupts the structural and functional characteristics of oncostreams, resulting in a complete loss of mesenchymal areas within gliomas and a reduction in glioma malignant behavior (Fig. 10).

**Fig. 10.**
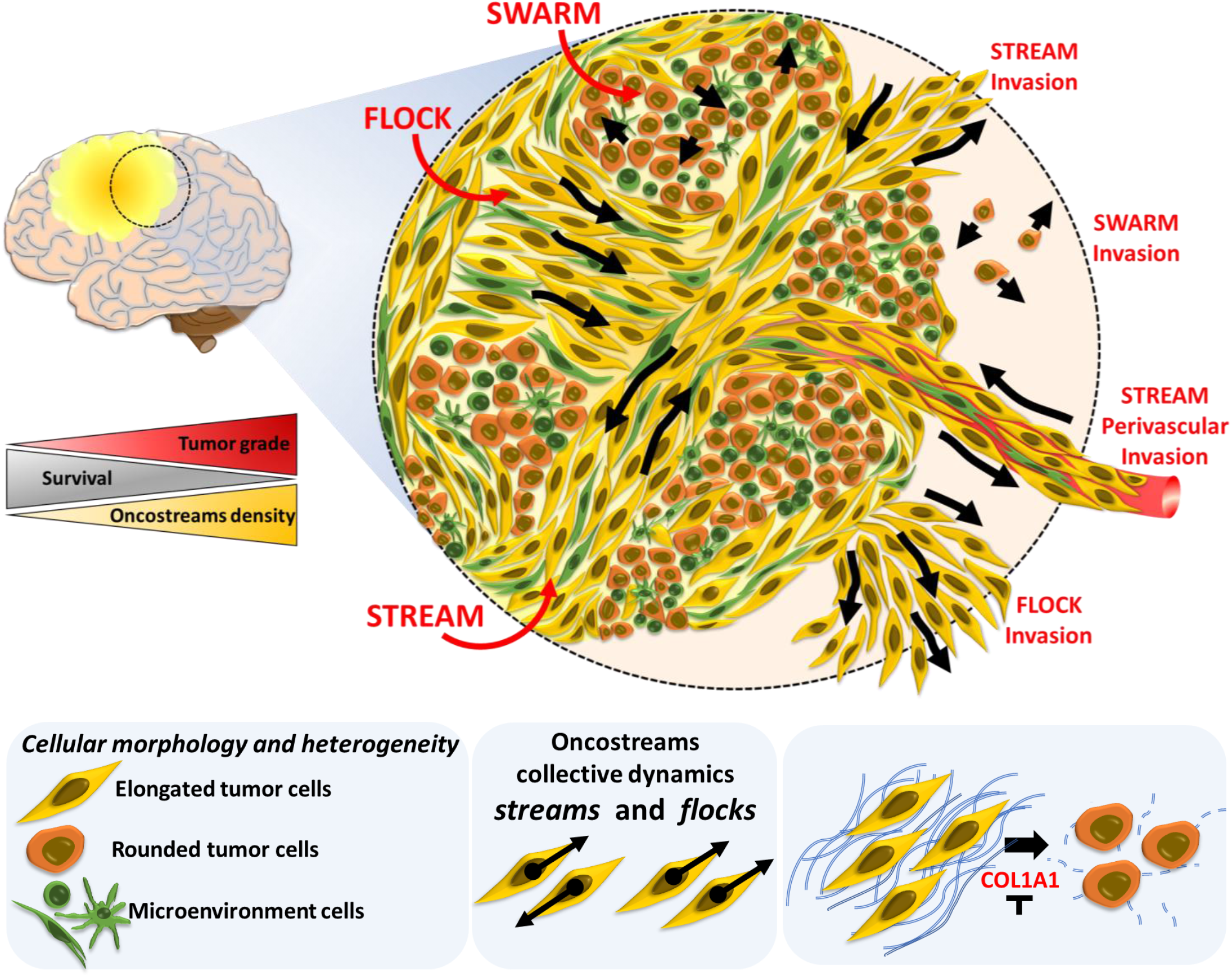
Oncostreams are COL1A1-rich multicellular dynamic mesenchymal structures that regulate glioma invasion and malignancy. Summary representation of mesenchymal dynamic fascicles (oncostreams) present in high grade gliomas. Our study reveals that oncostreams display directional collective motility patterns including streams and flocks. Non-directional collective motion (swarms) are represented by round cells that move do not have a preferred direction of motion. Directional dynamic patterns function as tumoral highways to facilitate the intra-tumoral spread of cells and participate in local invasion of normal brain. Oncostreams are areas of mesenchymal transformation defined by a molecular signature enriched in COL1A1. COL1A1 knockdown disrupts oncostream organization, decreases intratumoral heterogeneity and significantly increases animal survival. Our study reveals that oncostreams are anatomically and molecularly distinctive, are areas of mesenchymal transformation organized through interactions with the COL1A1 matrix, move by collective motion, and regulate glioma growth and invasion.

The analysis of the gene ontologies over-represented within oncostreams indicates that oncostreams denote areas enriched for “positive regulation of cell migration”, and in mesenchymal related genes. Interestingly, COL1A1 appeared as a central hub of oncostream organization and mesenchymal transformation. We postulate that oncostreams are the histopathological expression of patterns of collective motion (i.e., streams and flocks) in high grade glioma tumors. Different strategies of cell migration encountered in our gliomas are reminiscent of migratory characteristics observed during embryonic development^31, 32, 49^. In developmental biology, collective motion is represented by cells moving together in clusters, sheets, streams, or other multicellular arrangements^28, 31, 32^.

Our studies of oncostream dynamics at the tumor core are compatible with the results of Ralitza et al^50^. This group studied *ex-vivo* explant slices of spontaneous intestinal carcinoma, and showed that cells within the tumor core were highly dynamic and display directionally correlated cell motion^50^, similar to our results described herein. Recent *in silico* based mathematical modelling of glioma cell dynamics by our group, showed that only elongated cells, but not spherical cells, are able to form organized aligned cellular structures in a cell-density dependent manner^47^. Our modeling studies strongly support our *in-vivo* and *ex-vivo* data described in this manuscript.

Moreover, it has been described that increased matrix cross-linking, enzymatic remodeling and parallel orientation of matrix collagen fibers stiffens tissue, modifies cell morphology and promotes cell migration and invasion^36, 38, 39, 51, 52^. Our results support the proposal that oncostreams serve as highways to spread tumor, and non-tumor cells, throughout the tumor. Indeed, oncostream fascicles contain higher amounts of macrophages/microglia and mesenchymal cells. Dispersal of tumor and non-tumoral cells throughout the tumors could help explain the mixing of different clonal populations seen in molecular studies of high-grade gliomas^10^.

This study contributes to explaining how a particular feature of intratumoral heterogeneity, namely mesenchymal transformation, affects HGG progression. Our data indicate that the density of oncostreams plays a potential role in overall glioma malignant behavior in mouse and human gliomas.

Spatially resolved transcriptional analysis using laser capture microdissection provided novel insights into the molecular mechanisms that regulate oncostream functions. Oncostreams were defined by a unique transcriptomic signature that matched our immunohistochemical studies. COL1A1 overexpression within oncostreams was complemented with the overexpression of extracellular matrix proteins such as MMP9, MMP10, ADAMTS2, which are known to remodel and participate in the reorganization of collagen fibers. Oncostream fascicles were correspondingly enriched in COL1A1 when assessed by immunohistochemistry.

Within the extracellular matrix, collagen fibers constitute a scaffold for the organization of the tumor microenvironment and thus promote tumor infiltration and invasion. While collagen was previously thought to be a passive barrier that could reduce tumor invasion, it has now been shown that collagen fibers can serve as mechanical and biochemical tracks that facilitate cellular migration and tumor progression^36–38, 53^. Previously, multi-cancer computational analysis found that within a mesenchymal transformation signature in different cancers including gliomas, COL1A1 was one of the top differentially expressed genes^18, 22, 54^. COL1A1 is overexpressed in high grade malignant gliomas and its expression levels are inversely correlated with patient survival^55^ as indicated in https://www.cancer.gov/tcga. In our mouse glioma models and in human gliomas, tumors with higher density of oncostreams also express higher levels of COL1A1. COL1A1 is a consistently differentially expressed gene in the glioma mesenchymal signature identified in malignant gliomas and in glioma stem cells as described in previous studies^4, 9, 56^. Overall, our data are in agreement with a recent study by Puchalski *et al.*, which assigned genetic and transcriptional information to the most common morphological hallmarks of a glioma, emphasizing the importance of integrative histo-molecular studies^8^.

Surprisingly, our data indicate a remarkable plasticity of the mesenchymal phenotype in gliomas, similar to other studies^13, 15^. Genetic inhibition of COL1A1 within glioma cells depleted COL1A1 from tumors, eliminated oncostream structures, reduced the glioma malignant phenotype, and prolonged animal survival. Our findings are comparable with results from various studies that investigated the *in-vitro* and *in-vivo* consequences of collagen depletion, inhibition of collagen cross-linking or collagen synthesis inhibition on normalizing tumor ECM. In these studies, inhibition of collagen led to changes in the ECM which improved drug penetration, efficacy, as well as tumor access of therapeutic nano-particles or gene based therapies^57–61^. In addition, COL1A1 inhibition within glioma cells induced cell intrinsic and extrinsic changes in the TME. COL1A1 inhibition not only inhibits tumor cell proliferation and migration but also decreased the infiltration of microglia/macrophages, endothelial cells proliferation, and perivascular mesenchymal-like cells. As previously shown by other studies, glioblastomas exhibit complex interactions between tumoral and non-tumoral cells, (including macrophages, immune cells, and endothelial cells), which influence tumor growth, transformation, invasion, and response to treatment^9, 13, 62^. However, a major remodeling of the tumor mesenchymal phenotype in response to inhibition of COL1A1 has not been described earlier.

Moreover, we found that multicellular oncostream fascicles are detected in both *ex vivo* and *in vivo* glioma models, and that oncostreams facilitate tumor cell invasion, thereby increasing glioma aggressiveness. Our findings strongly support the importance of collective motility of glioma cells in the progression of tumor growth and invasion of normal brain parenchyma, and is supported by earlier studies of normal and pathological conditions^27, 34, 35, 63–67^.

In summary, our observations suggest that oncostreams are morphologically and molecularly distinct structures that represent areas of collective motion that contribute to tumor growth and invasion. These malignant dynamic structures overexpress COL1A1. COL1A1 knockdown eliminates oncostreams, reduces the mesenchymal phenotype, modifies the TME and slows tumor progression. Our findings open new vistas to understanding tumor mesenchymal transformation and its therapeutic treatment. We propose that depletion of COL1A1 within glioma cells is a promising approach to reprogram mesenchymal transformation in glioma tumors, and could be harnessed as a novel therapeutic approach, and reduce the glioma malignant phenotype.

## METHODS

### Glioma cell lines and culture conditions

Mouse glioma cells (NPA, NPD, NPAshCol1A1, NPDshCol1A1 and GL26) and human glioma cells (MSP-12, SJGBM2) were maintained at 37 °C with 5% CO2 and their respective media as described before^41–44^. Mouse NPA, NPD, NPAshCol1A1, NPDshCol1A1 neurospheres were derived from genetically engineered tumor using the Sleeping Beauty (SB) transposase system as previously described^41–44^. Mouse GL26 glioma cells were generated by Sugiura K and obtained from the frozen stock maintained by the National Cancer Institute (Bethesda, MD)^25^. MSP-12 human glioma cell lines were provided by Christine Brown, City of Hope, and SJGBM2 human glioma cells were provided by Children’s Oncology Group (COG) Repository, Health Science Center, Texas Tech University.

### Intracranial implantable syngeneic mouse gliomas

Glioma tumors were generated by stereotactic intracranial implantation into the mouse striatum of 3.0 x 10^4^ mouse glioma cells (either, NPA, NPD or, GL26) in C57BL/6 mice, or human glioma cells in immune-deficient NSG mice (SJGBM2) as described before^42–44, 68^. To test whether oncostream tumor cells help move other cells throughout the tumor we generated a co-implantation glioma model by intracranial implantation of highly malignant GL26-citrine cells with low aggressive human MSP12 glioma cells at a ratio of 1:30 (1,000 GL26-citrine cells and 30,000 MSP12 cells) in immune-deficient NSG mice. As controls, NSG mice were implanted with 30,000 MSP12 cells alone or 1,000 GL26-citrine cells alone as controls. Experiments were conducted according to the guidelines approved by the Institutional Animal Care (IACUC) and Use Committee at the University of Michigan. Stereotactic implantation was performed as previously described^42^.

### Genetically engineered mouse glioma models (GEMM)

We used genetically engineered mouse glioma models for survival analysis and histopathological analysis. Murine glioma tumors harboring different genetic drivers were generated using the Sleeping Beauty (SB) transposon system as described before^41–44^. Genetic modifications were induced in postnatal day 1 (P01) male and female wild-type C57BL/6 mice (Jackson Laboratory), according to IACUC regulations. shRNA targeting the COL1A1 gene was cloned as describe in detail in Supplementary Methods.

### Analysis of oncostreams in human glioma tissue

Oncostream presence was analyzed in unidentified H&E sections of paraformaldehyde-fixed paraffin-embedded (PFPE) human glioma samples obtained from primary surgery from the University of Michigan Medical School Hospital. To determine the presence of oncostreams in a large cohort of human glioma tissues we used the biospecimens from “The Cancer Genome Atlas Research Network” (TCGA) from the Genomic Data Commons Data Portal, National Cancer Institute, NIH (https://portal.gdc.cancer.gov). We analyzed primary Glioblastoma multiforme (TCGA-GBM) and Low-Grade Glioma (TCGA-LGG) databases. We selected cases that have available the Slide Image and diagnostic Slides. The diagnostic slides are available for TCGA-GBM: 389 patients and TCGA-LGG: 491 patients. The presence of oncostreams was scored on 100 TCGA-GBM Grade IV tissue samples and 120 TCGA-LGG samples.

### Cell aspect ratio and alignment analysis in H&E tumor sections

Images were obtained using bright-field microscopy of H&E stained paraffin sections (Olympus BX53 Upright Microscope). Tumors were imaged using 40X and 20X objectives. Images were processed using the program ImageJ as indicated in detail in supplementary methods.

### Deep learning analysis for oncostreams detection on H&E staining of glioma tissue

A fully convolutional neural network (fCNN) was trained in order to identify and segment oncostreams in histologic images^69^. We implemented a U-Net architecture to provide semantic segmentation of glioma specimens using deep learning^70–72^. Our oncostream dataset consisted of images from mouse tissues and open-source images from The Cancer Genome Atlas (TCGA). A total of 109 hematoxylin and eosin (H&E) stained histologic mouse images and 64 from TCGA were reviewed and oncostreams were manually segmented by the study authors (AC, A.E.A and P.R.L.). Images from both datasets were then augmented by randomly sampling regions within each image to generate unique patches (∼ 300 patches/image). The location and scale of each patch was randomly chosen to allow for oncostream segmentation to be scale invariant. The analysis is explained in further detail in Supplementary Methods.

### Immunohistochemistry on paraffin embedded brain tumors

This protocol was performed as described before^42^ and as is detailed in Supplementary Methods. Primary antibodies were diluted at the concentration indicated in Supplementary Table 8. Images were obtained using bright-field microscopy from five independent biological replicates (Olympus BX53 Upright Microscope). Ten different fields of each section were selected at random for study to include heterogeneous tumor areas. For immunofluorescence on paraffin embedded sections from brain tumors images were acquired with a laser scanning confocal microscope (LSM 880, Axio Observer, Zeiss, Germany). Integrated density was determined for the analysis of Col1a1 expression using Image J. For immunohistochemistry on vibratome brain tumor sections were left in 4% paraformaldehyde fixation for 48 hours and then transferred to PBS 0.1% sodium azide for an additional 24 hours at 4°C. A Leica VT100S vibratome was used to obtain 50 µm coronal brain sections. The immunohistochemistry protocol was performed as previously described^73, 74^.

### Laser capture microdissection (LCM) of brain tumors

Malignant glioma tumors were induced by intracranial implantation of dissociated NPA neurospheres in C57BL/6 mice as described above. LCM approach to analyze differential mRNA expression of intra-tumoral glioma heterogeneity was performed as described elsewhere^75^.

### RNA-Sequencing and bioinformatics analysis

RNA was isolated for laser microdissected tissues using the RNeasy Plus Micro Kit following the manufacturer recommendations (Qiagen). Before library preparation, RNA was assessed for quality using the TapeStation System (Agilent, Santa Clara, CA) using manufacturer’s recommended protocols. We obtained a RIN between 6 to 7 after laser microdissection of glioma tissue. A RIN of 6 was determined to be suitable for cDNA library preparation. 0.25 ng to 10 ng of total RNA was used for cDNA library preparation using a kit suitable for RNA isolation at pico-molar concentrations (MARTer Stranded Total RNA-Seq Kit v2-Pico Input Mammalian) following manufacturer recommended protocol (Clontech/Takara Bio #635005). Sequencing was performed by the UM DNA Sequencing Core, using the Illumina Hi-Seq platform and Bioinformatic analysis were executed by the UM Bioinformatics Core. Differentially expressed genes of all tumors were used for gene ontology (GO), Pathways analysis and genes analysis using iPathwayGuide (Advaita Corporation 2021). Network analysis of the DE genes were achieved using Cytoscape and Reactome App. Network was clustered by Reactome Functional Interaction (FI). Analysis of the expression of COL1A1 in normal tissue and in human gliomas were performed using the dataset of TCGA-GBM and TCGA-LGG from Gliovis (http://gliovis.bioinfo.cnio.es)^46^.

### Tumor explant glioma model and time-lapse confocal imaging

For the analysis of glioma dynamics, we generated tumors by intracranial implantation of 3 x 10^4^ NPA neurospheres which were used to carry out a 3D explant slice culture glioma model. C57BL6 mice were used for the dynamic analyses of the tumor core and B6.129(Cg)-Gt(ROSA)26Sortm4(ACTB-tdTomato,-EGFP)Luo/J-transgenic mice (Jackson laboratory, STOCK 007676) were used for invasion analysis. The red color correspond to a cell membrane-localized tdTomato (mT) fluorescence expression which is widespread in all cells and tissues. Mice were euthanized at 19 days’ post-implantation for NPA tumors and 31 days’ post-implantation for NPAshCOL1A1 tumors. Brains were removed, dissected, and embedded in a 4% agarose solution and kept on ice for 5 minutes. Then, brains were submerged in ice-cold and oxygenated media (DMEM High-Glucose without phenol red, GibcoTM, USA) and sectioned in a Leica VT100S vibratome (Leica, Buffalo Grove, IL) set to 300 μm in the z-direction. All steps were performed under sterile conditions in a BSL2 laminar flow hood. Brain tumor sections were transferred to laminin-coated Millicel Cell Culture Insert (PICM0RG50, Millipore Sigma, USA) (**Supplementary Fig. S23A**). Tumor slices were maintained in D-MEM F-12 media supplemented with 25% FBS, Penicillin-Streptomycin 10.000 U/ML at 37 °C with a 5% CO2 atmosphere. After 4-18 hours’ media were replaced with DMEM-F12 media supplemented with B27 2%, N2 1%, Normocin 0.2 %, Penicillin-Streptomycin 10.000 U/ML and growth factors EGF and FGF 40 ng/ml. For time-lapse imaging, slices were placed in an incubator chamber of a single photon microscope. We utilized an inverted Zeiss LSM880 laser scanning confocal microscope with AiryScan (Carl Zeiss, Jena, Germany), equipped with an incubation chamber kept at 37 °C with a 5% CO2. To validate the depth of imaging, in some experiments (see **Supplementary Fig. S23A-C**), before time lapse imaging, we obtained high resolution z-stacks with approximate dimensions of z=143.81, x=850.19, y=850.19. Time-lapse images were then obtained every ten minutes for 100-300 cycles, selecting the imaging plane to fall at the middle of the z-stack to avoid imaging at the bottom of the explant. Following movie acquisition, sections were fixed in 4% paraformaldehyde (PFA) for 2 days. Fixed sections were embedded in 2% agarose for H&E and immunohistochemistry analysis. Sections were processed and embedded in paraffin at the University of Michigan Microscopy & Image Analysis Core Facility using a Leica ASP 300 paraffin tissue processor/Tissue-Tek paraffin tissue embedding station (Leica, Buffalo Grove IL). Tumor explants were used for collagenase treatment. Sections were then treated for 48 hours with collagenase (C2399, MilliporeSigma, USA) at a concentration of 5, 10, or 15 units/ml or vehicle control. Following treatment, sections were fixed in 4% paraformaldehyde (PFA) for 2 days.

### Cranial window implantation and two photon intravital live imaging *in vivo*

Craniotomy and cranial window implantations were performed following previously described protocols by us and others^25, 76, 77^. The protocol was conducted according to the guidelines approved by the Institutional Animal Care and Use Committee (IACUC) at the University of Michigan. Briefly, mice were anesthetized and placed in a stereotactic frame. A craniotomy of 3×3 mm size was made over the right hemisphere between bregma and lambda. 5×10^4^ GFP^+^ NPA glioma cells were intracranially implanted at a depth of 0.8mm ventral, near the center of the craniotomy overlying the brain cortex of B6.129(Cg)-Gt(ROSA)26Sortm4(ACTB-tdTomato-EGFP)Luo/J mice. These mice were utilized to identify (in red) normal brain tissue, and thus establish tumor borders. The cranial window was covered with two round microscope cover glasses, and a metal head bar was positioned on the skull posterior to the cranial window. One week post NPA tumor cells’ injection and cranial window implantation, and two-weeks post NPAshCOL1A1 implantation, intravital live imaging was performed using a two-photon microscope (Bruker Technology) with a 20X water immersion objective (Olympus, NA 1.0) for 8-12 hours. To avoid imaging at the brain surface, first we acquired high-resolution 3D z-stacks spanning 0-330 µm depth from the surface of the brain. Z-stacks were then imported to Imaris viewer version 9.8 (Bitplane, Imaris, Oxford Instruments, MA, USA) to obtain 3D images. We then used Orthoslicer3D to reveal the depth of the imaging position in relation to the surface of the brain for time-lapse data acquisition. The detailed methodology is available in Supplementary Methods.

### Mathematical analysis of tumor cell movement

To determine the movement of cells in different areas of the tumor we performed localized statistical analysis in different zones of the tumor. We selected localized areas based on the organization of cells in clusters, group of cells moving together with similar distribution. Raw data of 4 movies from the tumor core and 4 movies from the tumor border were analyzed for 293 cycles (core) and 186 cycles (border) for a frame rate of Δ t = 10 min between image acquisition. To track cell motion, we used the software Fiji with the plugin TrackMate^78^. Analysis was performed as indicated in detail in Supplementary Methods.

### Classification of glioma migration patterns

To classify the collective cellular motion behavior of the three types of patterns called flock, stream, and swarm illustrated in Supplementary Figure S10F we used as criteria the orientation of each cell described by its unique angle velocity denoted θi. More precisely, we transformed the Angle Velocity Distribution graph into a histogram where we examined the distribution of all the values θi. A schematic representation of these distributions is depicted in Figure 4G. Considering a data-set θn n=1…N of orientations where N is the total number of cells, θn ɛ [0, 2π] is the direction of the cell n. We tested three types of distributions ρ to describe the dataset and gave a likelihood in each case as described in Supplementary Methods. The Akaike Weight (AW) indicates which pattern has the highest likelihood in each experimental situation^79^.

### Statistical Analysis

All in vivo experiments were performed using independent biological replicates, as indicated in the text and figures for each experiment. Data are shown as the mean ± SEM. Any difference was considered statistically significant when p < 0.05. In experiments that included one variable, the one-way ANOVA test was used. In experiments with two independent variables, the two-way ANOVA test was employed. A posterior Tukey’s multiple comparisons test was used for mean comparisons. Student’s t-test was used to compare unpaired data from two samples. Survival data were entered into Kaplan-Meier survival curves plots, and statistical analysis was performed using the Mantel log-rank test. Median survival is expressed as MS. Significance was determined if p<0.05. All analyses were conducted using GraphPad Prism (version 8.0.0) or SAS (2021 SAS Institute, Cary, NC). Each statistical test used is indicated within the figure legends.

## Data availability

All data associated with this study are in the paper and/or the Supplementary Information. RNA-Seq data was deposited at the NCBI’s Gene Expression Omnibus (GEO) with identifier GSE188970. Source data are provided with this paper. Further information and requests for resources and reagents should be directed to and will be fulfilled by the corresponding author P.R. Lowenstein.

## Code Availability

The analysis of oncostreams in mouse and human glioma tissue was performed using U-Net architecture to provide semantic segmentation of specimens using deep learning. Public GitHub repository for the project code can be found at https://github.com/MLNeurosurg/DeepStreams.

Analysis of glioma cells dynamics was performed using the Julia Programing Language. Link for this project Script and their dependencies can be found at public GitHub repository https://github.com/smotsch/analysis_glioma.

## ACKNOWLEDGMENTS

We thank all members of our laboratory for advice and comments on this work. This work was supported by National Institutes of Health, National Institute of Neurological Disorders and Stroke (NIH/NINDS) grants: R37-NS094804, R01-NS105556, R21-NS107894, R21-NS091555; R01-NS074387 to M.G.C.; National Institute of Neurological Disorders and Stroke (NIH/NINDS) grants: R01-NS076991, R01-NS096756, R01-NS082311, R01-NS122234, R01-NS127378 to P.R.L.; National Institute of Biomedical Imaging and Bioengineering (NIH/NIBI): R01-EB022563; National Cancer Institute (NIH/NCI) U01CA224160; Rogel Cancer Center at The University of Michigan G023089 to M.G.C. Ian’s Friends Foundation grant G024230, Leah’s Happy Hearts Foundation grant G013908, Pediatric Brain Tumor Foundation grant G023387 and ChadTough Foundation grant G023419 to P.R.L. RNA Biomedicine grant: F046166 to M.G.C. Health and Human Services, National Institutes of Health, UL1 TR002240 to Michigan Institute for Clinical and Health Research (MICHR), Postdoctoral Translational Scholars Program (PTSP), Project F049768 to A.C.

## AUTHORS INFORMATION

### Contributions

Conception and design: A. Comba, M.G. Castro, P. R. Lowenstein. Development of methodology: A. Comba, M.S. Faisal, P. J. Dunn, A. E. Argento, T. Hollon, W.N. Al-Holou, M.L. Varela, D.B. Zamler, S. Motsch, P. R. Lowenstein. Acquisition of data, analysis, and interpretation: A. Comba, M.S. Faisal, P. J. Dunn, A. E. Argento, T. Hollon, M.L. Varela, D.B. Zamler, Clifford Abel II, M.G. Castro, S. Motsch, P. R. Lowenstein. Human histopathology analysis and identification of oncostreams: A. Comba, C. Kleer, A. E. Argento, P. R Lowenstein. Laser microdissection protocol: A. Comba, P. R. Lowenstein, P. E. Kish, Alon Kahana. Development and establishment of intravital imaging using multiphoton microscopy: Comba, M.S. Faisal, P. R. Lowenstein, G. L. Quass, P. F. Apostolides. Development and experimental assistance with human glioma cell lines. development and assistance: C.E. Brown. Manuscript writing: A. Comba, S. Motsch, P. R. Lowenstein. Administrative, technical, or material support (i.e., reporting or organizing data, constructing databases): A. Comba, P. R. Lowenstein. Study supervision: M. G. Castro and P. R. Lowenstein. All authors reviewed the final version of the manuscript.

### Corresponding author

Correspondence to Pedro R. Lowenstein

## ETHICS DECLARATIONS

**Competing interests:** All authors of this paper declare no potential conflicts of interest.

## SUPPLEMENTARY INFORMATION

**Supplementary Material and Methods**

**Supplementary Figures**

**Supplementary Tables**

**Supplementary Videos**

## Notes

### Competing Interest Statement

The authors have declared no competing interest.

### Summary of Updates

The new data was added to the revised version of the manuscript. We included new data for changes in the tumor microenvironment, intravital imaging of glioma migration, and invasion and analysis of glioma invasion using intravital imaging in WT and shCOL1A1 inhibition models of glioma, and analysis of tumor borders invasion.

