## Supplementary Info and Figures for "Spatiotemporal analysis of glioma heterogeneity reveals Col1A1 as an actionable target to disrupt tumor mesenchymal differentiation, invasion and malignancy"

**This file includes:**

Supplementary Materials and Methods

Supplementary Figures S1-S37

Supplementary Tables S1, S3, S5, S6, S7, S8, S9

**Other Supplementary Materials:**

Movies #1 to #17

Table S2

Table S4

#### SUPPLEMENTARY METHODS

##### Genetically engineered mouse glioma models (GEMM)

To design and clone the shRNA targeting the Col1a1 gene (pT2-shCol1a1-GFP4), we tested 3 22 base pair sequences for the mouse Col1a1 gene selected from candidate sequences within the RNAi codex database (http://cancan.cshl.edu/cgi-bin/Codex/Codex.cgi) and InvivoGene (http://www.invivogen.com/sirnawizard). shCol1a1-A: CCCTGGTGATACTGGTGTTAAA, shCol1a1-B: GCAACAGTCGCTTCACCTACA, shCol1a1-C: GCAAGACAGTCATCGAATACA and shRNA Scramble: GAATCTAATCGTATCGTGGCTG. To confirm Col1a1 knockdown, NIH/3T3 mouse cells were transfected with the designed shRNA plasmids for 72 hours and subsequently analyzed by Western Blot. Col1a1 protein levels were normalized to β-actin levels and compared to the Scramble control. Genetic models generated include the following gene expression/inhibitions: (i) shp53, NRAS-G12V and shATRX **(NPA)**, (ii) shp53, NRAS-G12V, shATRX and IDH1-R132H **(NPAI),** (iii) shp53, NRAS-G12V and PDGFβ **(NPD),** (iv) shp53, NRAS-G12V, shATRX, shCol1a1 **(NPAshCOL1A1),** (v) shp53, NRAS-G12V, PDGFβ, shCol1a1 **(NPDshCOL1A1).** Plasmid sequences were verified by Sanger sequencing. Plasmid sequences described below were used to generate tumors: (i) pT2C-LucPGK-SB100X, transposon & luciferase expression; (ii) pT2-NRAS-G12V, RTK/RAS/PI3K activation; (iii) pT2-shp53-GFP4, p53 knock-down; (iv) pT2-shATRx-GFP4, ATRX knock-down; (v) pKT-IDH1(R132H)-IRES-Katushka, mIDH1 expression; (vi) pT2-shp53-PDGFβ-GFP4, p53 knock-down in combination with PDGFβ ligand overexpression. The pT2CAG-NRAS-G12V and pT2-shp53-GFP4 plasmids were the generous gift of Dr. John Ohlfest (University of Minnesota). Mice were injected with the necessary plasmids according to the protocol described (*1, 2*). Plasmid uptake in pups and tumor growth was monitored by IVIS® Spectrum imaging. Adult mice displaying symptoms of morbidity were transcardially perfused (*1, 2*).

##### Analysis of oncostreams in human glioma tissue

Oncostream presence was analyzed on tumors classified by histology grade (grade II, III and IV) as shown in **Table S1.** GBM were also analyzed by molecular subtypes: Classical (CL), Mesenchymal (MES), and Proneural (PN). LGG were classified in relation to grade and subtypes which consider IDH status and 1p/19q co-deletion (IDHmut-codel, IDHmut-non-codel and IDHwt)

(**Table S2**). H&E histology samples were analyzed at high magnification using the Slide Image Viewer from the data portal. Histological material containing brain tumors and oncostreams of both rodent and human gliomas were evaluated by CGK (board certified pathologist), AC and PRL. Concordance regarding the presence or absence of oncostreams, between the three evaluators, was >90%. Due to the need for good quality morphological preservation, only paraformaldehyde-fixed paraffin-embedded (PFPE) sections were used. Clinical data including age, sex, pathology, survival, treatment information, and data including information of MGMT DNA methylation status, IDH1 mutation status, G-CIMP DNA methylation status, were obtained from http://firebrowse.org, http://gliovis.bioinfo.cnio.es and https://portal.gdc.cancer.gov dat a portals (**Table S2**).

**Cell aspect ratio and alignment analysis in H&E tumor sections**

Image processing included color deconvolution to isolate nuclei, transforming images to 8-bit, and adjusting the threshold to remove background. Elliptical overlay masks were then imposed over the nuclei to match their shape. The shapes of these masks were then analyzed for the shape descriptors aspect ratio and circularity. For cell alignment analysis images were processed using ImageJ. Briefly, image processing included creating a ROI around the stream area and the non-stream area. These ROIs were then saved as separate images. Both images underwent color deconvolution to isolate nuclear stain, 8-bit conversion, and threshold adjustment to remove background. Images were then analyzed for the Feret's angle of the nucleus. The Feret's angle is the angle ($0^{o}$-$180^{o}$) that is taken from the x-axis to a line parallel with the longest distance across the particle being considered. Histograms were generated with these data in MATLAB using the Matplotlib plugin.

**Immunohistochemistry on paraffin embedded brain tumors (IHC-DAB)**

Following perfusion with Tyrode saline solution, mouse brains were fixed in 4% paraformaldehyde (PFA), and post-fixed for an additional 48 hours at 4 °C. Brains were then processed and embedded in paraffin at the University of Michigan Microscopy & Image Analysis Core Facility using a Leica ASP 300 paraffin tissue processor/Tissue-Tek paraffin tissue embedding station (Leica, Buffalo Grove IL). Tissue was sectioned using a rotary microtome (Leica) set to 5 µm in the z-direction. Endogenous Peroxidase quenching was carried out using a 0.3% $H_2O_2$ incubation for 5 minutes at room temperature. Heat-induced antigen retrieval was performed using 10mM citric acid, 0.05% Tween 20, pH 6.0. Tissue permeabilization and blocking were completed using PBS 0.2% Tween-20 with 5% goat serum for one hour at room temperature. Tissue was incubated with primary antibodies at $4^o$ C overnight at concertations detailed in **Table S8.** Tissue sections were then incubated with secondary biotinylated antibodies at a 1:1000 dilution in PBS with 0.2% Tween-20 overnight at $4^o$ C. ABC Avidin-Biotin-Complex Binding reagent (Vectastain Elite ABC kit) and Betazoid DAB Chromogen detection kit (BioCare BDB2004) were used according to the manufacturer's instructions.

**Immunofluorescence on paraffin embedded sections from brain tumors**

This protocol was performed as described before (*2*). Paraformaldehyde fixed and paraffin-embedded tissues were sectioned and then de-paraffinized and re-hydrated. Heat-induced antigen retrieval was performed using 10mM Citric Acid, 0.05% Tween 20, pH 6.0. Permeabilization was performed using 0.5% TritonX-100 in PBS for 30 minutes at room temperature while shaking. Tissue sections were blocked in 10% horse serum and 3% BSA in PBS. Primary antibodies were incubated at 4˚C overnight in a humid chamber in 3% BSA in PBS at concertation detailed in **Table S8.** Tissues were then incubated for 90 minutes at room temperature with Alexa Fluor™ 488 or Alexa Fluor™ 555 conjugated secondary antibodies (Invitrogen, Thermofisher Scientific). Col1a1 detection was performed using HRP-conjugated streptavidin was incubated for 60 minutes at room temperature. Alexa Fluor™ 488 Tyramide was incubated for 10 minutes. Nuclei were stained with DAPI (1:1000) in PBS for 5 minutes.

**Immunohistochemistry on vibratome brain tumor sections**

Vibratome sections were permeabilized in TBS-Triton-X 0.1% for 60 minutes. Antigen retrieval was performed in 10mM sodium citrate. Non-specific antibody binding was blocked with 10% goat serum in TBS-Triton-X 0.1% for 1 hour at room temperature. Brain sections were then incubated with primary antibody diluted in TBS-Triton-X 0.1%, 1% goat serum, and 0.1% sodium azide for 24hours at RT, in the dark. Sections were then washed 6 times in TBS-Triton-X 0.1% and then incubated with the secondary antibody diluted in 1% goat serum in TBS-Triton-X 0.1% for 24hrs at RT, in the dark. Finally, sections were washed 6 times and incubated with 5μg/ml of 4',6-diamidino-2-phenylindole (DAPI) (Life technologies, D21490) in PBS for 5 minutes. Sections were washed again 3 times and mounted on microscope slides with prolong gold anti-fade reagent (Invitrogen, P36930).

**Deep learning analysis for oncostreams detection on H&E staining of glioma tissue**

First, using the mice dataset only, six iterations of training/validation set splits (80%/20%) were generated by randomly sampling unique, non-overlapping regions from each of labelled histologic images and used for model selection/hyper-parameter tuning and model testing, respectively. Our fCNN was trained for binary classification of foreground (oncostream) and background (non-oncostream) using a binary cross-entropy loss function. Both pixel-level classification accuracy and intersection over union (IOU) were used as metrics to evaluate model performance. IOU values over the testing set there were manually annotated. Our model generalized well to both mouse and human data. In addition to having a mean IOU value of 80.3%, our model did not make any false positive errors in TCGA grade II gliomas (**Supplementary Table S9).** The model was trained using the Adam optimizer with an initial learn rate $\alpha = 0.0001$, $\beta1 = 0.9$, $\beta2 = 0.999$, and $\varepsilon = 10E\text{-}7$ (*3*). We used a scheduled learning rate decrease such that the rate was halved every 10 epochs and the model was trained for a total of 75 epochs for each iteration. The best performing model on the mouse dataset was then used as a pre-trained fCNN to initialize training on the TCGA data. We used both mouse and TCGA images for fCNN training with 20% of the TCGA data held out for model validation. Our fCNN was implemented using the model-level Python-based API, Keras (version 2.2.0), with a TensorFlow (*4*) (version 1.8.0) backend running on two NVIDIA GeForce 1080 Ti graphical processing units. This model is generalizable to both mouse and human images.

**Laser capture microdissection (LCM) of brain tumors**

This methodology conserves tissue histology integrity and RNA quality. Briefly, animals were perfused with Tyrode solution (0.8% NaCl, 0.0264% $CaCl_2$, 0.005% $NaH_2PO_4$, 0.1% glucose, 0.1%

$NaHCO_3$, 0.02% KCl) for 5 minutes followed by 30% Sucrose dissolved in Tyrode solution for 15 minutes. Tissue was maintained overnight at 4$^{o}$C in 30% sucrose solution prepared in RNAse free water. Brain tumors were cryopreserved in OCT and stored at -80$^{o}$C. Tissue was cryosectioned at 10 µm thickness, placed onto 2 µm polyethylene naphthalate (PEN) and stored at -80$^{o}$. Tumor sections were fixed in ethanol and stained with 4% Cresyl violet and 0.5% eosin dyes dissolved in ethanol. Oncostream areas from 3 independent mouse glioma samples were dissected from surrounding glioma tissue using a LMD7000 Leica laser capture microdissection (LCM). Tissue was collected in DNase/RNase free 0.5 mL PCR flat head tubes containing 30 µL of RLT lysis buffer and stored at -80$^{o}$ until RNA purification.

**RNA isolation, quality control and cDNA library preparation**

A total tissue area of 2.5 x $10^6$ to 7 x $10^6$ $\mu m^2$ was dissected for each group to increase the efficiency of the mRNA extraction and cDNA library preparation. Sample volume was adjusted to 350 µL with lysis buffer with 1% β-mercaptoethanol. DNA was removed using gDNA eliminator spin columns. RNA was eluted in 12 µL of RNase-free water warmed at 37 °C. Final libraries were checked for quality and quantity by bioanalyzer on a Eukaryote total RNA Pico chip using manufacturer's recommended instructions. The samples were pooled, sequenced on the Illumina HiSeq 4000, as paired-end 50 nt reads with 10% PhiX spike-in, according to manufacturer's recommended protocols. Tuxedo Suite software package was used for alignment, differential expression analysis, and post-analysis diagnostics by the University of Michigan Bioinformatics Core.

**RNA-Sequencing and bioinformatics analysis**

Quality of the raw reads data for each sample using FastQC (version v0.11.3) to identify features of the data that may indicate quality problems (e.g. low quality scores, over-represented sequences, inappropriate GC content). We used the Tuxedo Suite software package for alignment, differential expression analysis, and post-analysis diagnostics. Reads were aligned to the reference genome including both mRNAs and Ensemble lncRNAs (UCSC mm10) using TopHat (version 2.0.13) and Bowtie2 (version 2.2.1). Default parameter settings were used for alignment, with the exception of: “b2-very-sensitive” telling the software to spend extra time searching for valid alignments. FastQC was used for a second round of quality control (post-alignment), to ensure that only high quality data would be input to expression quantitation and differential expression analysis. Cufflinks/CuffDiff (version 2.1.1) technique was implemented for differential expression analysis, using UCSC mm10.fa as the reference genome sequence. For expression quantitation, normalization, and differential expression analysis, “--multi-read-correct” was used to adjust expression calculations for reads that map in more than one locus, as well as “compatible-hits-norm” and “upper-quartile–norm” for normalization of expression values. Diagnostic plots were generated using the CummeRbund R package.

Genes and transcripts have been identified as differentially expressed based on three criteria: test status = “OK”, FDR ≤ 0.05, and fold change ≥ ± 1.5. Genes and isoforms were annotated with NCBI Entrez GeneIDs and text descriptions. Volcano plots demonstrating total genes and DE genes were produced with the R base package.

**Collagenase treatment of tumor explant brain slices model**

Glioblastoma tumors were generated by intracranial implantation of 3.0 x $10^4$ NPA neurosphere cells into the striatum of C57BL/6 mouse brains. At 19 days following implantation, mice were euthanized per ULAM guidelines. Brains underwent explant sectioning as described previously. Following sectioning, tissue sections were maintained on laminin coated Millicel Cell Culture for 24 hours ate described above. Sections were then treated for 48 hours with collagenase (C2399, Millipore Sigma, USA) at a concentration of 5, 10, or 15 units/ml or vehicle control. Following treatment, sections were fixed in 4% paraformaldehyde (PFA) for 2 days. Fixed sections were embedded in 2% agarose. Sections were then processed and embedded in paraffin at the University of Michigan Microscopy & Image Analysis Core Facility using a Leica ASP 300 paraffin tissue processor/Tissue-Tek paraffin tissue embedding station (Leica, Buffalo Grove IL).

**Craniotomy and cranial window implantation**

Mice were anesthetized using an intraperitoneal injection of injectable anesthetics ketamine and dexmedetomidine. One dose of carprofen was given pre-operatively, and another dose was given

24hr post-surgery. Eye ointment was applied prior to surgery, and hairs overlying the skull were clipped with a hair-clipper. Following anesthesia induction, the mouse was placed in a stereotactic apparatus and betadine disinfectant was applied to the incision site. The skin was cut, retracted and periosteum, as well as musculature on the surface of the skull, was gently scraped to ensure entire cranium dry and rough. Then, a craniotomy was made over the right hemisphere in between the bregma and lambda (size, 3x3 mm; center relative to Bregma: lateral right, 0.5mm; posterior, 0.5mm). Multiple rounds of slow and delicate drilling were performed to ensure that underlying brain matter did not overheat. The skull was moistened with chilled sterile saline solution in between rounds of drilling to further reduce heat and swelling. Once the edges of the craniotomy are thin enough, the skull fragment in a single piece was then removed using fine forceps or a 26G needle. Intracranial injection of 5x$10^4$ $GFP^+$ NPA or NPAshCol1A1 glioma cells were implanted at a depth of 0.8mm ventral through the craniotomy overlying the brain cortex of B6.129(Cg)-Gt(ROSA)26Sortm4(ACTB-tdTomato-EGFP)Luo/J using a Hamilton syringe in a volume of 1ul over the period of 5min. These tdTomato mice were utilized to identify (in red) normal brain tissue, and thus establish tumor borders.

Following the injection, imaging window was constructed from two layers of standard microscope cover glass (Electron Microscopy Sciences), of which one was small (3mm) equal to the size of the craniotomy and another was large (5mm), which were joined with an ultraviolet curable optical adhesive (NOR-71, Norland). Two layered coverslip was carefully placed over the exposed brain filled with saline to facilitate visual access to tumor and brain during the imaging. A larger piece was attached to the cranium and a smaller insert was fitted snugly into the craniotomy. The cyanoacrylate-based adhesive (Krazy glue) was spread with a needle towards the edges of the larger coverslip to avoid leeching of the glue underneath of the coverslip obscuring the brain. To permanently secure the coverslip on the skull, the coverslip was held in place by gentle pressure from a blunt needle of the syringe. A thin layer of glue was then applied over the entire surface of the exposed skull except for the location of the craniotomy. Once the adhesive was set, the blunt needle was slowly lifted away, and the coverslip was further reinforced to the skull with dental cement. A lightweight, metallic headbar was positioned on the skull posterior to the cranial window and affixed using dental cement to stabilize the imaging plane and minimize motion artifacts during the time-lapse imaging. A dental acrylic well was created around the cranial window to immerse the imaging objective. Upon hardening of the dental cement, the animal was removed from and allowed to recover on a heating pad in its home cage while under constant supervision. The animal was placed in a new home cage with only one animal per cage and allowed to recover. The animals were recovered from anesthesia using Atipamezole via intra-peritoneal injection (1.0 mg/Kg) to reverse Ketamine/Xylazine, and a single subcutaneous injection of Buprenorphine (0.1mg/kg subcutaneous) was administered as post-operative pain relief. Water-soaked chow in ceramic crock was placed on the cage floor post-op. The following day the animal was received a second dose of carprofen for pain relief.

**Two photon intravital live imaging *in vivo***:

One-week NPA tumor cells' injection and cranial window implantation, and two-weeks post NPAshCOL1A1 implantation, mice were monitored for tumor growth using an IVIS bioluminescence imaging system. Mice with positive bioluminescent signal were placed in the glass jar and anesthesia was induced using isoflurane (3%). After anesthesia was achieved, mouse was transported to the multiphoton imaging platform and held in place using the headbar and headbar clamp. The breadboard (Thorlabs) was fitted with a nose cone for applying inhalational isoflurane anesthesia while the mouse was placed in the headbar clamp. Water base immersion oil was used to fill the dental acrylic well and the objective of the microscope was lowered towards the cranial window until it was situated just beneath the surface layer of oil. The animal was placed on a heating pad to maintain physiologic body temperature regulated by a rectal thermal probe (Stoelting), under continuous light anesthesia with isoflurane (1%). Dehydration was prevented by giving 300µl isotonic saline every three hours using a catheter placed subcutaneously. Eyes were lubricated with eye-ointment to prevent drying while under anesthesia. The vitals of the mice were monitored using a mouse thigh sensor with MouseOX Plus (Star Life Sciences Corp). To visualize glioma dynamics in real-time in the living animal with a Bruker Optima two-photon microscope, using a Chameleon Ti-sapphire laser (Coherent) tuned to 920 nm and attenuated with a Pockels cell to adjust the power at the sample so that approximately the same signal level could be achieved at a constant frame rate starting from

150mW for the surface to 300mW at imaging depths of 300-330 µm. The images were acquired with a 20X water immersion objective (Olympus, NA 1.0). To avoid imaging at the brain surface, first we acquired high-resolution 3D z-stacks spanning 0-330 µm depth from the surface of the brain. Z-stacks were then imported to Imaris viewer version 9.8 (Bitplane, Imaris, Oxford Instruments, MA, USA) to obtain 3D images. We then used Orthoslicer3D to reveal the depth of the imaging position in relation to the surface of the brain for time-lapse data acquisition. Images within the cortex were taken of the GFP+ tumor (peak excitation: 700nm) and TdTomato brain (peak excitation: 920nm) every 5min for up to 8-12 hours. Emitted fluorescence was split with a dichroic mirror to separate green (GFP) and red (Tdtomato) channels. Once imaging was complete, animals were removed from the stereotactic frame and allowed to recover in a clean cage under a heating lamp.

**Mathematical analysis of tumor cell movement**

We used as parameters for the cell size (called 'blob') 20µm and a threshold of 1 together with the DoG method (Difference of Gaussian detectors). We then obtained for each experiment several paths for many different cells. To reduce the erratic behavior, we filtered the trajectories. From the data obtained with TrackMate, we then smoothed out the paths which allowed to estimate the velocity of each cell. We simply use a Gaussian Kernel as a filter (with standard deviation $\sigma = 2$ and a stencil of 9 points). Thus, we had an estimation of the positions $x_i(t)$ and velocities $v_i(t)$ of the cells i. From the velocity $v_i(t)$, we also deduced the velocity direction $\vartheta_i(t)$.
After obtaining these parameters we applied several statistics to investigate cell behavior. Each cell is characterized by a position $x_i \in R2$ and a velocity $v_i \in R2$. The velocity vector v can be decomposed into a speed (scalar) $c = |v|$ and angle direction $\vartheta$ such that $v = c (\cos \vartheta, \sin \vartheta)$. We utilized three statistics to visualize the distribution of velocity: (i) distribution of velocity angle $\vartheta$ (provides indication of the overall direction of the cells); (ii) distribution of speed c and (iii) distribution of the velocity vector $v = (vx, vy)$. Since v is a vector, we need to plot a 'heatmap' to visualize its distribution.
To understand the organization of oncostream dynamics we analyzed the cell-cell correlation using two further statistics using pair wise information: (i) Neighbor density: taking a reference cell xi, we estimate the probability to have another cell $x_j$ nearby, and, (ii) velocity correlation:

depending on the position of a nearby $x_j$, estimate the correlation: $\omega_i \cdot \omega_j$ where $\omega_i$ (resp. $\omega_j$) is the (normalized) velocity direction of cell *i* (resp. $_j$). A correlation of +1/-1 indicates that cells are moving in the same/opposite direction, whereas 0 indicates that they move in orthogonal direction. Orthogonal direction is when $\omega_i \cdot \omega_j = 0$. Details on how to estimate numerically the correlation functions are described in **Fig. S17 A, B, C**.

**Classification of glioma migration patterns**

Our data-set consists of cell orientations $\theta_n \in [0, 2\pi)$ with $n = 1..N$ indexing the cells and $N$ being the total number of cells in a given zone. Zones were defined by either varying density, alignment, geographical distribution, and/or movement patterns of cells for each experiment. We build a criteria to decide whether the underlying distribution of orientations $\theta$ is *more like* a flock, a stream or a swarm. With this aim, we define three types of distributions $\rho_{flock}$, $\rho_{stream}$ and $\rho_{swarm}$ and use *likelihood* estimation to determine which distribution is more likely (given the data-set $\{\theta_n\}_{n=1..N}$). First, we present the three distributions used to describe a flock, a stream and a swarm.

**Flock: wrapped normal distribution**. In a flock, the distribution of orientations should be *uni-modal*, i.e. the orientations are centered around a given direction denoted $\overline{\theta}$. Thus, we use for $\rho_{flock}$ a *wrapped* Gaussian distribution:

$$\rho_{flock}(\theta;\ \overline{\theta}, \sigma) = \frac{1}{\sigma\sqrt{2\pi}} \sum_{k=-\infty}^{\infty} {}^{-\frac{|\theta-\overline{\theta}+2\pi k|^2}{2\sigma^2}}, \qquad \text{(Eq. 1)}$$

where $\overline{\theta}$ represents the 'peak' of the distribution and $\sigma$ measures its spreading. Summing in *k* is necessary to ensure that the distribution $\rho_{flock}$ is $2\pi$ periodic in $\theta$. The two free parameters $\overline{\theta}$ and $\sigma$ of the distribution are determined by maximizing the *likelihood*, i.e. by determining $\overline{\theta}_*$ and $\sigma_*$ the maximizers of the following quantity:

$$\overline{\theta}_*, \sigma_* = \text{argmax}_{\overline{\theta},\sigma} \mathcal{L}(\overline{\theta}, \sigma) \quad \text{with} \quad \mathcal{L}(\overline{\theta}, \sigma) = \prod_{n=1}^{N} \rho_{flock}(\theta_n;\ \overline{\theta}, \sigma). \qquad \text{(Eq. 2)}$$

After some standard computations (taking the log), we find an explicit solution:

$$\overline{\theta}_* = \mathrm{Arg}(\overline{\mathbf{z}}) \quad , \quad \sigma_*^2 = -\ln R_e^2$$

with $\overline{\mathbf{z}} = \frac{1}{N}\sum_{n=1}^{N} {}^{i\theta_n}$ (using complex number) and $R_e^2 = \frac{N}{N-1}\left(|\overline{\mathbf{z}}|^2 - \frac{1}{N}\right)$.

**Stream: symmetric wrapped normal distribution**. In a stream, the distribution of orientations is only $\pi$ periodic since the distribution of cells moving in both directions inside the stream should be alike. Thus, we use for $\rho_{stream}$ a symmetric version of $\rho_{flock}$:

$$\rho_{stream}(\theta;\ \overline{\theta}, \sigma) = \frac{1}{2}\Big(\rho_{flock}(\theta;\ \overline{\theta}, \sigma) + \rho_{flock}(\theta + \pi;\ \overline{\theta}, \sigma)\Big) \qquad \text{(Eq. 3)}$$

where $\rho_{flock}(\theta;\ \overline{\theta}, \sigma)$ is defined in (Eq. 1) . The estimation of the two free parameters $\overline{\theta}$ and $\sigma$ can be estimated as previously by maximizing the likelihood. We notice for that the following identity:

$$\begin{aligned} \rho_{stream}(\theta;\ \overline{\theta}, \sigma) &= \frac{1}{2\sigma\sqrt{2\pi}} \sum_{k=-\infty}^{\infty} (^{-\frac{|\theta-\overline{\theta}+2\pi k|^2}{2\sigma^2}} + ^{-\frac{|\theta-\overline{\theta}+2\pi k+2\pi|^2}{2\sigma^2}}) \\ &= \frac{1}{2\sigma\sqrt{2\pi}} \sum_{k=-\infty}^{\infty} (^{-\frac{|2\theta-2\overline{\theta}+4\pi k|^2}{2(2\sigma)^2}} + ^{-\frac{|2\theta-2\overline{\theta}+4\pi k+2\pi|^2}{2(2\sigma)^2}}) \\ &= \frac{1}{2\sigma\sqrt{2\pi}} \sum_{k=-\infty}^{\infty} {}^{-\frac{|2\theta-2\overline{\theta}+2\pi k|^2}{2(2\sigma)^2}} = \rho_{flock}(2\theta;\ 2\overline{\theta}, 2\sigma). \end{aligned} \qquad \text{(Eq. 4)}$$

Thus, the parameters $\overline{\theta}_*$ and $\sigma_*$ for the stream distribution are estimated using with $\overline{\mathbf{z}}_\mathbf{2} = \frac{1}{N}\sum_{n=1}^{N} {}^{i2\theta_n}$ in lieu of $\overline{\mathbf{z}}$.

**Swarm: uniform distribution.** In a swarm, the orientations of the cells are 'random' meaning that all directions are equally probable. Thus, the distribution $\rho_{swarm}$ is simply constant:

$$\rho_{swarm}(\theta) = \frac{1}{2\pi}.$$

There is no free parameters. The normalization $2\pi$ is to make sure that the distribution satisfies $\int_0^{2\pi} \rho_{swarm}(\theta)\,\mathrm{d}\theta = 1$.

**Model selection.** Since we have now three distributions $\rho_{flock}$, $\rho_{stream}$ and $\rho_{swarm}$, we can determine the likelihood of our data-set $\{\theta_n\}_{n=1..N}$ in each case. The higher the likelihood, the higher the chance the model (flock, stream or flock) is selected. However, this strategy will never select a swarm since there is no free parameter in $\rho_{swarm}$ and a flock or stream distribution can be made uniform with $\sigma$ large. One could penalize having free parameters in $\rho_{flock}$ and $\rho_{stream}$ using information criterion (e.g. AIC, BIC) but even then swarm distributions are not selected, a flock or a stream will always be selected.

For this reason, we *fix* a priori the value of $\sigma$ for the flock and stream distribution $\rho_{flock}$ and $\rho_{stream}$. In other words, a distribution will be considered a flock if the distribution is not "too" flat. We choose:

$$\sigma^2_{flock} = 2 \quad , \quad \sigma^2_{stream} = 1. \qquad \text{(Eq. 5)}$$

We give an illustration with the experiment "scene 5" zone D in figure 3. The associated log-likelihood gives:

$$\ln(\mathcal{L}_{flock}) = -46798.93, \;\; \ln(\mathcal{L}_{stream}) = -43583.16, \;\; \ln(\mathcal{L}_{swarm}) = -43732.28. \qquad \text{(Eq. 6)}$$

Thus, in this example, we select a stream formation since the log-likelihood is the highest.

#### Analysis of the tumor borders

**Phase separation.** We denote $h(x,y)$ the original image of green intensity where $(x,y)$ is a position (pixel position) and $h(x,y) \in [0,1]$ the intensity of green color (i.e. 1 indicate a tumor, 0 no tumor). We want to separate the image into two regions: inside the tumor ("+1") and outside ("$-1$").

To do so, we define a threshold $h_* = .12$ and then apply the Allen-Cahn equation to the function $c_0(x,y) = h(x,y) - h_*$:

$$\partial_t c = c(1 - c^2) + \gamma \Delta c \tag{1}$$

where $\gamma = .5$ is the diffusion parameter to *smooth* the distribution. The source term ``$c(1 - c^2)$'' makes the function $c(x, y, t)$ converge to either $-1$ or $+1$. The dynamics [1] can be interpreted as a gradient flow with respect to the energy:

$$\mathcal{E}[c] = \int_{\mathbb{R}^2} \frac{1}{4}(c^2 - 1)^2 + \frac{\gamma}{2}|\nabla c|^2 \, \mathrm{d}\mathbf{x}. \tag{2}$$

We present in Supplementary Figure S28 the evolution of the $c(x, y, t)$ over time, transitioning from a representative original image (in red/blue scale instead of green) into a *binary* image with only two values $-1$ and $+1$.

**Statistic of the border.** Once the image has been split into two regions (using Allen-Cahn equation), we investigate the sinuosity of the border. The sinuosity is defined as the ratio between the length of the curve *L* and the distance joining the end points of the curve (see Figure 8H-I). The more ’sinuous’ a curve is, the large the sinuosity. We create a boxplot computing the sinuosity for all the experiments regrouping the experiments with/without collagen.

### Supplementary Tables

**Table S1. Analysis of oncostreams on TCGA glioma diagnostic slides from the Genomic Data Commons Portal**

| TCGA Manual analysis | | | | |
|---|---|---|---|---|
| **Grade** | **Recurrence** | **Total Tumors** | **OS Positive** | **OS Negative** |
| **GBM-Grade IV** | Primary | 100 | **47** | 53 |
| **LGG-Grade III** | Primary | 70 | **6** | 64 |
| **LGG-Grade II** | Primary | 50 | **0** | 50 |

**Table S3. Summary results of oncostreams detection using manually histopathological analysis and deep learning segmentation**

| Accuracy of methodologies for Oncostreams analysis | | | | | |
|---|---|---|---|---|---|
| Tumor | Method | Oncostreams | | Concordance | |
| | | Positive | Negative | Images # | Percentage (%) |
| GBM - Grade IV | Manually | 78 | 31 | 92/109 | 84.4 |
| | Deep Learning | 78 | 31 | | |
| LGG - Grade III | Manually | 29 | 97 | 113/126 | 89.7 |
| | Deep Learning | 24 | 102 | | |
| LGG - Grade II | Manually | 0 | 61 | 60/61 | 98.4 |
| | Deep Learning | 1 | 60 | | |

**Table S5. Enriched Gene Ontologies (GOs) biological process within oncostream fascicles**

| GO ID | Go Name | countDE | countAll | pv_fdr |
|---|---|---|---|---|
| GO:0030335 | positive regulation of cell migration | 9 | 561 | 0.000845 |
| GO:2000147 | positive regulation of cell motility | 9 | 584 | 0.000845 |
| GO:0040017 | positive regulation of locomotion | 9 | 600 | 0.000845 |
| GO:0051272 | positive regulation of cellular component movement | 9 | 604 | 0.000845 |
| GO:0030334 | regulation of cell migration | 10 | 932 | 0.002304 |
| GO:2000145 | regulation of cell motility | 10 | 982 | 0.002731 |
| GO:0040012 | regulation of locomotion | 10 | 1029 | 0.003686 |
| GO:0051270 | regulation of cellular component movement | 10 | 1072 | 0.004748 |
| GO:0006950 | response to stress | 18 | 3567 | 0.006971 |
| GO:0032963 | collagen metabolic process | 4 | 111 | 0.007461 |
| GO:0016477 | cell migration | 11 | 1488 | 0.011264 |
| GO:0040011 | locomotion | 12 | 1842 | 0.01536 |
| GO:0048870 | cell motility | 11 | 1647 | 0.022187 |
| GO:0051674 | localization of cell | 11 | 1647 | 0.022187 |
| GO:0051240 | positive regulation of multicellular organismal process | 12 | 1973 | 0.022674 |
| GO:0030198 | extracellular matrix organization | 5 | 301 | 0.022674 |
| GO:0043062 | extracellular structure organization | 5 | 302 | 0.022674 |
| GO:0009605 | response to external stimulus | 14 | 2673 | 0.027927 |
| GO:0006952 | defense response | 10 | 1521 | 0.040737 |

**Table S6. Pathways of the Network for COL1A1**

| Pathways of the Network including Col1a1 | p-value | FDR | Genes | minusLOGFDR |
|---|---|---|---|---|
| Relaxin signaling pathway(K) | 2.96E-07 | 2.52E-05 | MAPK3,JUN,FOS,MMP9,NFKB1,COL1A1,ACTA2 | 4.598823558 |
| AGE-RAGE signaling pathway in diabetic complications(K) | 2.31E-05 | 3.99E-04 | MAPK3,JUN,STAT3,NFKB1,COL1A1 | 3.398907391 |
| Focal adhesion(K) | 5.91E-04 | 3.55E-03 | FYN,MAPK3,JUN,COL1A1,CTNNB1 | 2.450138812 |
| Human papillomavirus infection(K) | 1.01E-03 | 5.07E-03 | HDAC2,MAPK3,ISG15,NFKB1,COL1A1,CTNNB1 | 2.29537768 |
| Extracellular matrix organization(R) | 2.00E-03 | 7.99E-03 | ADAMTS2,MMP9,MMP10,COL1A1,TLL1 | 2.097529324 |
| RNA Polymerase II Transcription(R) | 2.69E-03 | 9.98E-03 | HDAC2,IRAK1,UBC,MAPK3,JUN,FOS,COL1A1,SP1,CTNNB1 | 2.00075198 |
| VEGFR3 signaling in lymphatic endothelium(N) | 3.43E-03 | 0.0103 | MAPK3,COL1A1 | 1.987162775 |
| Platelet activation(K) | 8.79E-03 | 0.0213 | FYN,MAPK3,COL1A1 | 1.671620397 |
| Beta3 integrin cell surface interactions(N) | 9.77E-03 | 0.0213 | CYR61,COL1A1 | 1.671620397 |
| Integrin signalling pathway(P) | 0.0172 | 0.0343 | FYN,MAPK3,COL1A1 | 1.46470588 |
| IL4-mediated signaling events(N) | 0.0207 | 0.0414 | COL1A1,SP1 | 1.382999659 |
| Amoebiasis(K) | 0.0434 | 0.0472 | NFKB1,COL1A1 | 1.326058001 |

**Table S7. Pathways of the Modules**

| Module | GeneSet | FDR | Nodes | NegLOGFDR |
|---|---|---|---|---|
| M1 | Alzheimer disease-presenilin pathway(P) | 0.0461 | ACTA2,MMP9 | 1.336299075 |
| | Leukocyte transendothelial migration(K) | 0.0461 | CDH5,MMP9 | 1.336299075 |
| | Relaxin signaling pathway(K) | 0.0461 | ACTA2,MMP9 | 1.336299075 |
| | Fluid shear stress and atherosclerosis(K) | 0.0461 | CDH5,MMP9 | 1.336299075 |
| | Effects of Botulinum toxin(N) | 0.0461 | SYT1 | 1.336299075 |
| | Synaptic_vesicle_trafficking(P) | 0.0461 | SYT1 | 1.336299075 |
| | Extracellular matrix organization(R) | 0.0461 | MMP9,MMP10 | 1.336299075 |
| | Syndecan-1-mediated signaling events(N) | 0.0461 | MMP9 | 1.336299075 |
| | Wnt signaling pathway(P) | 0.0461 | ACTA2,CDH5 | 1.336299075 |
| | Nicotinic acetylcholine receptor signaling pathway(P) | 0.0461 | ACTA2 | 1.336299075 |
| | S1P2 pathway(N) | 0.0461 | CDH5 | 1.336299075 |
| | Osteopontin-mediated events(N) | 0.0461 | MMP9 | 1.336299075 |
| | amb2 Integrin signaling(N) | 0.0461 | MMP9 | 1.336299075 |
| | Syndecan-4-mediated signaling events(N) | 0.0461 | MMP9 | 1.336299075 |
| | Validated transcriptional targets of AP1 family members Fra1 and Fra2(N) | 0.0461 | MMP9 | 1.336299075 |

| Module | GeneSet | FDR | Nodes | NegLOGFDR |
|---|---|---|---|---|
| M2 | Beta5 beta6 beta7 and beta8 integrin cell surface interactions(N) | 0.0405 | CYR61 | 1.392544977 |
| | Beta2 integrin cell surface interactions(N) | 0.0405 | CYR61 | 1.392544977 |
| | Primary immunodeficiency(K) | 0.0405 | CIITA | 1.392544977 |
| | Beta3 integrin cell surface interactions(N) | 0.0405 | CYR61 | 1.392544977 |
| | RhoA signaling pathway(N) | 0.0405 | CYR61 | 1.392544977 |
| | Antimicrobial peptides(R) | 0.0405 | LCN2 | 1.392544977 |
| | Iron uptake and transport(R) | 0.0405 | LCN2 | 1.392544977 |
| | AP-1 transcription factor network(N) | 0.0405 | CYR61 | 1.392544977 |
| | Interferon gamma signaling(R) | 0.0405 | CIITA | 1.392544977 |
| | Antigen processing and presentation(K) | 0.0405 | CIITA | 1.392544977 |
| | Regulation of nuclear beta catenin signaling and target gene transcription(N) | 0.0405 | CYR61 | 1.392544977 |
| | IL-17 signaling pathway(K) | 0.0405 | LCN2 | 1.392544977 |
| | Post-translational protein phosphorylation(R) | 0.0464 | CYR61 | 1.333482019 |
| | Interleukin-4 and Interleukin-13 signaling(R) | 0.0481 | LCN2 | 1.317854924 |
| | Toxoplasmosis(K) | 0.049 | CIITA | 1.30980392 |

| Module | GeneSet | FDR | Nodes | NegLOGFDR |
|---|---|---|---|---|
| M3 | Ether lipid metabolism(K) | 0.0581 | ENPP2 | 1.235823868 |
| | Interferon gamma signaling(R) | 0.0581 | GBP1 | 1.235823868 |
| | Glycerophospholipid biosynthesis(R) | 0.0581 | PLA1A | 1.235823868 |
| | NOD-like receptor signaling pathway(K) | 0.0581 | GBP1 | 1.235823868 |
| | Ras signaling pathway(K) | 0.0796 | PLA1A | 1.099086932 |

| Module | GeneSet | FDR | Nodes | NegLOGFDR |
|---|---|---|---|---|
| M4 | Reelin signalling pathway(R) | 7.08E-03 | VLDLR | 2.150236727 |
| | Reelin signaling pathway(N) | 7.42E-03 | VLDLR | 2.12976001 |
| | Lissencephaly gene (LIS1) in neuronal migration and development(N) | 7.42E-03 | VLDLR | 2.12976001 |
| | EPHA forward signaling(N) | 7.42E-03 | EPHA4 | 2.12976001 |
| | Urokinase-type plasminogen activator (uPA) and uPAR-mediated signaling(N) | 7.42E-03 | VLDLR | 2.12976001 |
| | Plasma lipoprotein assembly, remodeling, and clearance(R) | 0.012 | VLDLR | 1.920818754 |
| | EPH-Ephrin signaling(R) | 0.0152 | EPHA4 | 1.818156412 |
| | Axon guidance(K) | 0.0307 | EPHA4 | 1.512861625 |

| Module | GeneSet | FDR | Nodes | NegLOGFDR |
|---|---|---|---|---|
| M5 | Extracellular matrix organization(R) | 2.12E-04 | COL1A1,ADAMTS 2,TLL1 | 3.67434069 |
| | VEGFR3 signaling in lymphatic endothelium(N) | 0.026 | COL1A1 | 1.585026652 |
| | Beta3 integrin cell surface interactions(N) | 0.026 | COL1A1 | 1.585026652 |
| | IL4-mediated signaling events(N) | 0.026 | COL1A1 | 1.585026652 |
| | Beta1 integrin cell surface interactions(N) | 0.026 | COL1A1 | 1.585026652 |
| | ECM-receptor interaction(K) | 0.026 | COL1A1 | 1.585026652 |
| | Protein digestion and absorption(K) | 0.026 | COL1A1 | 1.585026652 |
| | Amoebiasis(K) | 0.026 | COL1A1 | 1.585026652 |
| | AGE-RAGE signaling pathway in diabetic complications(K) | 0.026 | COL1A1 | 1.585026652 |
| | Platelet activation(K) | 0.0323 | COL1A1 | 1.490797478 |
| | Binding and Uptake of Ligands by Scavenger Receptors(R) | 0.0325 | COL1A1 | 1.488116639 |
| | Relaxin signaling pathway(K) | 0.0341 | COL1A1 | 1.467245621 |
| | Integrin signalling pathway(P) | 0.0413 | COL1A1 | 1.384049948 |

| Module | GeneSet | FDR | Nodes | NegLOGFDR |
|---|---|---|---|---|
| M6 | Neutrophil degranulation(R) | 0.0374 | S100A11 | 1.427128398 |

**Table S8 –List of antibodies used for immuno-histochemistry analysis**

| Antibody name | Company | Catalog # | Host | Dilution |
|---|---|---|---|---|
| Anti-Nuclei Antibody, Clone 235-1 (HuNu) | Millipore Sigma | MAB1281 | Mouse | 1:100 |
| Anti-Green Fluorescent Protein (GFP) | Rockland | 600-101-215 | Goat | 1:1000 |
| Anti-Alpha Smooth Muscle Actin (α-SMA/ACTA2) | Abcam | ab5694 | Rabbit | 1:500 |
| Anti-Glial Fibrillary Acidic Protein (GFAP) | Millipore Sigma | AB5804 | Rabbit | 1:1000 |
| Anti-Neurofilament-L (C28e10) | Cell Signaling | 2837 | Rabbit | 1:100 |
| Anti-E-Cadherin (24E10) | Cell Signaling | 3195 | Rabbit | 1:400 |
| Anti-N-Cadherin | Abcam | AB18203 | Rabbit | 1:1000 |
| Anti-Nestin | Novus | NB100-1604 | Chicken | 1:800 |
| Anti Sox2 | Invitrogen | MA1-014 | Mouse | 1:200 |
| Anti-Iba1 Antibody [EPR16588] | Abcam | ab178846 | Rabbit | 1:500 |
| Anti-Brdu (Bu20a) | Cell Signaling | 5292 | Mouse | 1:200 |
| Anti-Col1a1 | Abcam | ab34710 | Rabbit | 1:500 |
| Anti-Fibronectin | Abcam | ab2413 | Rabbit | 1:1000 |
| Anti-CD68 | Abcam | ab125212 | Rabbit | 1:1000 |
| Anti-CD31 | Cell Signaling | 77699 | Rabbit | 1:100 |
| Anti-Survivin | Cell Signaling | 2808 | Rabbit | 1:500 |
| Anti-PCNA | Cell Signaling | 2586 | Mouse | 1:1000 |
| Anti-Cleaved Caspase 3 | Cell Signaling | 9661 | Rabbit | 1:400 |
| Anti-TdTomato | SICGEN | AB8181-200 | Goat | 1:200 |
| Anti- Glial Fibrillary Acidic Protein (GFAP) | EMD Millipore | AB5541 | Chicken | 1:500 |
| Anti- Green Fluorescent Protein (GFP) | Abcam | AB290 | Rabbit | 1:500 |

| Dataset | Number of Slides | IOU (%, mean) | IOU (%, sd) | Range (%) |
|---|---|---|---|---|
| Mouse | 30 | 76.6 | 14.2 | 41.1 - 94.4 |
| TCGA II | 11 | 100* | 0 | 100 |
| TCGA III | 11 | 73.6 | 11.7 | 54.9 - 87.9 |
| TCGA IV | 12 | 75.5 | 12.7 | 52.6 - 97.3 |
| All | 64 | 80.3 | 11.2 | 52.9 - 100 |

**Table S9-** IOU values over the testing set. * No oncostreams were present in TCGA II data and our model did not have any false positive errors for this dataset

**Other Supplementary information**

**Table Supplementary S2. Oncostreams analysis using TCGA-GBM and TCGA-LGG histological tissues database**

**Table Supplementary S4. Oncostreams' differentially gene expression analysis**

**Movies 1 to 4. Glioma dynamics at the tumor core.**
Time lapse confocal imaging of explant brain slice cultures of NPA glioma cores. Movement of tumor cells GFP positive (green) were analyzed within the tumor core. Imaging was obtained every 10 minutes for the duration of 293 cycles.

**Movies 5 to 9. Glioma dynamics at the tumor border.**
Glioma dynamic at the tumor border were analyzed using explant slice cultures glioma model by intracranial implantation of GFP positive NPA cells into the striatum of B6.129(Cg)-Gt(ROSA)26Sortm4(ACTB-**tdTomato**-EGFP)Luo/J- transgenic mice. Normal brain parenchyma is visualized in red. Imaging was obtained every 10 minutes for the duration of 186 cycles.

**Movies 10 to 13. Glioma dynamics of NPAshCOL1A1 tumors**
Cell glioma dynamic of NPA-ShCOL1A1 tumors were analyzed using explant slice cultures glioma model by intracranial implantation of GFP positive NPAshCOL1A1 cells into the striatum of B6.129(Cg)-Gt(ROSA)26Sortm4(ACTB-**tdTomato**-EGFP)Luo/J- transgenic mice. Normal brain parenchyma is visualized in red. Imaging was obtained every 10 minutes for the duration of 220 cycles.

**Movies 14-16. *In vivo* NPA glioma dynamics patterns employing 2-Photon intravital imaging**
GFP positive NPA glioma cells were intracranially implanted into the cortex of the brain of B6.129(Cg)-Gt(ROSA)26Sortm4(ACTB-tdTomato-EGFP)Luo/J- transgenic mice. Glioma dynamics patterns were evaluated by time-lapse intravital imaging using a two-photon microscope. Normal brain parenchyma is visualized in red and the glioma cells in green. Images were obtained every 5 minutes.

**Movie 17. *In vivo* NPAshCOL1A1 glioma dynamics patterns employing 2-Photon intravital imaging**
GFP positive NPAshCOL1A1 glioma cells were intracranially implanted into the cortex of the brain of B6.129(Cg)-Gt(ROSA)26Sortm4(ACTB-tdTomato-EGFP)Luo/J- transgenic mice. Glioma dynamics patterns were evaluated by time-lapse intravital imaging using a two-photon microscope. Normal brain parenchyma is visualized in red and the glioma cells in green. Images were obtained every 5 minutes.

Fig. S1

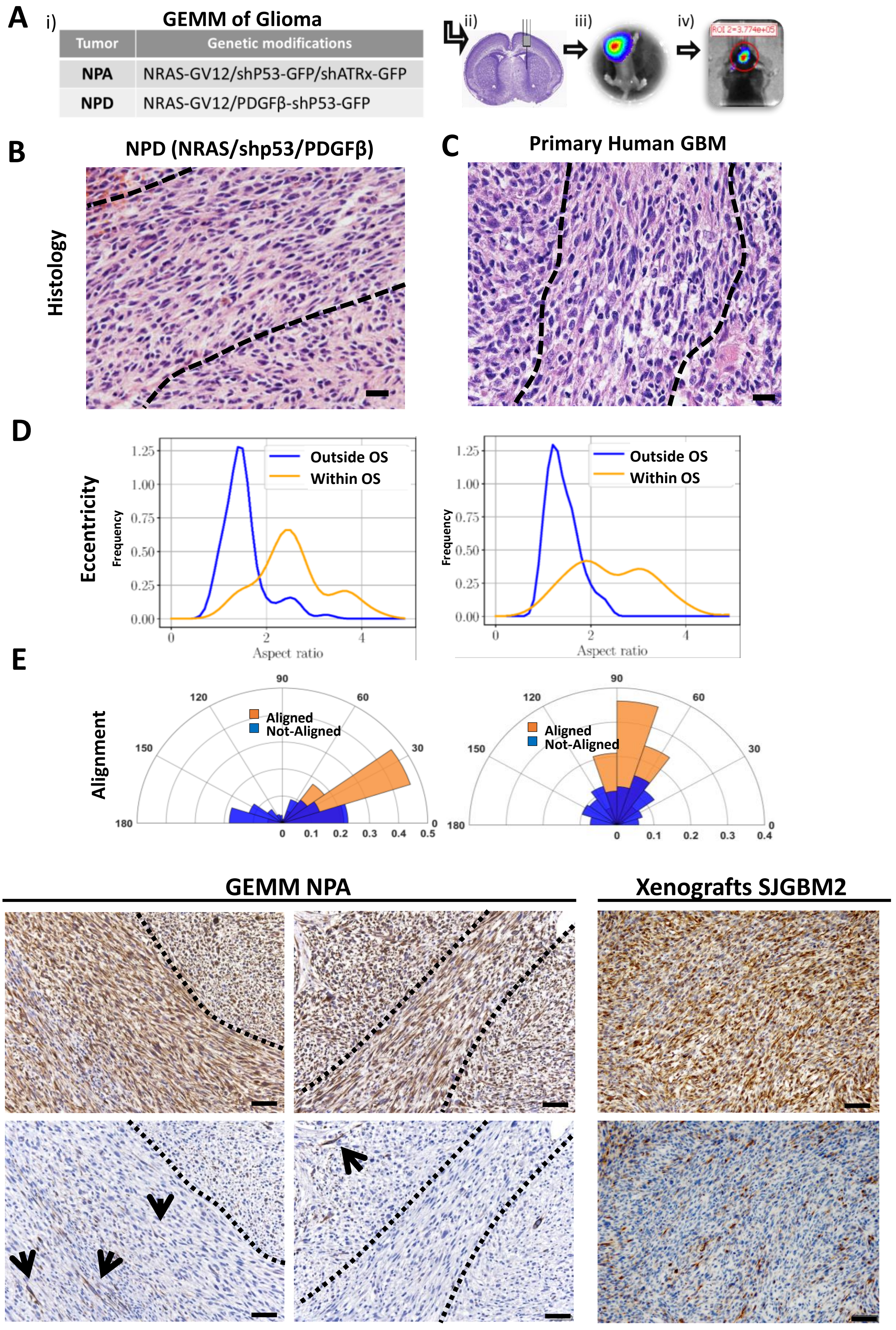
A
i)
GEMM of Glioma
Tumor
Genetic modifications
NPA
NRAS-GV12/shP53-GFP/shATRx-GFP
NPD
NRAS-GV12/PDGFβ-shP53-GFP
ii)
iii)
iv)
B
NPD (NRAS/shp53/PDGFβ)
Histology
C
Primary Human GBM
D
Eccentricity
Frequency
Outside OS
Within OS
Aspect ratio
E
Alignment
Aligned
Not-Aligned
F
GEMM NPA
Xenografts SJGBM2
Nestin
Neurofilaments

**Fig. S1: Oncostream characterization in mouse and human glioma tissue. A)** Schematic representation of the GEMM of glioma generated using the Sleeping Beauty transposase system; i) genetic makeup of NPA and NPD gliomas, ii) plasmid injection into the lateral ventricle of neonatal mice, iii) bioluminescence imaging of a neonatal mouse 1 day post-injection, iv) bioluminescence detection at 70 days post-injection. **B-C)** Representative H&E 5 µm sections from gliomas showing that elongated glioma cells forming oncostreams (dotted line) are present in NPD (NRAS/shp53/PDGFβ) genetically engineered mouse of glioma models (GEMM), and in human glioma tissue. Scale bars: 50 µm. **D**) Cellular eccentricity and alignment analysis on glioma GEMM (NPD) and human glioma representative images. Histograms of cellular aspect ratio **(D)** and cellular alignment **(E)** shows cells within oncostreams are elongated and aligned, whereas outside of oncostreams, they are rounded and not-aligned. **F)** Immunohistochemistry staining for Neurofilaments-L and Nestin in GEMM-NPA tumors and xenograft models (SJGBM2 human cells), showing that oncostreams are not specially organized alongside brain axonal pathways . Scale bars: 50 µm. Arrowheads indicate the presence of Neurofilaments immunoreactivity.

Fig. S2

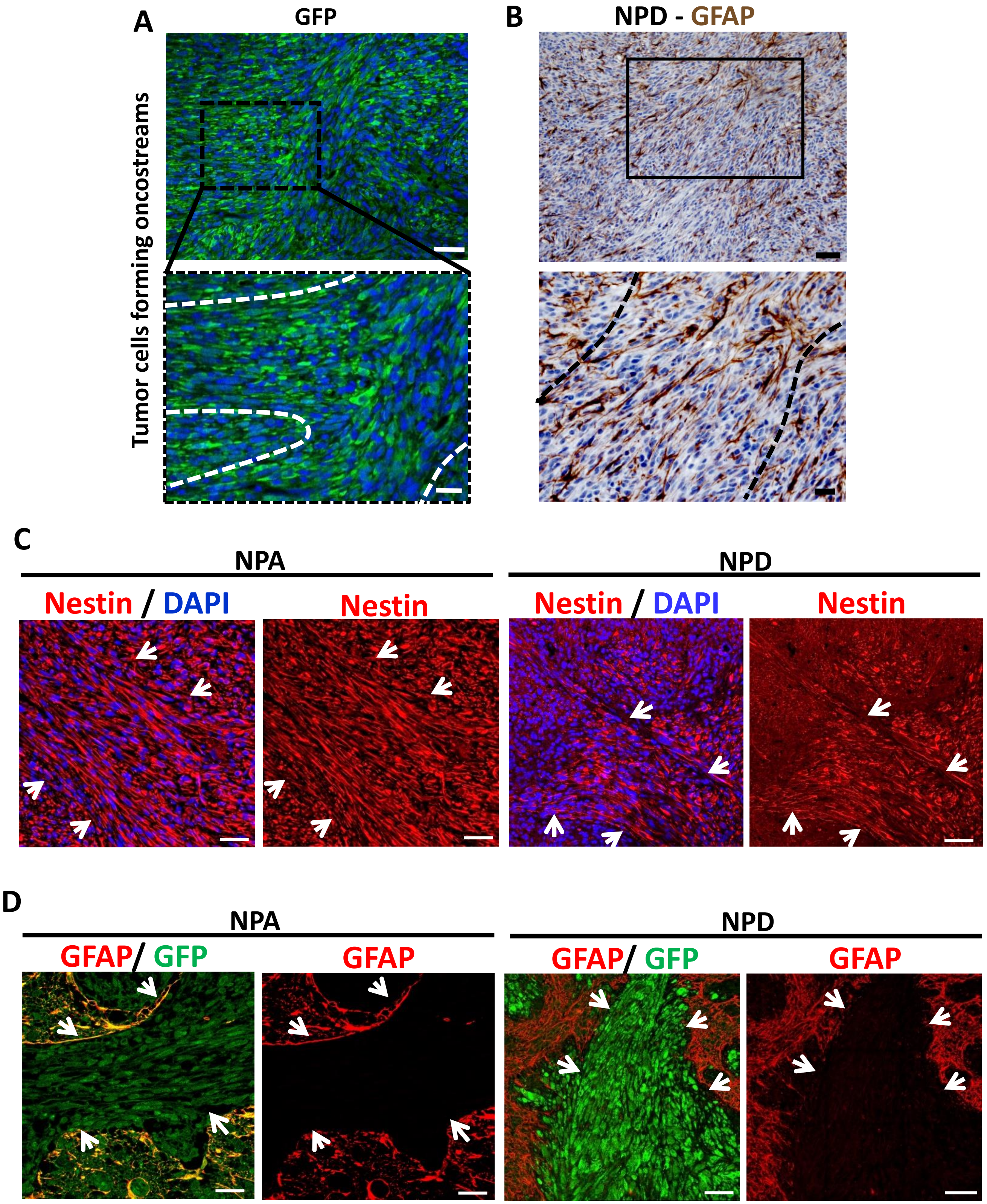
A
GFP
Tumor cells forming oncostreams
B
NPD - GFAP
C
NPA
NPD
Nestin / DAPI
Nestin
Nestin / DAPI
Nestin
D
NPA
NPD
GFAP/ GFP
GFAP
GFAP/ GFP
GFAP

**Fig. S2: Oncostreams are formed by aligned tumor cells and include cells from the tumor microenvironment.**

Oncostreams' cellular heterogeneity was analyzed by immunofluorescence staining. **A)** GFP expression in NPA tumors show oncostreams formed by tumor cells (green = tumor cells; and blue = DAPI stained nuclei). Scale bar: 100 µm (top) and 50 µm (bottom). **B)** Immunohistochemistry analysis on GEMM of glioma (NPD) shows alignment of GFAP+ cells within oncostreams. Scale bars: 50 µm (top) and 20 µm (bottom). **C)** Representative confocal images of GEMM NPA and NPD glioma illustrates the presence of Nestin (red) within oncostreams. Nestin is a highly expressed marker in glioblastomas and expression correlate with tumor prognosis. Nuclei were stained with DAPI (left). Scale bar: 50 µm. **D)** Confocal images of GEMM NPA and NPD glioma show the expression of GFP+ (green) in tumor cells within streams, and tumor microenvironment GFAP+ cells (red) surrounding the oncostream (right panel). GFAP, is expressed by reactive astrocytes or neoplastic astrocytes. Oncostreams either display GFAP+ cells throughout the structures (Fig. S2B) or surrounding the oncostreams (Fig. S2D). GFAP+ cells within oncostreams most likely are tumor cells, while GFAP+ cells surrounding oncostreams are reactive astrocytes. Scale bar: 50 µm. Arrowheads indicate oncostreams areas surrounded by GFAP cells.

Fig. S3

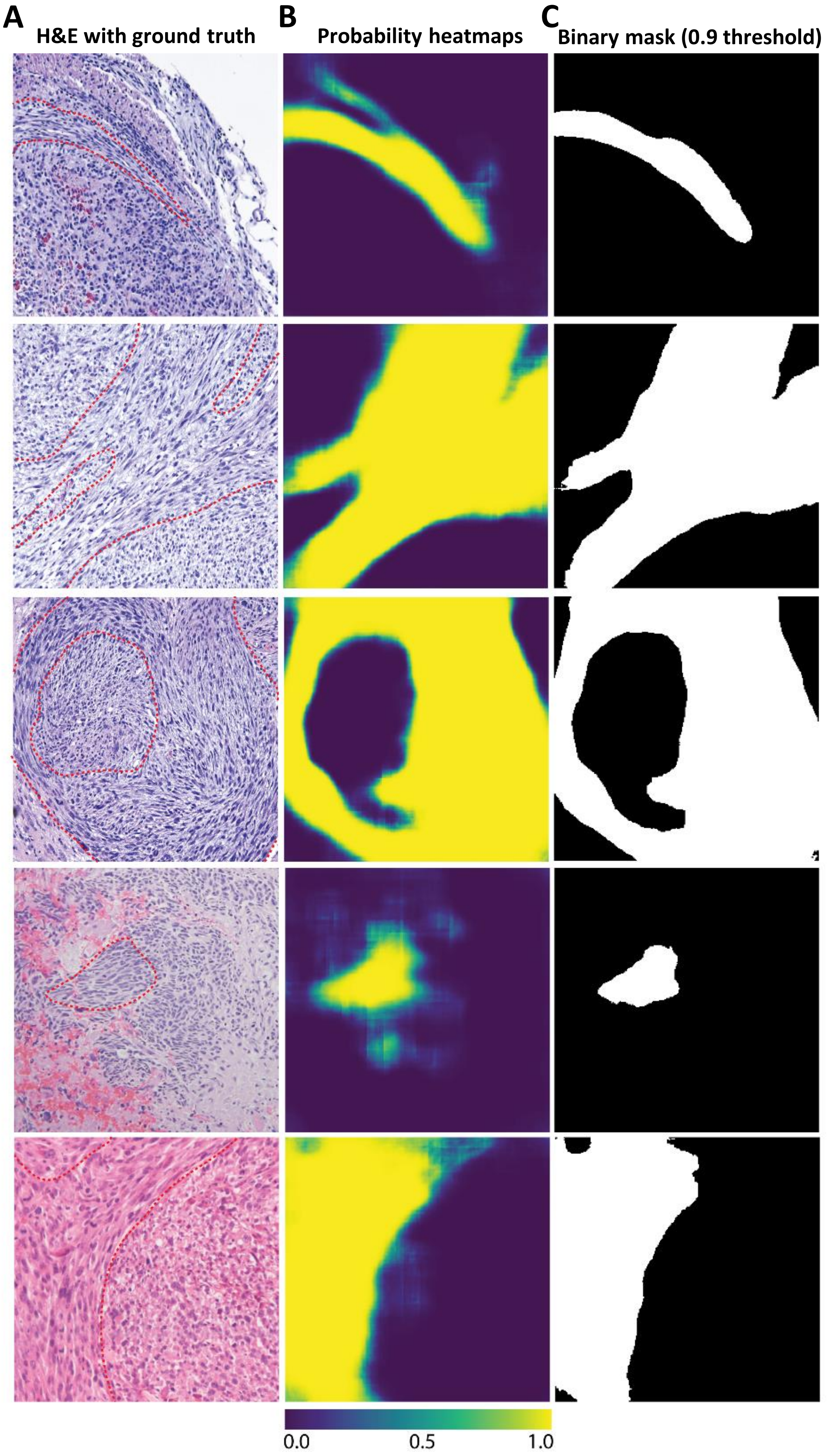
A
H&E with ground truth
B
Probability heatmaps
C
Binary mask (0.9 threshold)
0.0
0.5
1.0

**Fig. S3: A deep learning model identifies oncostreams in mouse glioma H&E histological images.**

We implemented a U-Net architecture to provide semantic segmentation of oncostream fascicles in glioma histological sections. **A)** Mouse H&E images are shown with oncostreams manually segmented (stippled/dashed red lines). **B)** Model output is a semantic segmentation probability heatmap with each pixel being assigned a probability of being within an oncostream (foreground, yellow), or not (background, deep purple). Probability heatmaps for each corresponding H&E images are shown. **C)** Probability heatmaps can be converted to binary masks (foreground versus background) using probability thresholding. Oncostream binary masks with probability threshold of >0.9 are shown. Threshold was tuned during validation as a hyperparameter.

Fig. S4

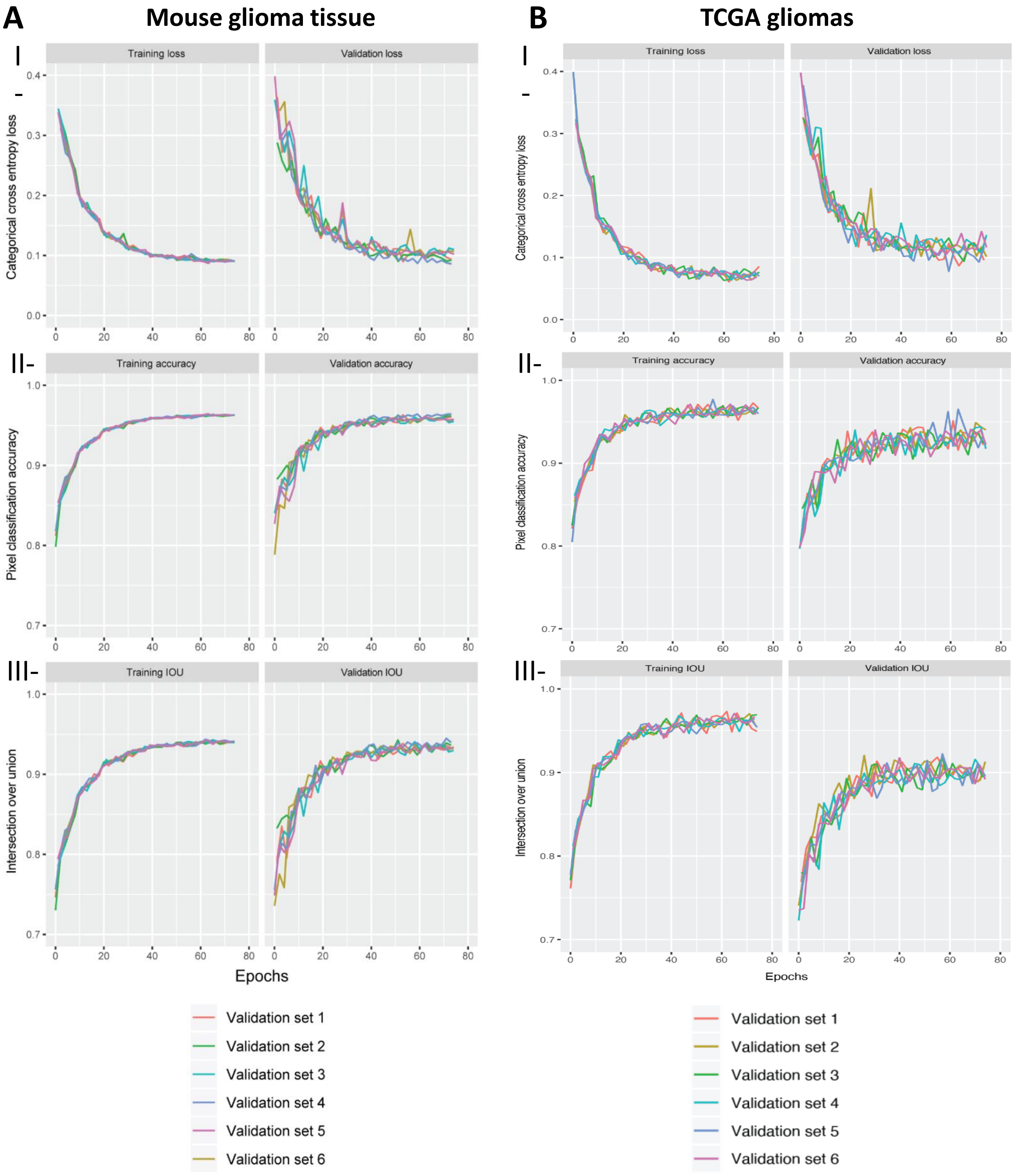


**Fig. S4. Training and validation curves. A-B)** Training and validation curves for a fully convolutional neural network trained using the mouse dataset (**A**) and TCGA dataset (**B**) are shown for the **(I)** cross entropy loss, **(II)** pixel classification accuracy, and **(III)** intersection over union (IOU) metric. We performed 6 iterations of random training-validation dataset splits (80%/20%) and held out the TCGA data for validation. Training and validation metrics stabilized after 75 epochs.

Fig. S5

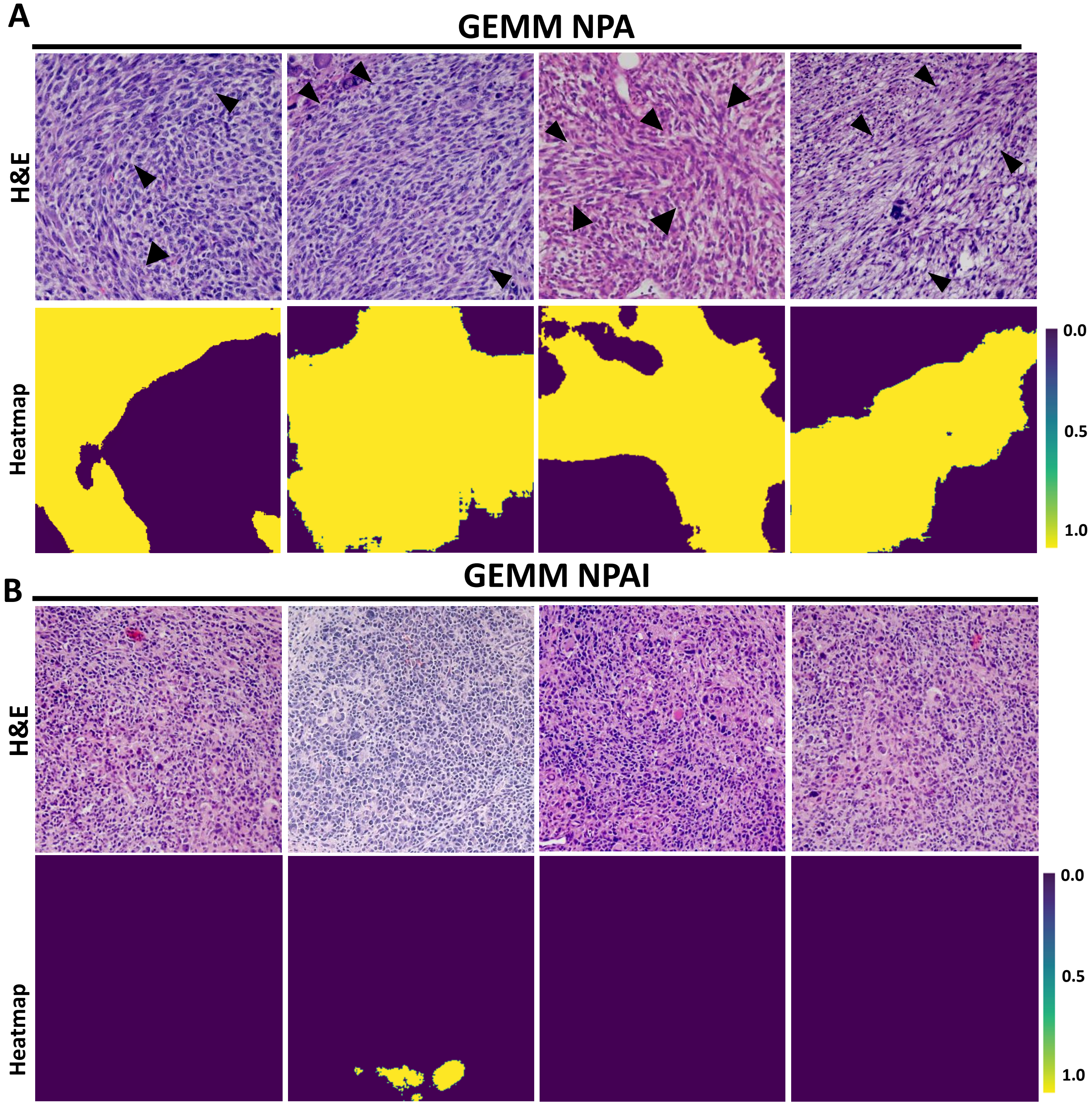


**Fig. S5: Schematic segmentation of oncostreams areas on IDH-WT (NPA) and IDH-Mutant (NPAI) mouse genetically engineered glioma model. A-B)** Representative images of oncostreams manually segmented on H&E stained sections of NPA (IDH-WT) gliomas (**A**) and NPAI (IDH-Mut) gliomas (**B**); (oncostreams are indicated with black arrowheads). 'Heatmap' row illustrates the semantic segmentation probability heatmaps for each corresponding H&E image.

**Fig. S6**

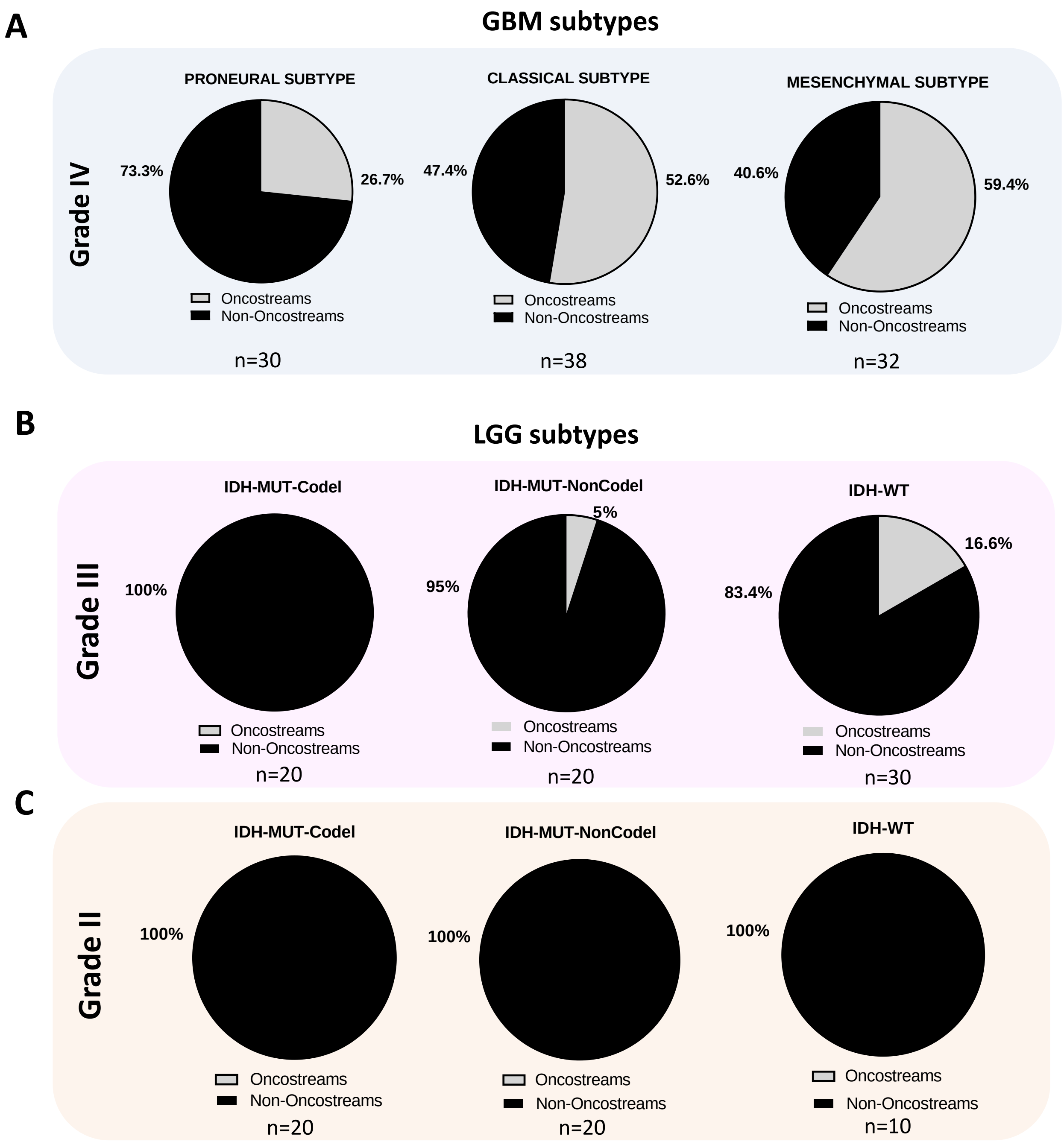


**Fig. S6. A)** Pie charts show percentage of tumors displaying oncostreams in GBM tumors (Grade IV) classified by molecular subtypes. Oncostreams are displayed in Proneural subtype (26.7%), Classical subtype (52.6% %) and Mesenchymal subtype (59.4% %) . **B)** Pie charts show percentage of tumors showing oncostreams in LGG tumors (Grade III) classified by molecular subtypes (IDH1-status and 1p-19q codeletion). Oncostreams are present in 5% of IDH1-Mutant-1p-19q-No-codeletion and in 16.6% of IDH1-WT tumors. **C)** Pie charts show percentage of tumors with oncostreams in LGG tumors (Grade II) classified by molecular subtypes (IDH1-status and 1p-19q codeletion). No oncostreams are observed in Grade II gliomas.

**Fig. S7**

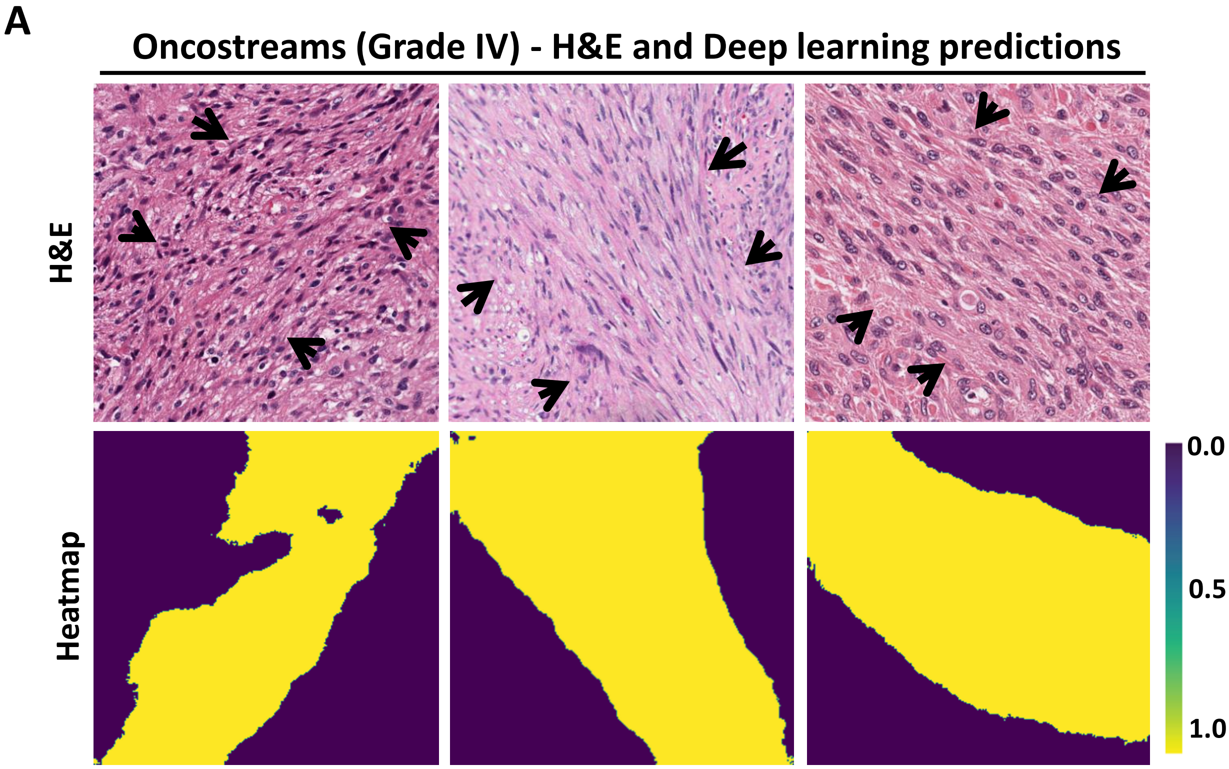


**Fig. S7: Oncostreams are identified by deep learning in human high-grade gliomas (Glioblastoma - Grade IV). A)** Representative H&E images of TCGA-glioblastoma multiforme (WHO Grade IV) diagnostic slides show the presence of manually segmented oncostreams (indicated by black arrows). The second row shows semantic segmentation probability heatmaps for each corresponding H&E image.

Fig. S8

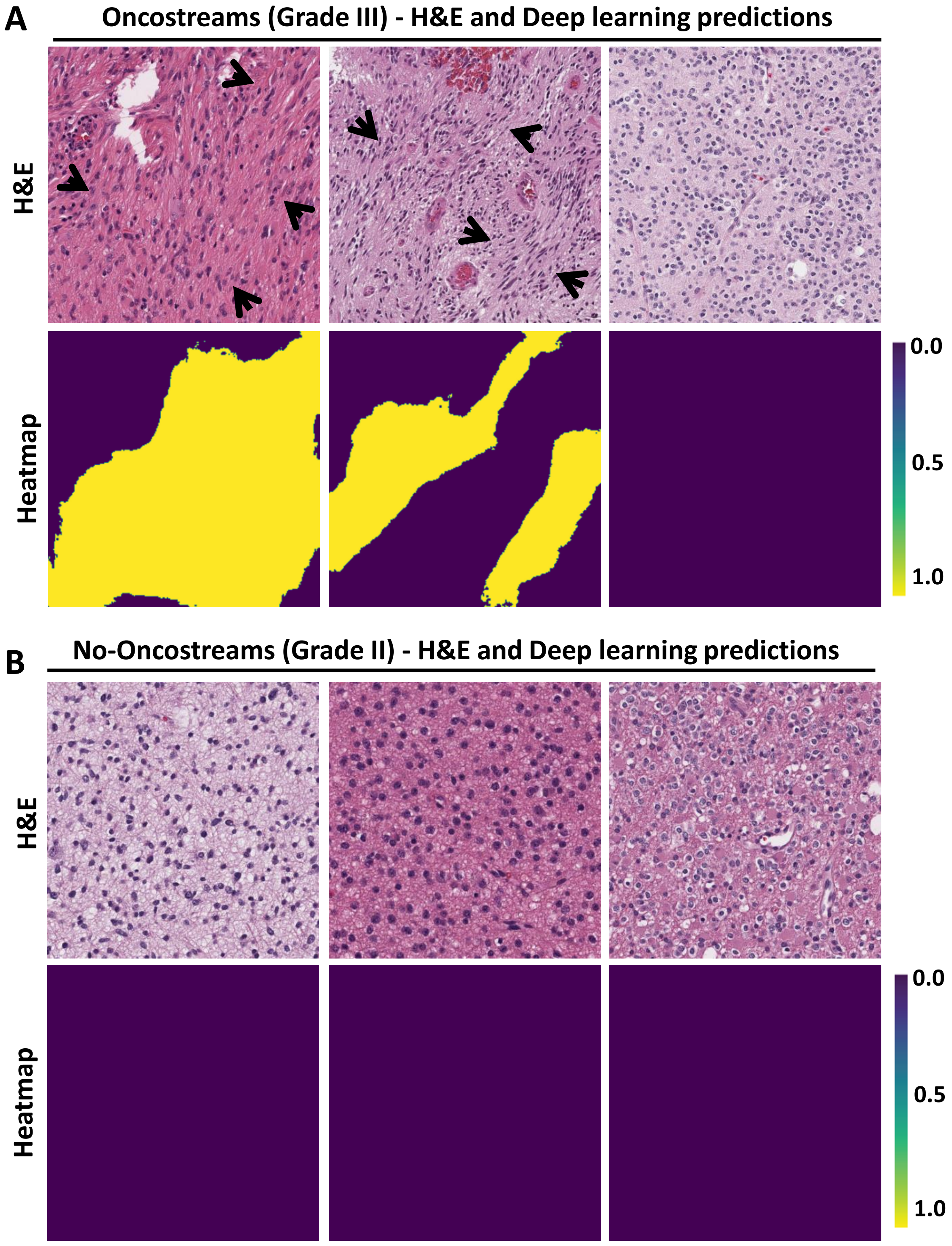


**Fig. S8: Oncostreams are observed in a low percentage of human gliomas WHO grade III but not in WHO grade II. A-B)** Representative H&E images of TCGA gliomas WHO Grade III **(A)** and Grade II **(B)** samples show the presence of manually segmented oncostreams (indicated by black arrows). Semantic segmentation probability heatmaps for each corresponding H&E image are shown in the row labeled 'heatmap'.

Fig. S9

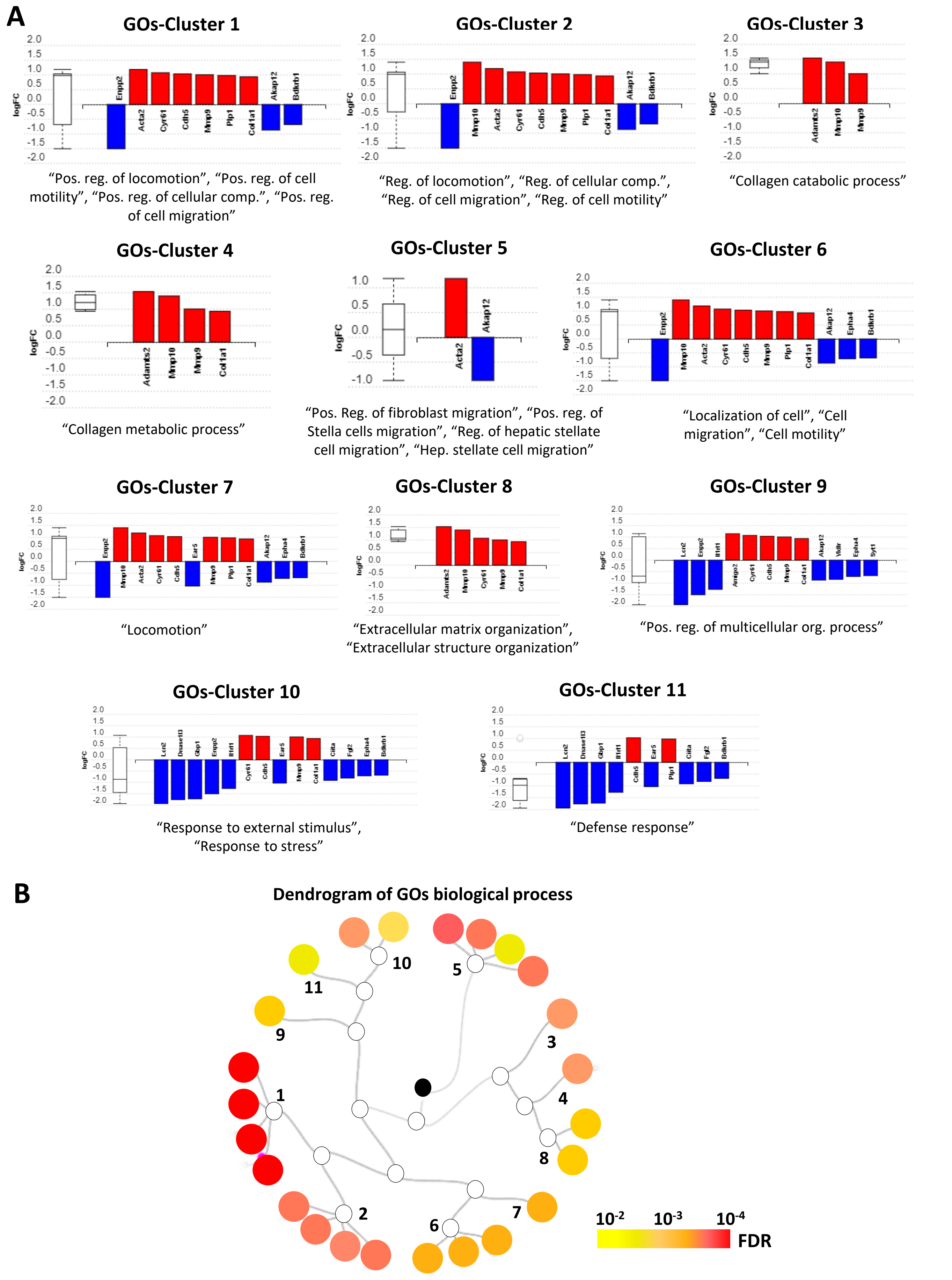
A
GOs-Cluster 1
"Pos. reg. of locomotion", "Pos. reg. of cell motility", "Pos. reg. of cellular comp.", "Pos. reg. of cell migration"
GOs-Cluster 2
"Reg. of locomotion", "Reg. of cellular comp.", "Reg. of cell migration", "Reg. of cell motility"
GOs-Cluster 3
"Collagen catabolic process"
GOs-Cluster 4
"Collagen metabolic process"
GOs-Cluster 5
"Pos. Reg. of fibroblast migration", "Pos. reg. of Stella cells migration", "Reg. of hepatic stellate cell migration", "Hep. stellate cell migration"
GOs-Cluster 6
"Localization of cell", "Cell migration", "Cell motility"
GOs-Cluster 7
"Locomotion"
GOs-Cluster 8
"Extracellular matrix organization", "Extracellular structure organization"
GOs-Cluster 9
"Pos. reg. of multicellular org. process"
GOs-Cluster 10
"Response to external stimulus", "Response to stress"
GOs-Cluster 11
"Defense response"
logFC
B
Dendrogram of GOs biological process
10^-2 10^-3 10^-4
FDR

**Fig. S9. Enrichment in migration and extracellular matrix biological process suggest oncostreams are mesenchymal migratory structures.**

**A)** Bar graphs show clusters of enriched GOs biological process sharing same differentially expressed genes. Pos: Positive, Reg: Regulation. **B)** Dendrogram of GOs biological process. Significant results are organized hierarchically based on overlap in associated DE genes. Node color represent significant differences (FDR corrected p-value), black dot represent the center of the dendrogram.

**Fig. S10**

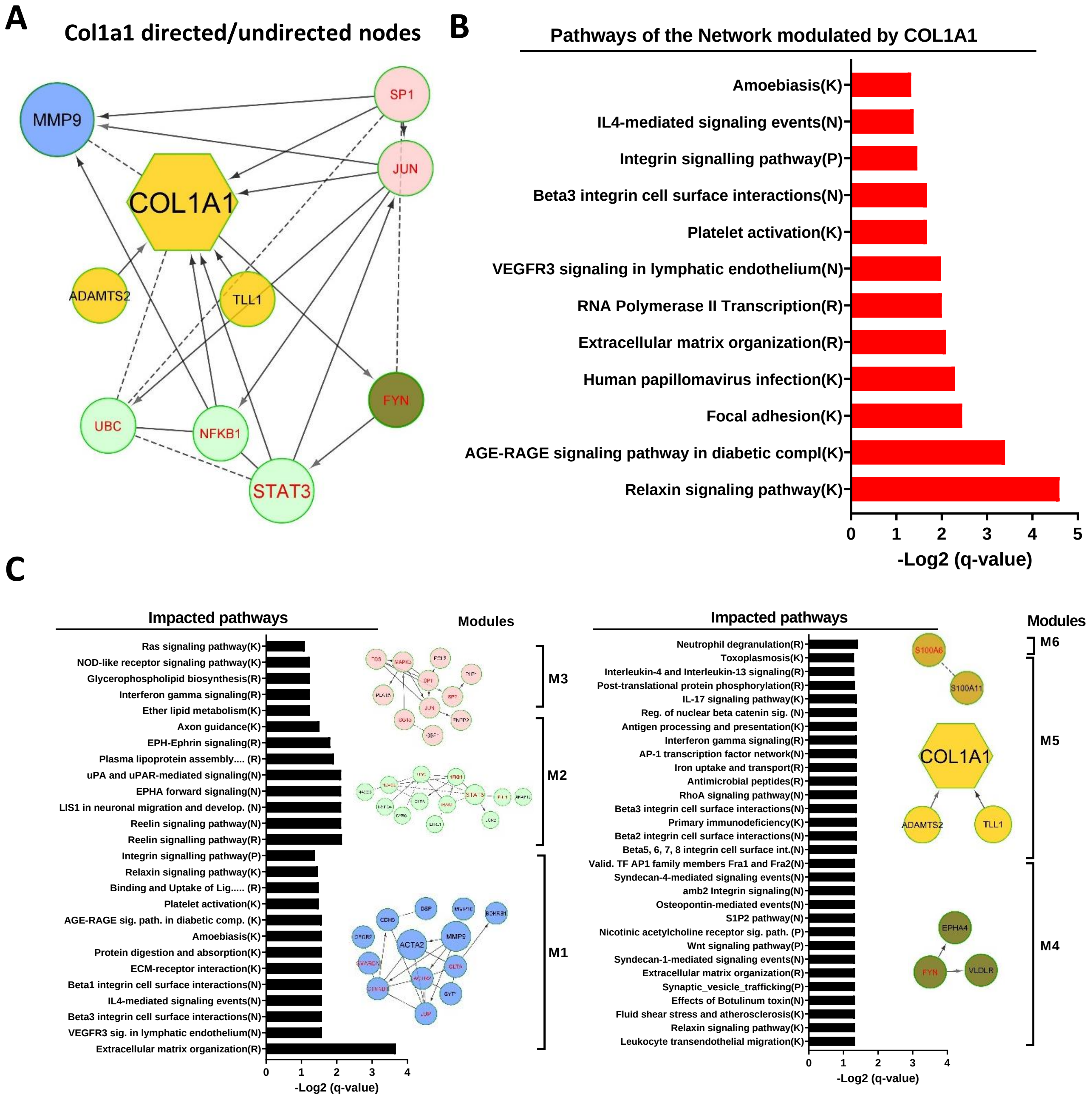


**Fig. S10. Pathway enrichment analyses of the COL1A1 network derived from DE genes of oncostreams vs no-oncostreams. A)** COL1A1 network showing its first neighbors. COL1A1 is a hub node with a degree of 9 interactions in the network (2nd most connected DE node). **B)** Functional enrichment analysis of the gene ontology (GO) terms of the COL1A1 network. The bar graph displays over-represented GOs biological process that include COL1A1. GO term significance was determined by a cutoff of q-value (FDR) < 0.05. GO terms were plotted against the minus Log 2 of the q-value (FDR). **C)** Functional enrichment analysis of the impacted pathways in each module of the network. Modules (nodes of the same color) represent a highly interacting group of genes in the network. The bar graph shows the overrepresented pathways plotted according to the minus Log2 q-value (FDR). Pathway-selection cutoff was set at a q-value (FDR) <0.05.

### Fig. S11

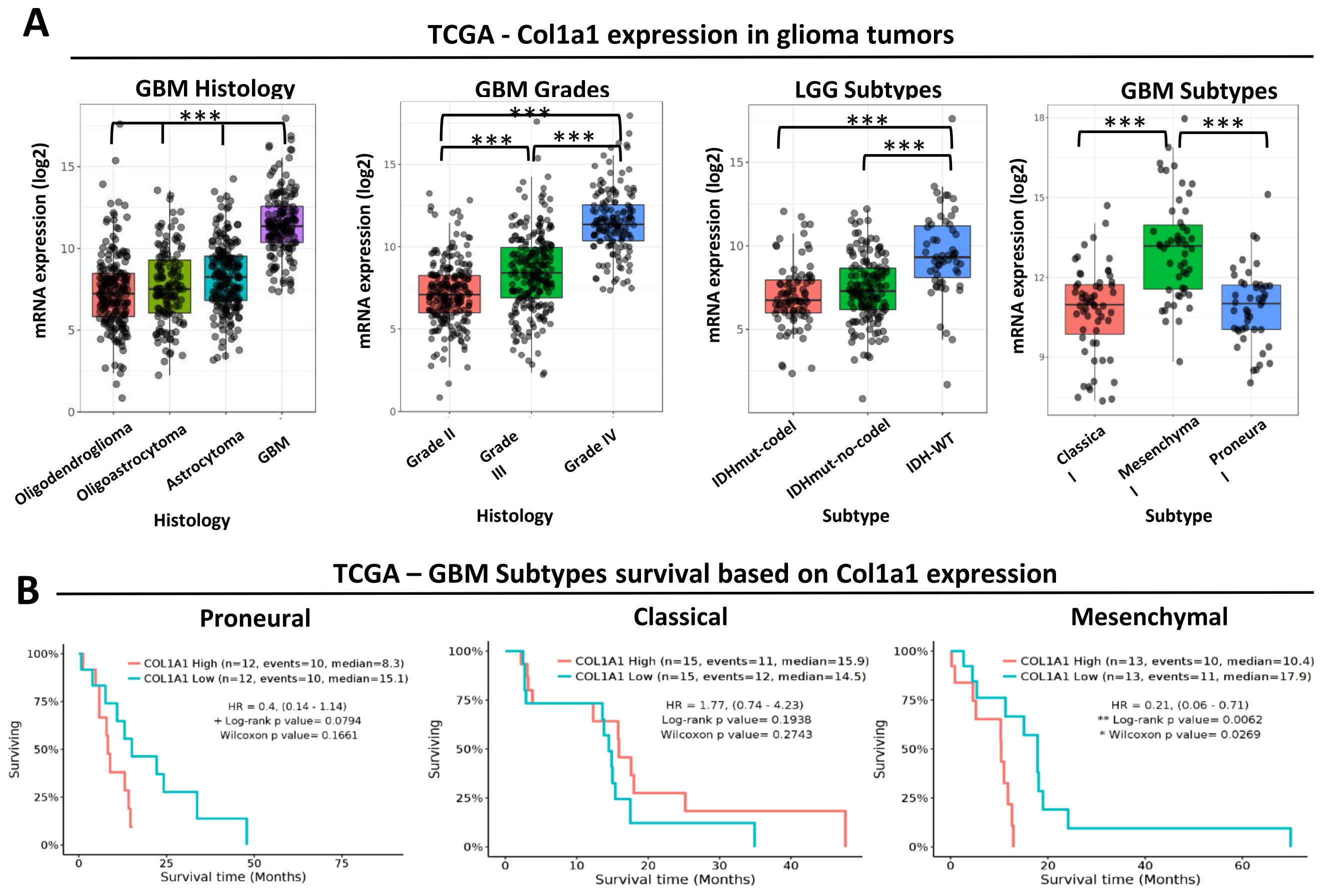


**Fig. S11. Evaluation of COL1A1 expression levels associated with human glioma aggressiveness and clinical outcomes. A)** mRNA expression analysis of COL1A1 related to glioma tumor histology, grade and LGG-subtypes and GBM-subtypes (TCGA-GBMLGG database). Graph shows the $\log_2$ mRNA expression levels. Statistical significance was given within the corresponding databases (Tukey's Honest Significant Difference (HSD). **B)** Kaplan-Meier survival curves of GBM patients comparing COL1A1 high vs low expression in GBM molecular subtypes (TCGA-GBMLGG database). Data was analyzed from the Gliovis website (http://gliovis.bioinfo.cnio.es). GBM tumors with high COL1A1 expression show a significant increased survival compared to GBM COL1A1 low within the mesenchymal subtype.

**Fig. S12**

**A**

| shRNA | Sequence |
|---|---|
| Scramble | GAATCTAATCGTATCGTGGCTG |
| shCol1a1-A | CCCTGGTGATACTGGTGTTAAA |
| shCol1a1-B | GCAACAGTCGCTTCACCTACA |
| shCol1a1-C | GCAAGACAGTCATCGAATACA |

**B**

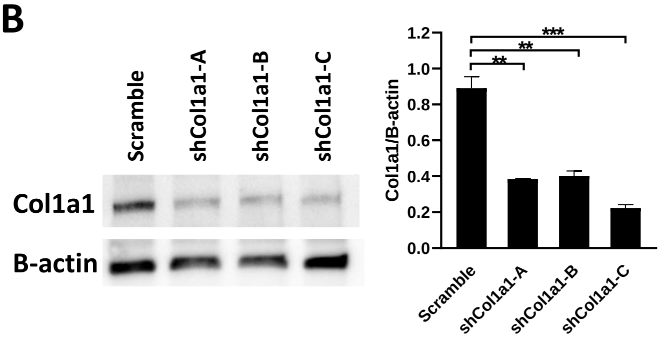


**C**

| Tumor Model | Genetic Modifications |
|---|---|
| NPA | Nras-GV12/shP53/shATRx |
| NPAshCol1a1 | Nras-GV12/shP53/shATRx/shCol1a1 |
| NPD | Nras-GV12/shP53/PDGFβ |
| NPDshCol1a1 | Nras-GV12/shP53/PDGFβ/shCol1a1 |

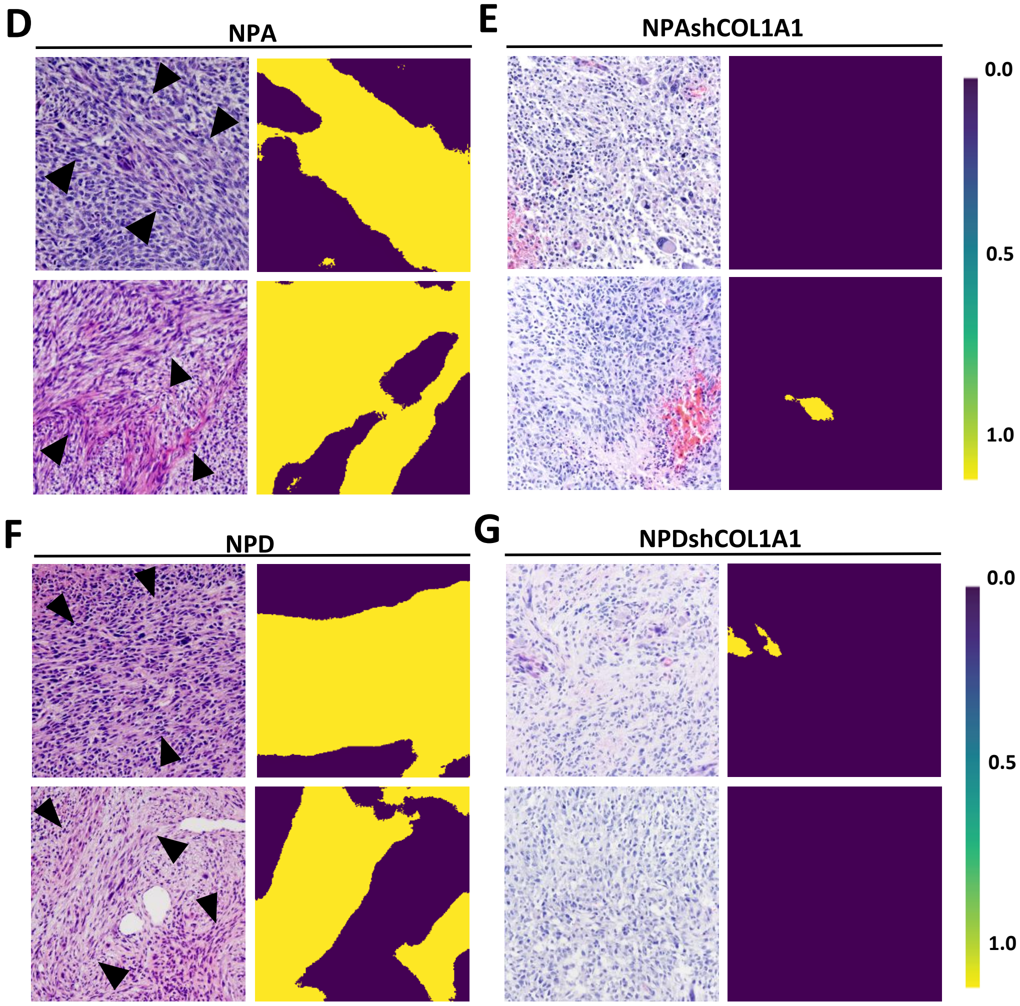

**Fig. S12. Genetically engineered mouse glioma models for COL1A1 knockdown.**

**A)** Scramble and shRNAs sequences. These oligonucleotides were cloned in PT2 vectors. **B)** Western Blot (WB) analysis shows downregulation of COL1A1 expression levels in NIH-3T3 mouse cells transfected for 48 hours with the corresponding vectors (Scramble control and 3 different shRNAs). Bar graphs represent the quantitative analysis of the WB. Values were calculated by normalizing COL1A1 expression to β-actin. Quantification was performed using ImageJ software. Statistical significance was determined using One-way ANOVA test. **$p < 0.01$, ***$p < 0.001$. Error bars represent ±SEM. **(C)** Genetic modifications used to induce different mouse glioma models. **D-G)** Representative images of oncostreams manually segmented on H&E stained sections of GEMM of NPA (**D**), NPAshCOL1A1 **(E)**, NPD **(F)** and NPDshCOL1A1 **(G)** gliomas (oncostreams are indicated with black arrowheads). 'Heatmap outputs illustrate the semantic segmentation probability heatmaps for each corresponding H&E image.

Fig. S13

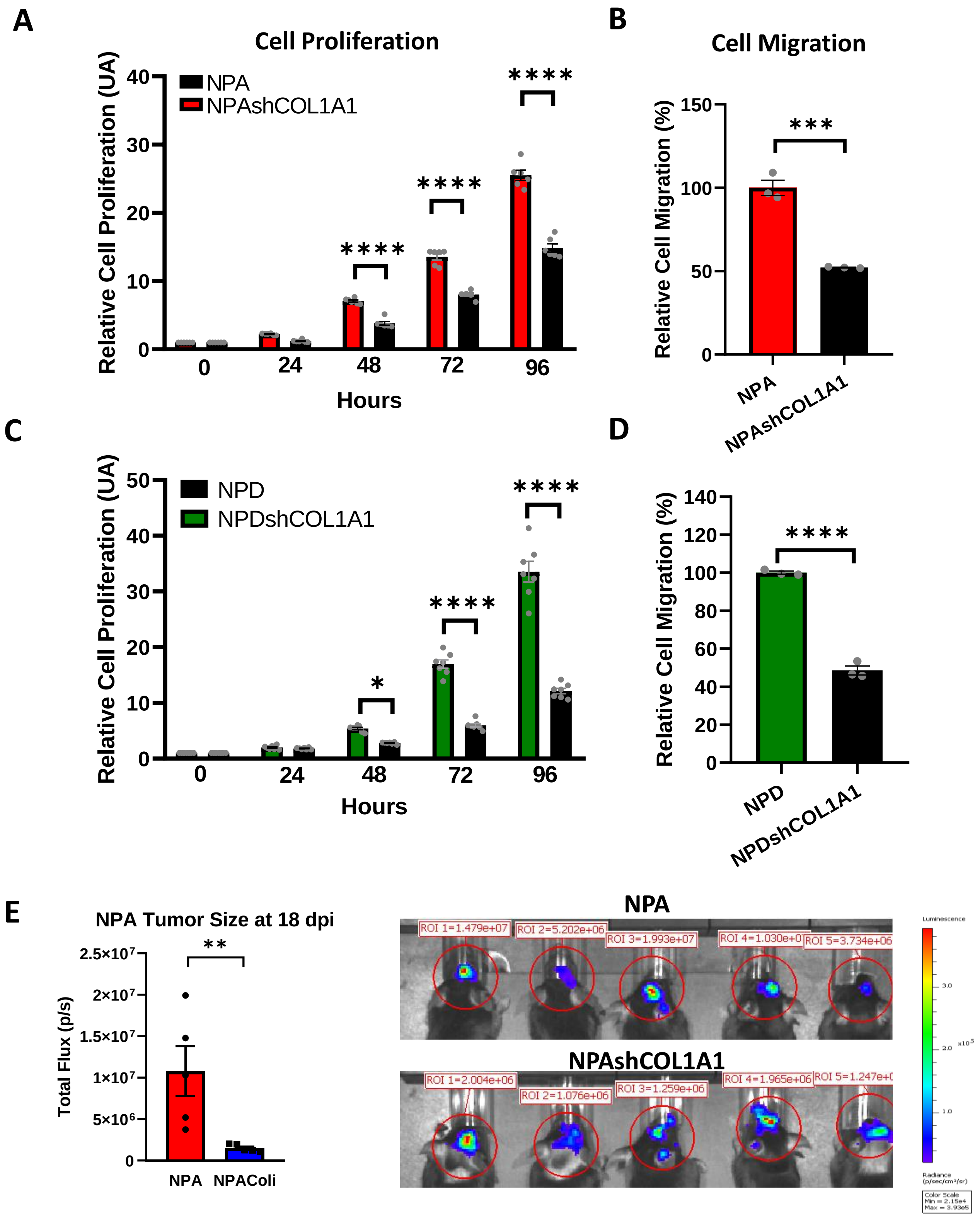

A
Cell Proliferation
Relative Cell Proliferation (UA)
NPA
NPAshCOL1A1
0
10
20
30
40
****
****
****
0
24
48
72
96
Hours
B
Cell Migration
Relative Cell Migration (%)
0
50
100
150
***
NPA
NPAshCOL1A1
C
Relative Cell Proliferation (UA)
NPD
NPDshCOL1A1
0
10
20
30
40
50
*
****
****
0
24
48
72
96
Hours
D
Relative Cell Migration (%)
0
20
40
60
80
100
120
140
****
NPD
NPDshCOL1A1
E
NPA Tumor Size at 18 dpi
Total Flux (p/s)
0
5×10^6
1×10^7
1.5×10^7
2×10^7
2.5×10^7
**
NPA
NPAColi
NPA
ROI 1=1.479e+07
ROI 2=5.202e+06
ROI 3=1.993e+07
ROI 4=1.030e+0
ROI 5=3.734e+06
NPAshCOL1A1
ROI 1=2.004e+06
ROI 2=1.076e+06
ROI 3=1.259e+06
ROI 4=1.965e+06
ROI 5=1.247e+0

**Fig. S13. Downregulation of COL1A1 decreases cell proliferation and cell migration *in vitro*, and tumor growth *in vivo.***

Neurospheres obtained from GEMM of glioma controls (NPA and NPD) and COL1A1 downregulation (NPAshCOL1A1 and NPDshCOL1A1) were employed to determine cell proliferation and cell migration. **A** and **C)** Cell proliferation was assessed at 0, 24, 48 and 96 h (n=6 replicates) and was determined using Cell Titer-Glo Assay. **B** and **D)** Cell migration was measured after 15 h using transwell assay, and number of cells was determined using Cell Titer-Glo Assay. Data are expressed as percentage of migrating cells relative to the control. Error bars represent ±SEM. n=3,*p<0.05, ****p<0.0001. **E)** Tumor growth was determined measuring *in vivo* luminesce of C57BL mice bearing NPA or COL1A1 downregulation (NPAshCOL1A1) at 18 days post-implantation (n=5 replicates). Error bars represent ±SEM. t-test, **P<0.01.

Fig. S14

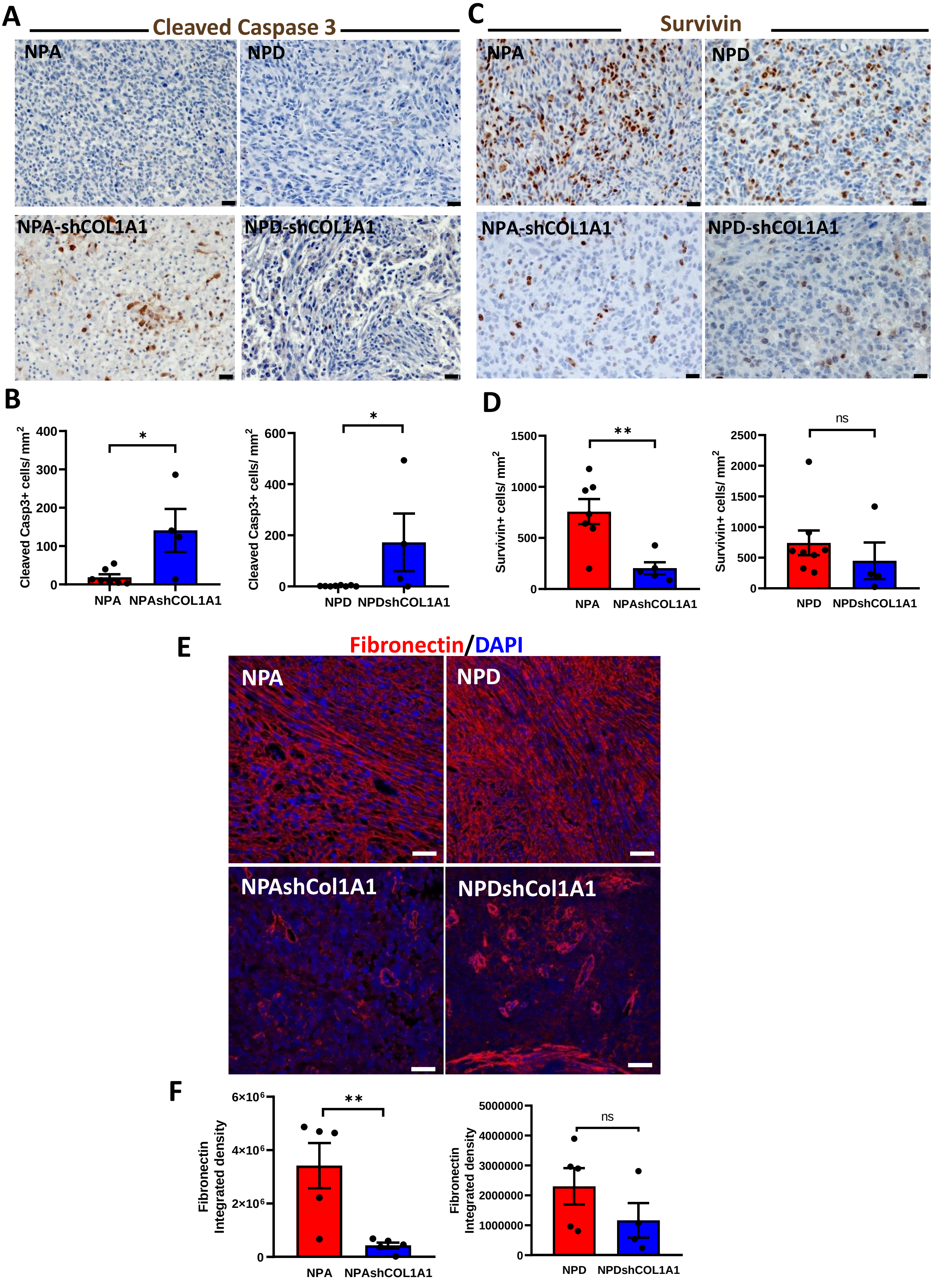

A
Cleaved Caspase 3
NPA
NPD
NPA-shCOL1A1
NPD-shCOL1A1
C
Survivin
NPA
NPD
NPA-shCOL1A1
NPD-shCOL1A1
B
Cleaved Casp3+ cells/ mm2
400
300
200
100
0
*
NPA
NPAshCOL1A1
600
400
200
0
*
NPD
NPDshCOL1A1
D
Survivin+ cells/ mm2
1500
1000
500
0
**
NPA
NPAshCOL1A1
2500
2000
1500
1000
500
0
ns
NPD
NPDshCOL1A1
E
Fibronectin/DAPI
NPA
NPD
NPAshCol1A1
NPDshCol1A1
F
Fibronectin Integrated density
6×10^6
4×10^6
2×10^6
0
**
NPA
NPAshCOL1A1
5000000
4000000
3000000
2000000
1000000
0
ns
NPD
NPDshCOL1A1

**Fig. S14. COL1A1 genetic downregulation induced changes in apoptosis and decreased the expression of the mesenchymal marker fibronectin.**

Immunohistochemical analysis (**A** and **C**) on GEMM of glioma controls (NPA and NPD) and COL1A1 downregulation (NPAshCOL1A1 and NPDshCOL1A1). **A)** Representative images of Cleaved Caspase 3 expression (DAB). Scale bar: 20 µm. **B)** Bar graphs represent Cleaved Caspase 3+ cells quantification (number of cell/$mm^2$) using QuPath positive cells detection. Error bars represent ±SEM, (NPA: n=7, NPAshCOL1A1: n=4, NPD: n=8, NPDshCOL1A1: n=4), t-test, *p<0.05. **C)** Representative images of Survivin expression (DAB). Scale bar: 20 µm**.**. **D)** Bar graphs represent Survivin+ cells quantification (number of cell/$mm^2$) using QuPath positive cells detection. Error bars represent ±SEM, (NPA: n=7, NPAshCOL1A1: n=5, NPD: n=8, NPDshCOL1A1: n=4), t-test, **p<0.001, ns: no significant. **E)** Immunofluorescence analysis on GEMM of glioma controls (NPA and NPD) and COL1A1 downregulation (NPAshCOL1A1 and NPDshCOL1A1). Representative images of Fibronectin expression in red (Alexa 555) and nuclei in blue (DAPI). Scale bar: 50 µm**. F)** Bar graphs represent Fibronectin quantification in terms of fluorescence integrated density. Error bars represent ±SEM, (NPA: n=5, NPAshCOL1A1: n=5, NPD: n=5, NPDshCOL1A1: n=4), t-test, **p<0.001, ns: no significant.

#### Fig. S15

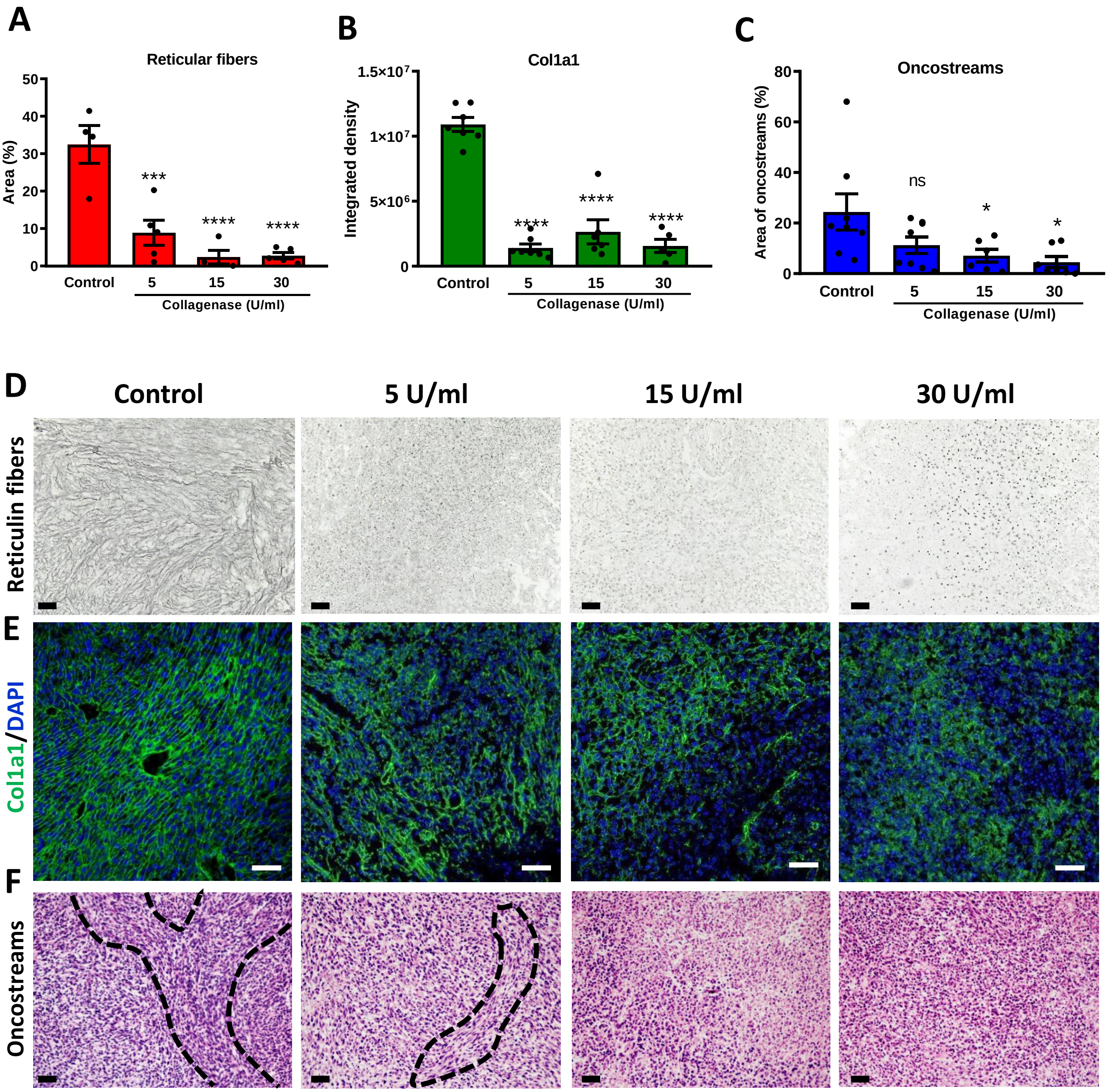


**Fig. S15. Collagenase treatment depletes deposited collagen fibers and eliminated oncostream mesenchymal fascicles.** <u>**A-F)**</u> Explant glioma brain slides were treated with different concentrations of collagenase (5 U/ml, 15 U/ml and 30 U/ml) for 48 hours. **A)** Bar graphs show the quantification of reticular fibers (Reticulin Silver Plating Kit acc. to Gordon & Sweets). Error bars represent ±SEM. Control: n=8, 5 U/ml: n=8, 15 U/ml: n=6, 30 U/ml: n=7. One-way ANOVA, ***p<0.001, ****p<0.0001. **B)** Bar graphs represent COL1A1 quantification in terms of fluorescence integrated density. Error bars represent ±SEM. Control: n=7, 5 U/ml: n=7, 15 U/ml: n=6, 30 U/ml: n=5. One-way ANOVA, ****p<0.0001. **C)** Identification and quantification of oncostream areas using Image J. Error bars represent ±SEM. Control: n=8, 5 U/ml: n=8, 15 U/ml: n=6, 30 U/ml: n=7. One-way ANOVA, *p<0.05, ns: no significant. **D-F)** Representative images display reticular fiber expression and organization (**D**); COL1A1 expression and organization in green (Alexa 488) and nuclei in blue (DAPI) (**E**); and H&E stained paraffin sections (**F**). Images (20x objective) Scale bars: 50 µm.

Fig. S16

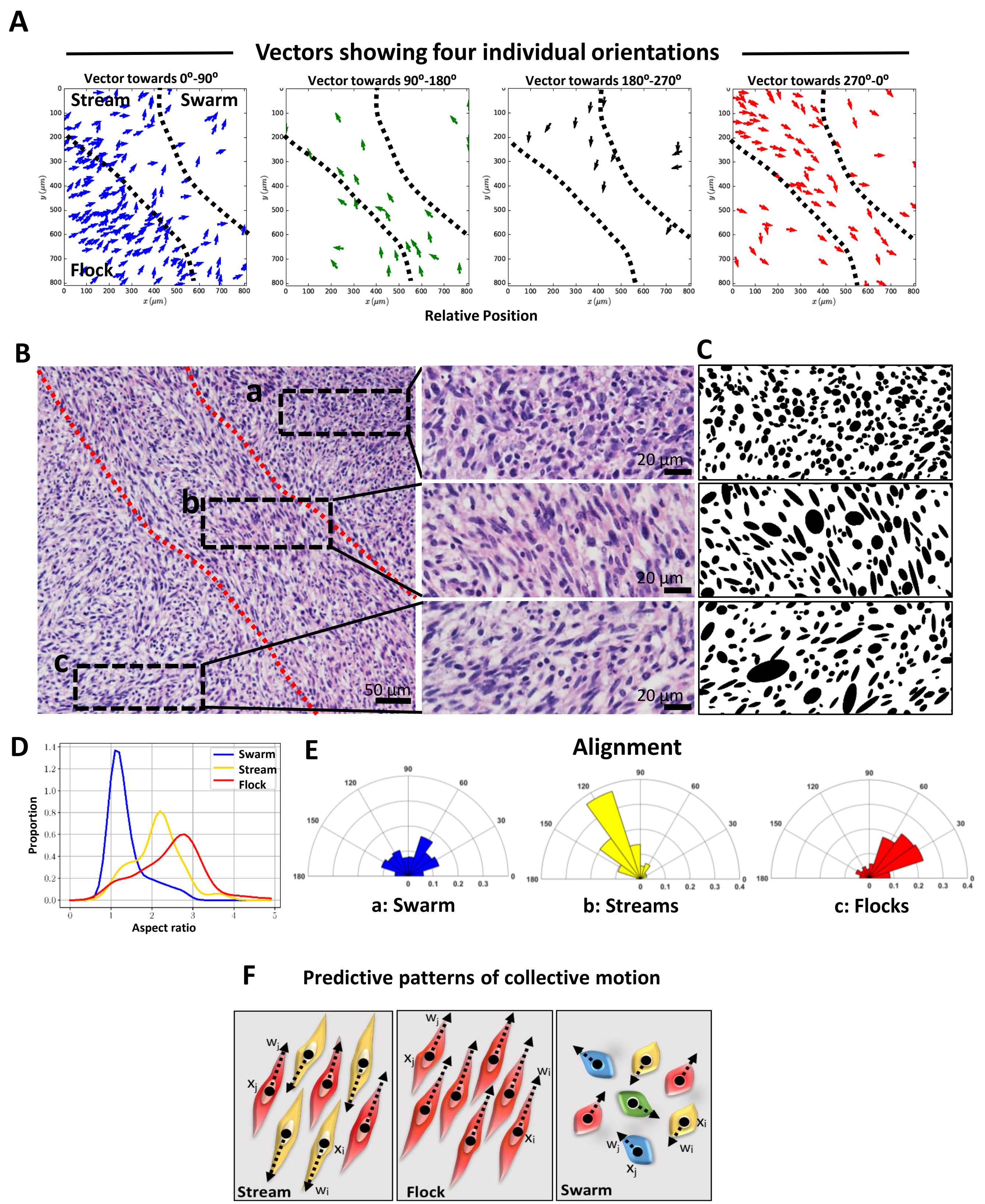
A
Vectors showing four individual orientations
Vector towards 0°-90°
Vector towards 90°-180°
Vector towards 180°-270°
Vector towards 270°-0°
Stream
Swarm
Flock
y (μm)
x (μm)
Relative Position
B
C
a
b
c
20 μm
50 μm
D
Swarm
Stream
Flock
Proportion
Aspect ratio
E
Alignment
a: Swarm
b: Streams
c: Flocks
F
Predictive patterns of collective motion
Stream
Flock
Swarm

**Fig. S16. Dynamic patterns are aligned to cellular morphology and histological features in gliomas. A)** Vectors indicating average direction per cell for each area (from Fig. 7B-D). **B)** H&E staining of a 5 μm microtome section taken from an organotypic slice culture glioma model used for confocal imaging and shown in **Fig. 7B.** Dotted white lines define different zones which display different dynamic patterns in **Fig. 7D**, i.e., a=*swarm*, b=stream and c=flock, and boxes are shown at higher power to the right. **C)** Masks generated by particle analysis in Image J were used to calculate the alignments shown in Fig. 2D. **D)** Histogram of cell eccentricity analysis of organotypic H&E stained glioma sections. Aspect ratio of cells are shown for a swarm (blue; ~1), a stream (yellow; >2) and a flock (red; >2). **E)** Alignment analysis of cells: Angle histogram plots show areas of high proportion of aligned cells (stream and flock) (narrow range of angle orientation), and areas of no preferred orientation (swarm). **F)** Dynamic self-organizing collective motion patterns predicted by our *in silico* agent-based modeling, in which cell eccentricity and density determine pattern formation: 1. Stream (↑↓), 2. Flock (↑↑), 3. Swarm (↕↔).

**Fig. S17**

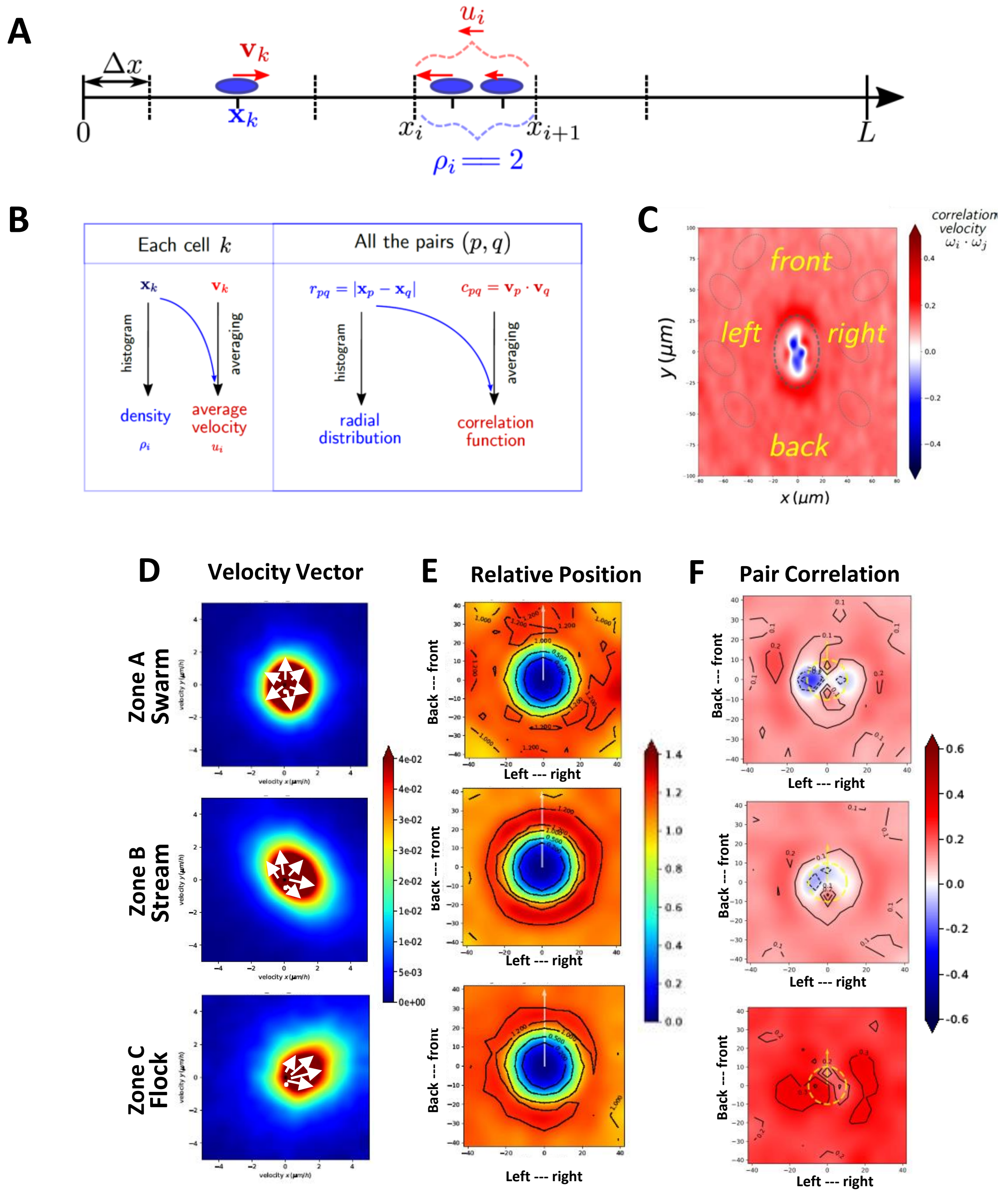
A
$\Delta x$
$\mathbf{v}_k$
$\mathbf{x}_k$
$u_i$
0
$x_i$
$x_{i+1}$
$L$
$\rho_i == 2$
B
Each cell $k$
All the pairs $(p, q)$
$\mathbf{x}_k$
$\mathbf{v}_k$
histogram
averaging
density
$\rho_i$
average velocity
$u_i$
$r_{pq} = |\mathbf{x}_p - \mathbf{x}_q|$
$c_{pq} = \mathbf{v}_p \cdot \mathbf{v}_q$
radial distribution
correlation function
C
front
left
right
back
$y\,(\mu m)$
$x\,(\mu m)$
correlation velocity
$\omega_i \cdot \omega_j$
D
Velocity Vector
E
Relative Position
F
Pair Correlation
Zone A Swarm
Zone B Stream
Zone C Flock
Back --- front
Left --- right

**Fig. S17: Statistical analysis performed to study glioma dynamics.**

**A)** Correlation functions: estimation of the density $\rho_i$ and average velocity $u_i$. **B)** Radial distribution and correlation function are estimated in the same way as density and average velocity, but we use pairwise information (e.g. distances between neighbors). **C)** Heat map plot of the pair wise correlation of the velocity. We estimated the correlation depending on the position of a nearby $x_j$ in different positions (front-to-back and left-to-right). Positive correlation (+) indicates cells are moving in same directions (red) and negative correlation (-) cells are moving in opposite directions (blue). **D)** Heat map plot of the distribution of the velocity vector in different zones of the tumor core. Velocity is shown in the x-axis and y-axis in µm/hr. **E)** Frequency for each zone of the relative position between two cells. In the frame of reference of one cell, we estimated the probability of having another cell, $x_j$, nearby. **F)** Pair-wise correlation with nearby neighbors. Depending on the relative position $\mathbf{x}_j$-$\boldsymbol{x}_j$, we estimate the correlation between the velocity directions $\boldsymbol{\omega}_i$ and $\boldsymbol{\omega}_j$. Axes indicate cell position and distance in µm of the neighboring cells. Dotted yellow line show the average size of the centered cell.

**Fig. S18**

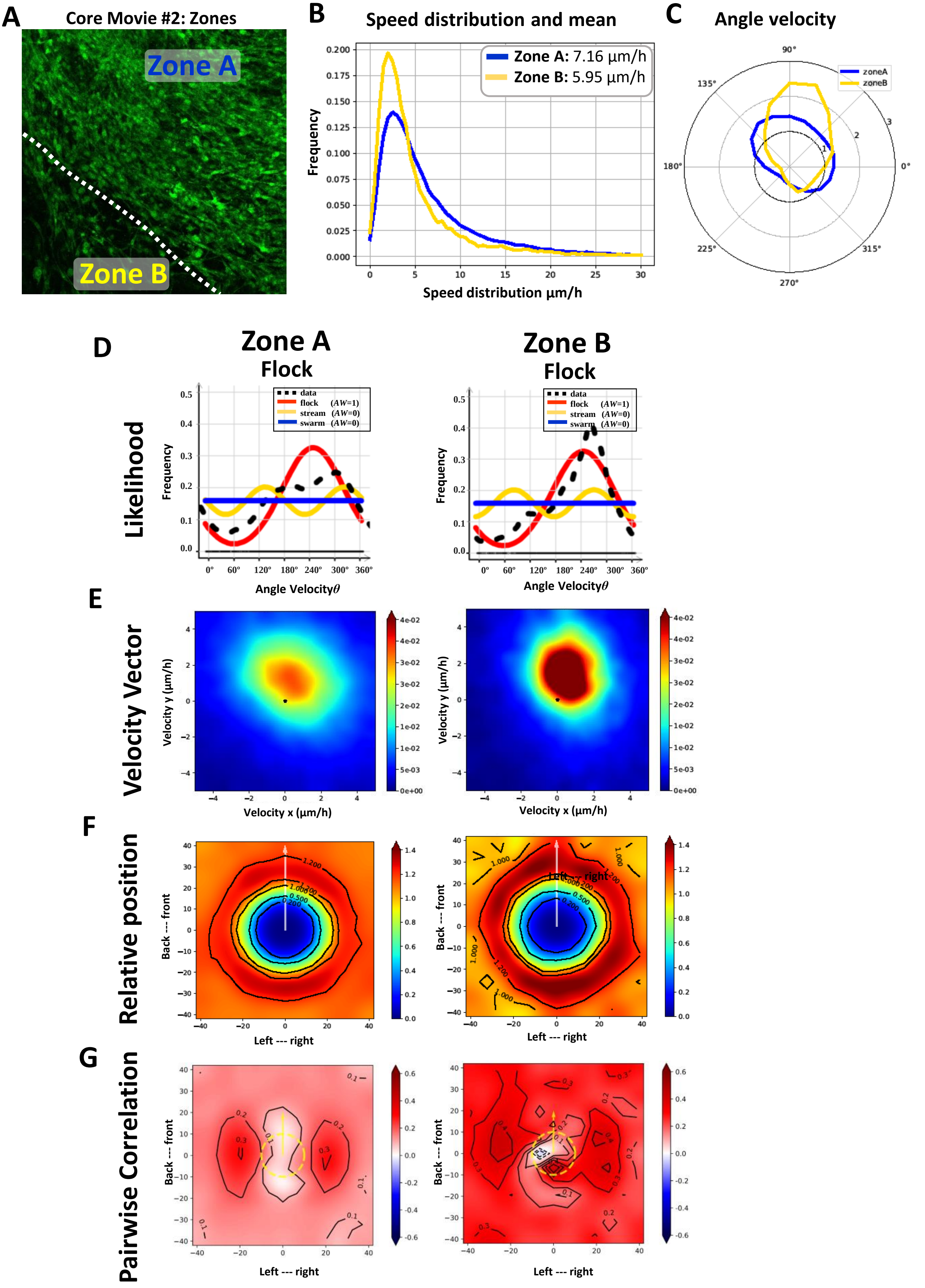

A
Core Movie #2: Zones
Zone A
Zone B
B
Speed distribution and mean
Zone A: 7.16 µm/h
Zone B: 5.95 µm/h
Frequency
Speed distribution µm/h
C
Angle velocity
zoneA
zoneB
D
Zone A
Flock
Zone B
Flock
Likelihood
data
flock (AW=1)
stream (AW=0)
swarm (AW=0)
Frequency
Angle Velocityθ
E
Velocity Vector
Velocity y (µm/h)
Velocity x (µm/h)
F
Relative position
Back --- front
Left --- right
G
Pairwise Correlation
Back --- front
Left --- right

**Fig. S18: Collective dynamic patterns observed by confocal imaging within the glioma tumor core: analysis of core movie #2. A)** Statistical analysis of different regions of the movie, Zone A and Zone B, illustrated on a representative time-lapse confocal still image. **B)** Speed distribution ($\mu m/hr$) in Zone A (blue) and B (yellow). Inner panel shows mean speed for each zone. **C)** Angle Velocity distribution analysis ($\vartheta$) performed by zones. **D)** Likelihood analysis histograms to classify dynamic pattern formation. Zone A: flock, Zone B: flock. AW: Akaiki Weight. AW=0 or AW=1. **E)** Heat map of the distribution of velocity vectors in each zone. **F)** Histogram plot showing relative position with nearby neighbors within each zone. X and Y axes are in µm. Scale bar shows frequency represented in colors. **G)** Histograms of pair-wise correlation with nearby neighbors for each zone. X and Y axes are in µm. Scale bar shows correlation represented as colors.

Fig. S19

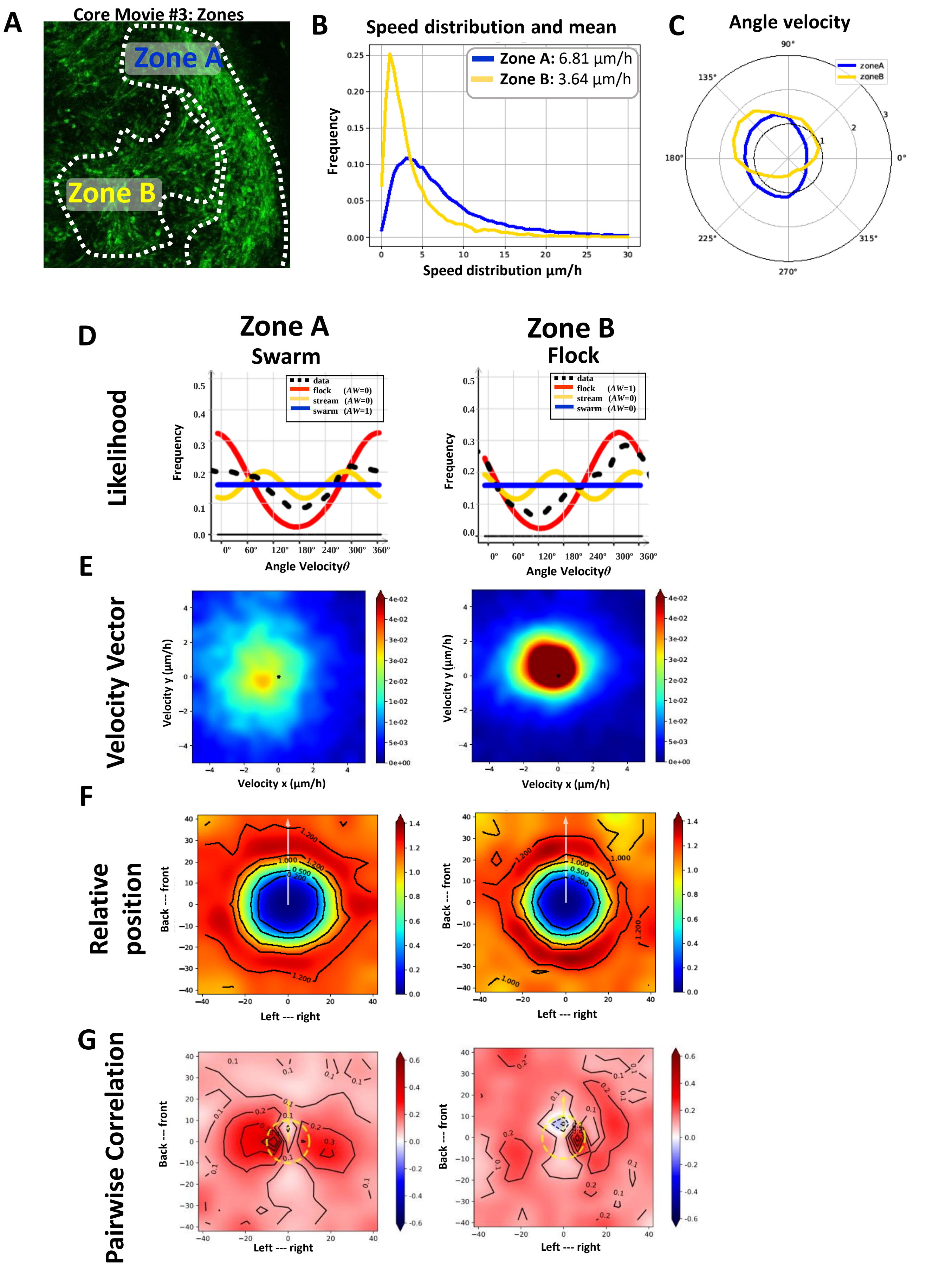
A
Core Movie #3: Zones
Zone A
Zone B
B
Speed distribution and mean
Zone A: 6.81 µm/h
Zone B: 3.64 µm/h
Frequency
Speed distribution µm/h
C
Angle velocity
zoneA
zoneB
D
Zone A
Swarm
Zone B
Flock
Likelihood
data
flock (AW=0)
stream (AW=0)
swarm (AW=1)
flock (AW=1)
swarm (AW=0)
Frequency
Angle Velocity$\theta$
E
Velocity Vector
Velocity y (µm/h)
Velocity x (µm/h)
F
Relative position
Back --- front
Left --- right
G
Pairwise Correlation
Back --- front
Left --- right

**Fig. S19: Collective dynamic patterns observed by confocal imaging within the glioma tumor core: analysis of core movie #3. A)** Statistical analysis of different regions of the movie, Zone A and Zone B, illustrated on a representative time-lapse confocal still image. **B)** Speed distribution ($\mu m/hr$) in Zone A (blue) and B (yellow). Inner panel shows mean of the speed for each zone. **C)** Angle Velocity distribution analysis ($\vartheta$) performed by zones. **D)** Likelihood analysis histograms to classify dynamic pattern formation. Zone A: flock, Zone B: flock. AW: 0 or AW:1. **E)** Heat map plot of the distribution of the velocity vector in each zone. **F)** Histogram plot showing interposition with nearby neighbor for each zone. X and Y axes are in µm. Scale bar shows frequency represented in colors. **G)** Histograms of pair-wise correlation with nearby neighbors for each zone. X and Y axes are in µm. Scale bar shows correlation represented in colors.

**Fig. S20**

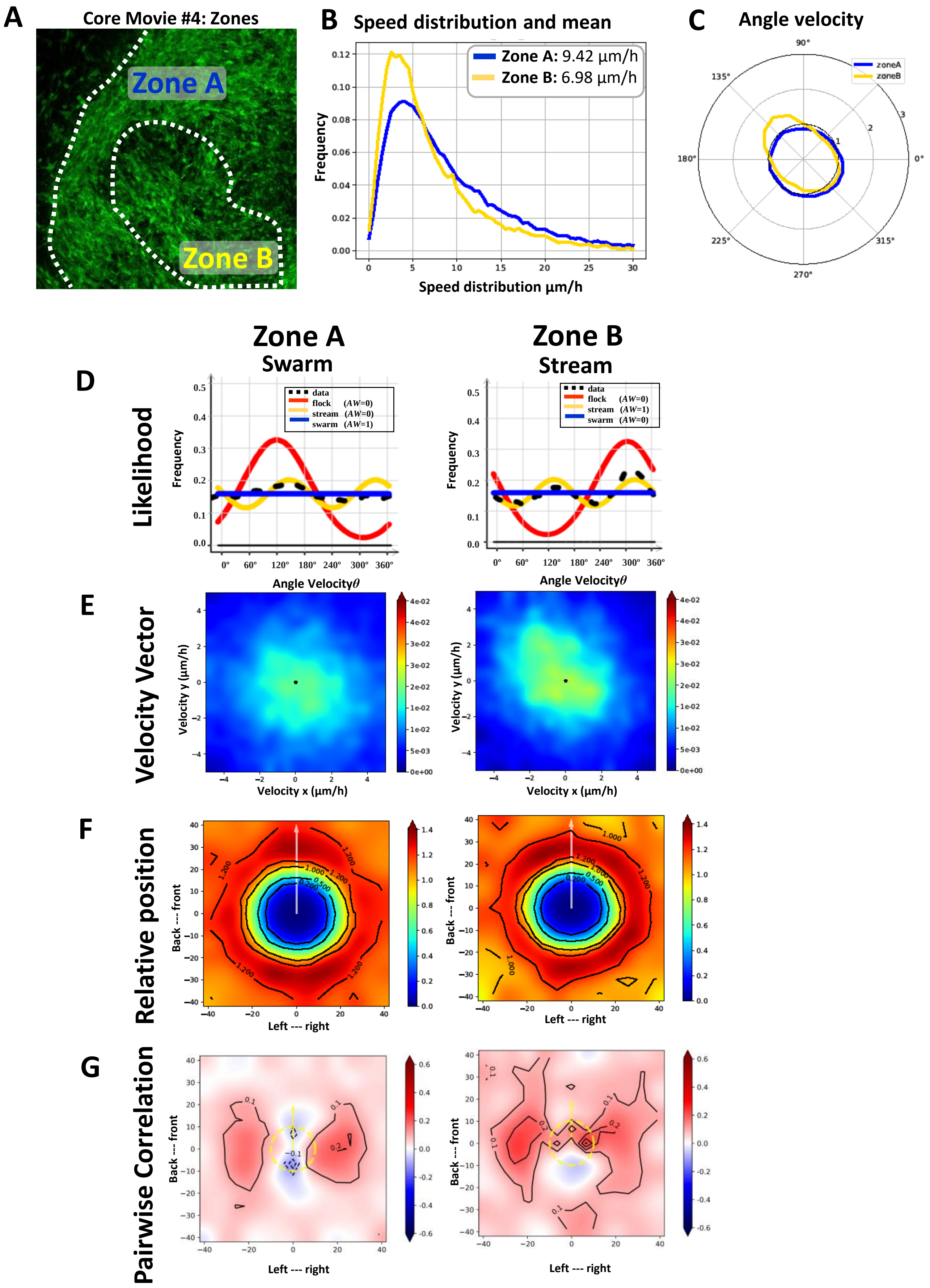
A
Core Movie #4: Zones
Zone A
Zone B
B
Speed distribution and mean
Zone A: 9.42 µm/h
Zone B: 6.98 µm/h
Frequency
Speed distribution µm/h
C
Angle velocity
zoneA
zoneB
Zone A
Swarm
Zone B
Stream
D
Likelihood
data
flock (AW=0)
stream (AW=0)
swarm (AW=1)
flock (AW=0)
stream (AW=1)
swarm (AW=0)
Frequency
Angle Velocity θ
E
Velocity Vector
Velocity y (µm/h)
Velocity x (µm/h)
F
Relative position
Back --- front
Left --- right
G
Pairwise Correlation
Back --- front
Left --- right

**Fig. S20: Collective dynamic patterns observed by confocal imaging within the glioma tumor core: analysis of core movie #4. A)** Statistical analysis of different regions of the movie, Zone A and Zone B, illustrated on a representative time-lapse confocal still image. **B)** Speed distribution ($\mu m/hr$) in Zone A (blue) and B (yellow). Inner panel shows mean of the speed for each zone. **C)** Angle Velocity distribution analysis ($\vartheta$) performed by zones. **D)** Likelihood analysis histograms to classify dynamic pattern formation. Zone A: swarm, Zone B: Stream. AW: 0 or AW:1. **E)** Heat map plot of the distribution of the velocity vector in each zone. **F)** Histogram plot showing interposition with nearby neighbor for each zone. X and Y axes are in µm. Scale bar shows frequency represented in colors. **G)** Histograms of pair-wise correlation with nearby neighbors for each zone. X and Y axes are in µm. Scale bar shows correlation represented in colors.

Fig. S21

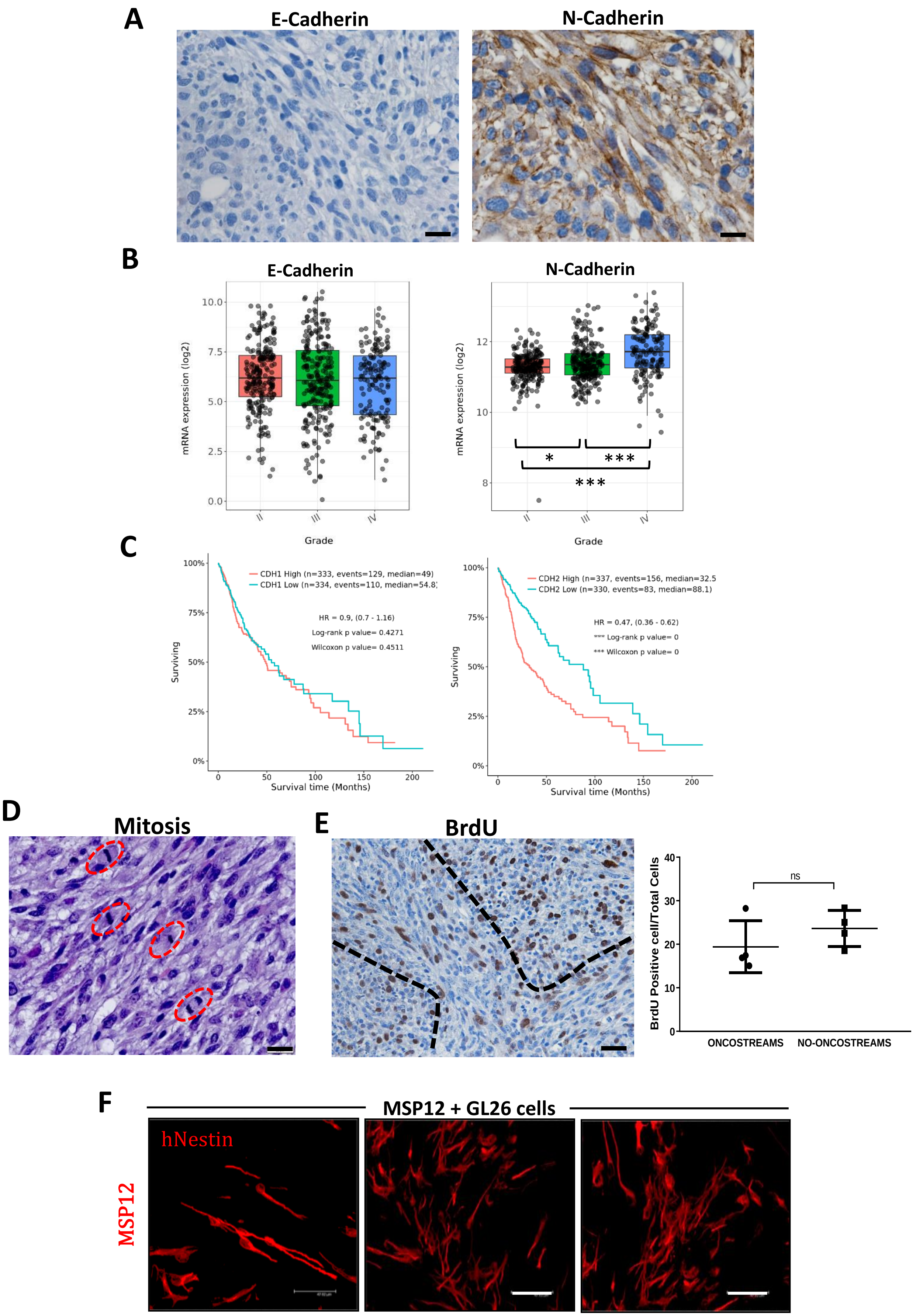

A
E-Cadherin
N-Cadherin
B
E-Cadherin
N-Cadherin
mRNA expression (log2)
10.0
7.5
5.0
2.5
0.0
12
10
8
*
***
***
II
III
IV
Grade
C
CDH1 High (n=333, events=129, median=49)
CDH1 Low (n=334, events=110, median=54.8)
HR = 0.9, (0.7 - 1.16)
Log-rank p value= 0.4271
Wilcoxon p value= 0.4511
CDH2 High (n=337, events=156, median=32.5
CDH2 Low (n=330, events=83, median=88.1)
HR = 0.47, (0.36 - 0.62)
*** Log-rank p value= 0
*** Wilcoxon p value= 0
Surviving
100%
75%
50%
25%
0%
0
50
100
150
200
Survival time (Months)
D
Mitosis
E
BrdU
BrdU Positive cell/Total Cells
40
30
20
10
0
ns
ONCOSTREAMS
NO-ONCOSTREAMS
F
MSP12 + GL26 cells
hNestin
MSP12

**Fig. S21: Characterization of Oncostream' s cellular functions. A**) Immunohistochemistry analysis of E-cadherin and N-cadherin expression within NPA gliomas. Gliomas are negative for E-cadherin and positive for N-cadherin both within and outside oncostream structures. **B)** mRNA expression analysis of N-cadherin and E-cadherin comparing glioma tumors of different grade using the TCGA- GBMLGG database. Graph shows the $log_2$ mRNA expression levels. Statistical significance was given within the corresponding databases (Tukey's Honest Significant Difference (HSD). **C)** Kaplan-Meier survival curves of glioma patients comparing N-cadherin and E-cadherin high vs low expression in glioma tumors (TCGA-GBMLGG database). All these data was analyzed from the Gliovis website (http://gliovis.bioinfo.cnio.es). **D)** Analysis of glioma cell mitosis within oncostreams in H&E stained sections. Mitosis is orientated in the same orientation of oncostreams. Red lines indicate mitosis. **E)** Proliferation analysis comparing oncostream (dotted lines) with no-oncostream areas. Positive BrDu cells were counted by Image-J software. Scale bars: 50 µm. BrDu positive cells per total cells in the visual field were counted; n=4. Ten fields of each section were selected at random. Error bars represent ± SEM; paired t-test. **F)** Immunofluorescence images of human-nestin (red) highlighting MSP12 cells. MSP-12 cells were implanted with GL26-citrine cells. MSP12 cells adopt a bipolar structure only when aligned to GL26-citrine cells. Scale bar: 47.62 µm. This image complement Fig. 7K.

Fig. S22

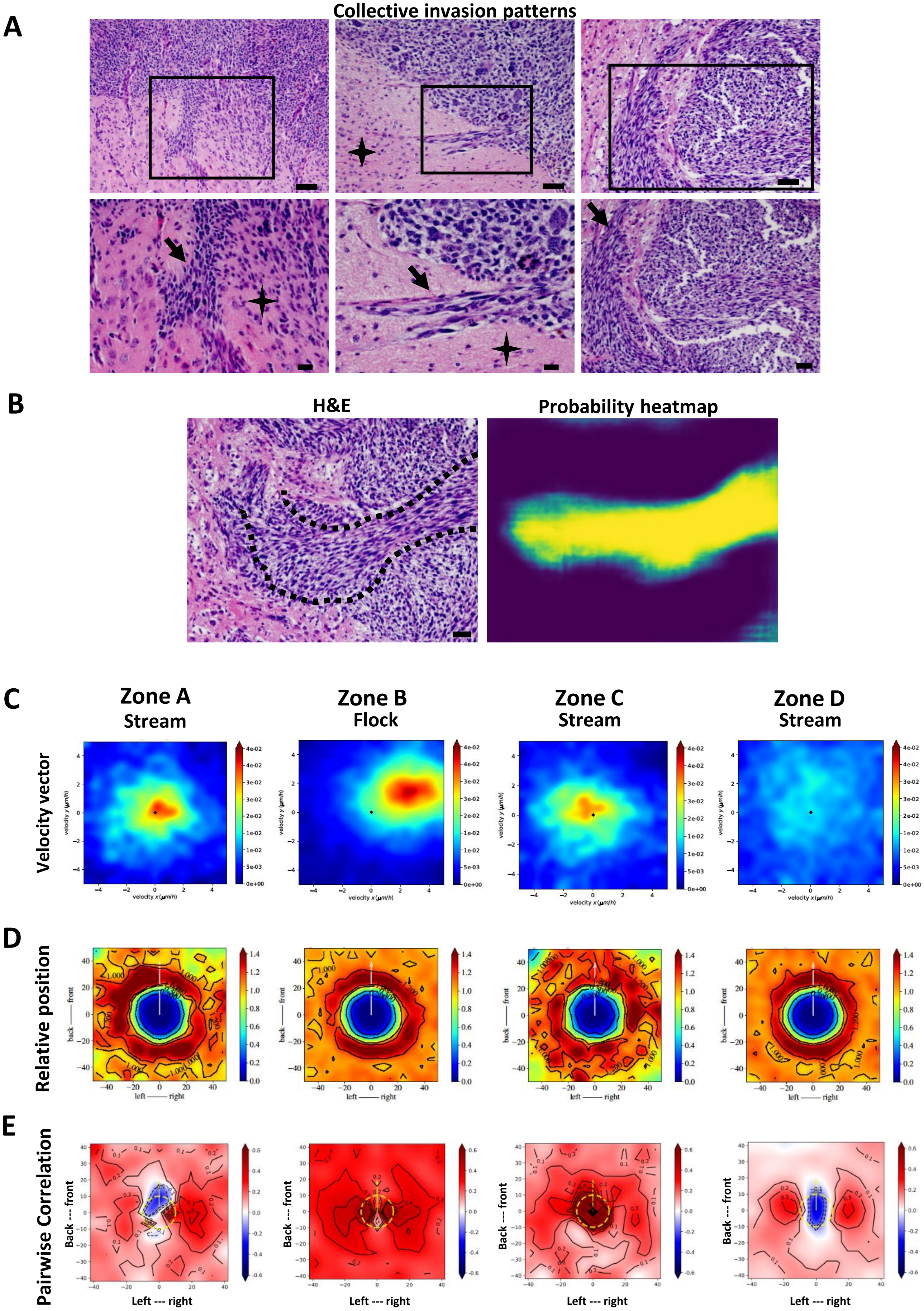

A
Collective invasion patterns
B
H&E
Probability heatmap
C
Zone A
Stream
Zone B
Flock
Zone C
Stream
Zone D
Stream
Velocity vector
D
Relative position
E
Pairwise Correlation
Left --- right
Back --- front

**Fig. S22: Invasion analysis using confocal time-lapse imaging on the tumor border: Movie #5. A)** Representative images of oncostream invasion (top row, black boxes) from H&E stained sections of GEMM of glioma. Arrows indicate oncostream collective invasion. Areas highlighted by black boxes are shown at higher magnification in the lower row. Stars indicate sites of single cell invasion. Scale bars: 50 µm (top) and 20 µm (bottom). **B)** Representative images of oncostream invasion (dotted line indicates an oncostream at the tumor border) on H&E stained sections of NPA gliomas (left), and its detection by deep learning analysis (right). Scale bar=20 µm. **C)** Heatmap plot of the distribution of velocity vectors in zones A, B, C and D of border movie #5 (shown in **Fig. 5**). **D)** Histogram of relative cell positions in relation to nearby neighbors. For each cell, $x_{i,}$ we estimate the probability to find another cell, $x_{j,}$ nearby. Probability scale bars are represented by colors. Axes shows cells position left-to-right and front-to-back in µm. **E)** Pairwise correlation of motion with nearby neighbors. Scale bars of probability are represented by colors. Axes indicate cell position and distance in µm from neighboring cells. Dotted yellow line indicates the location of $x_i$.

Fig. S23

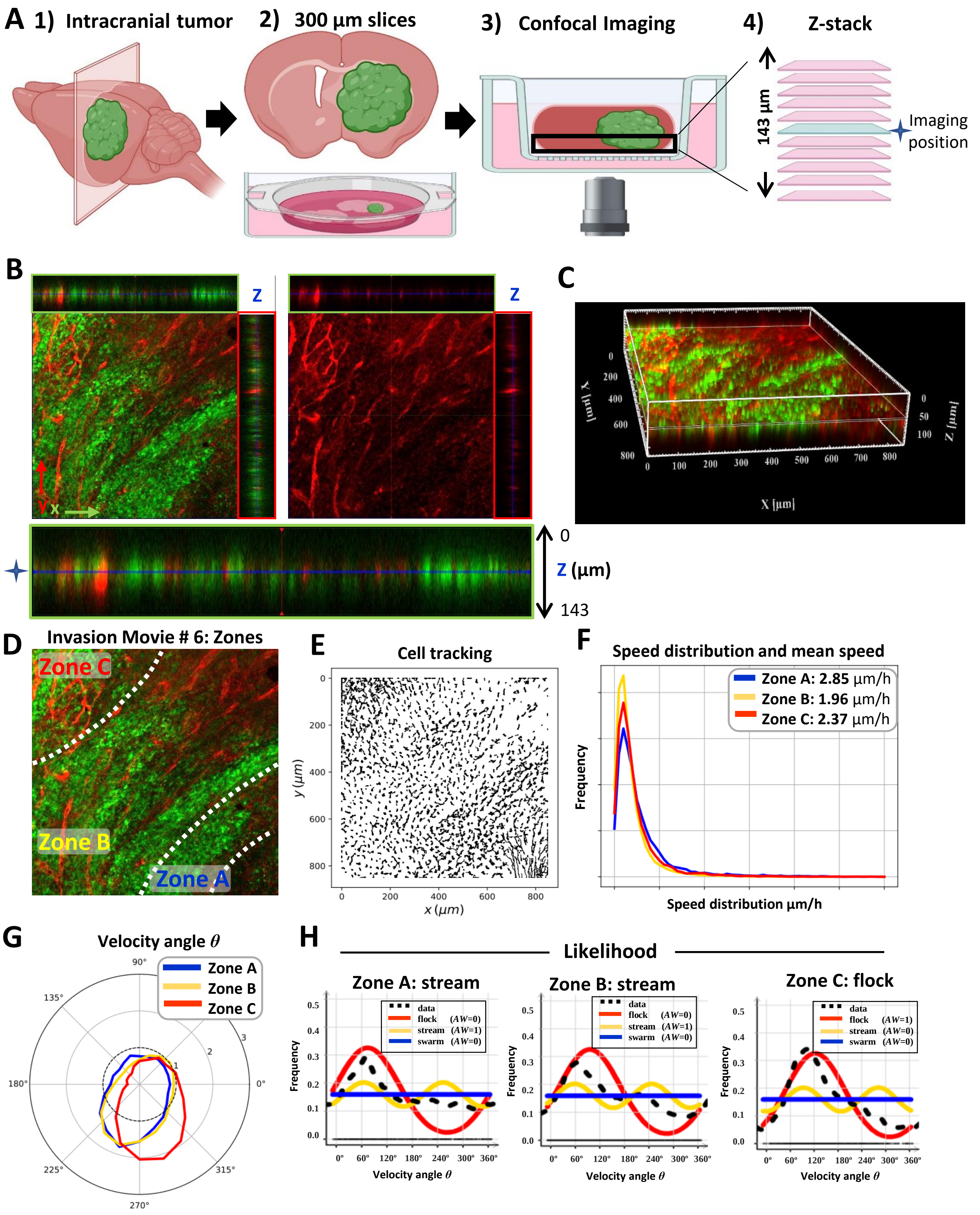
A 1) Intracranial tumor
2) 300 µm slices
3) Confocal Imaging
4) Z-stack
143 µm
Imaging position
B
Z
Z
0
Z (µm)
143
C
X [µm]
Z [µm]
D Invasion Movie # 6: Zones
Zone C
Zone B
Zone A
E
Cell tracking
y (µm)
x (µm)
F
Speed distribution and mean speed
Zone A: 2.85 µm/h
Zone B: 1.96 µm/h
Zone C: 2.37 µm/h
Frequency
Speed distribution µm/h
G
Velocity angle θ
Zone A
Zone B
Zone C
H
Likelihood
Zone A: stream
Zone B: stream
Zone C: flock
data
flock (AW=0)
stream (AW=1)
swarm (AW=0)
flock (AW=1)
stream (AW=0)
Frequency
Velocity angle θ

**Fig. S23**

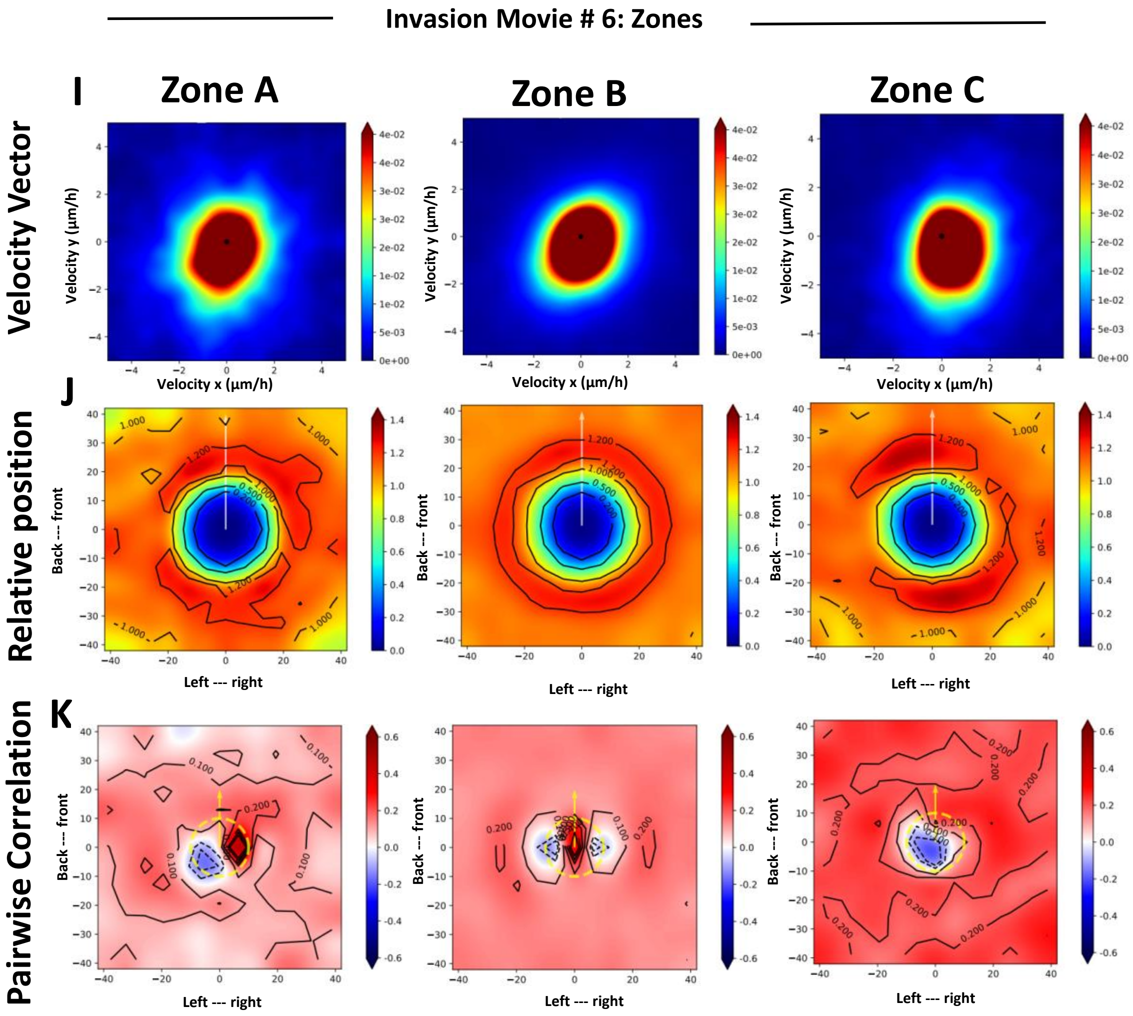
Invasion Movie # 6: Zones
I
Zone A
Zone B
Zone C
Velocity Vector
Velocity y (μm/h)
Velocity x (μm/h)
J
Relative position
Back --- front
Left --- right
K
Pairwise Correlation
Back --- front
Left --- right

**Fig. S23. Brain tumor explant model setup and invasion analysis using time lapse-confocal imaging: Movie #6**

**A)** Schematic representation of the experimental setup and site of imaging. 1) NPA-GFP+ cells were intracranially implanted, and brains from tdTomato mice bearing GFP+ tumors were isolated 19 days post-tumor implantation. 2) These brains were sliced into 300 µm thick coronal sections and transferred to a laminin-coated cell culture Insert. 3) Cell culture insets containing brain tumor slices were placed in the incubator chamber of an inverted Zeiss LSM880 laser scanning confocal microscope at 37 °C with a 5% CO2 atmosphere. The black rectangle represents the imaging area that extends from the bottom to ~143 µm; not being able to image further into the brain slice. 4) A high-resolution z-stack was obtained (dimensions of x=850.19, y=850.19, z=143.81). The location for time-lapse imaging was selected to fall at the middle of the z-stack (~70 µm) to avoid imaging at the bottom of the explant. Red fluorescent protein: normal brain parenchyma. Green fluorescent protein: tumor cells. **B)** Orthogonal projections from z-stacks revealed the cross-sectional aspects of the tumor volume and the position of time-lapse confocal imaging of Movie #6, an NPA glioma. Green rectangles (on top) show the profile of the x-z plane. Red rectangles (on the right) show the profile of the y-z plane. The left image illustrates tumor cells (in green) and brain parenchyma (in red); the right image shows only the brain parenchyma. The larger green rectangle at the bottom of **(Fig. S23B)** illustrates the orthogonal z-stack projection of the x-z plane (from 0 to 143 µm) and the exact plane of imaging, in the middle of z plane (indicated by a star and a blue line). **C)** A high-resolution 3D z-stack spanning up to 143 µm depth (starting at the bottom of the brain slice) was acquired on a confocal microscope, imported into the Imaris viewer, and used to reconstruct this 3D image. 30 x-y frames from the brain's surface were taken at a depth increment of 4.96 µm at a resolution of 1024x1024 pixels. XYZ axes of the 3D image are shown in white (850x850x143 µm), and the yellow line shows the exact imaging plane for time-lapse data acquisition *in vivo* (at ~70 µm depth). Red fluorescent protein: normal brain parenchyma. Green fluorescent protein: tumor cells. **D)** Representative time-lapse scanning confocal image of an NPA glioma growing at the tumor border, and showing Zones A, B, C. **E)** Tracking analysis of individual cell paths using the TrackMate plugin from Image-J. **F)** Histogram of speed distribution and mean speed ($\mu m/h$) of Zones A, B and C. **G)** Angle Velocity distribution analysis ($\vartheta$) performed by zones. The plot shows the overall direction and magnitude of cell movement. **H)** Likelihood analysis of the dynamic patterns at the tumor border. The frequency distribution *ρ flock* (red), *ρ stream* (yellow) and *ρ swarm* (blue) are shown. The estimation of the black line (data) uses a *non-parametric* estimation. AW: 0 or AW:1. **I)** Heat map plot of the distribution of the velocity vectors in each zone. **J)** Histogram plot showing the relative position with nearby neighbors for each zone; *x* and *y* axes are in µm. Bars to the right of each figure indicate probability values in color. **K)** Histograms of pairwise correlations with nearby neighbors for each zone; *x* and *y* axes are in µm. Bars to the right of each figure indicate correlation values in colors.

Fig. S24

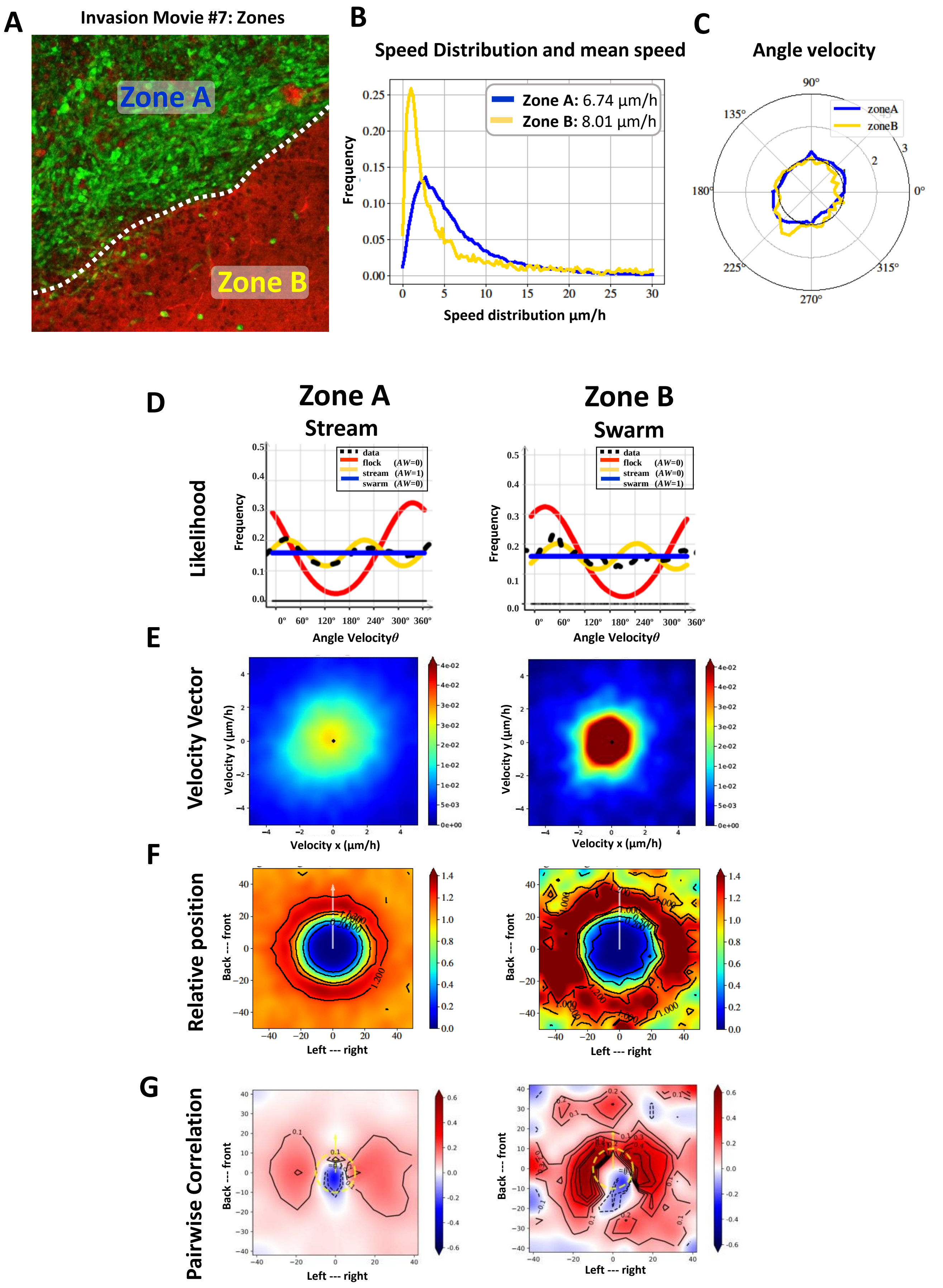
A
Invasion Movie #7: Zones
Zone A
Zone B
B
Speed Distribution and mean speed
Zone A: 6.74 µm/h
Zone B: 8.01 µm/h
Frequency
Speed distribution µm/h
C
Angle velocity
zoneA
zoneB
D
Zone A
Stream
Zone B
Swarm
Likelihood
data
flock (AW=0)
stream (AW=1)
swarm (AW=0)
flock (AW=0)
stream (AW=0)
swarm (AW=1)
Frequency
Angle Velocityθ
E
Velocity Vector
Velocity y (µm/h)
Velocity x (µm/h)
F
Relative position
Back --- front
Left --- right
G
Pairwise Correlation
Back --- front
Left --- right

**Fig. S24: Invasion analysis using confocal time-lapse imaging on the tumor border: Movie #7) A)** Labeling of different regions of the tumor border for movie #6. Representative time-lapse confocal images subdivided into dynamic Zones A and B. **B)** Speed distribution ($\mu m/hr$) in Zone A (blue) and B (yellow). Inner panel shows mean speed for each zone. **C)** Angle Velocity distribution analysis ($\vartheta$) performed by zones. **D)** Likelihood analysis histograms classify dynamic motion patterns. Zone A: stream, Zone B: swarm. AW: 0 or AW:1. **E)** Heat map plot of the distribution of the velocity vectors in each zone. **F)** Histogram plot showing the relative position with nearby neighbors for each zone. *x* and *y* axes are in µm. Bars to the right of each figure indicate probability values in color. **G)** Histograms of pairwise correlations with nearby neighbors for each zone. *x* and *y* axes are in µm. Bars to the right of each figure indicate correlation values in colors.

**Fig. S25**

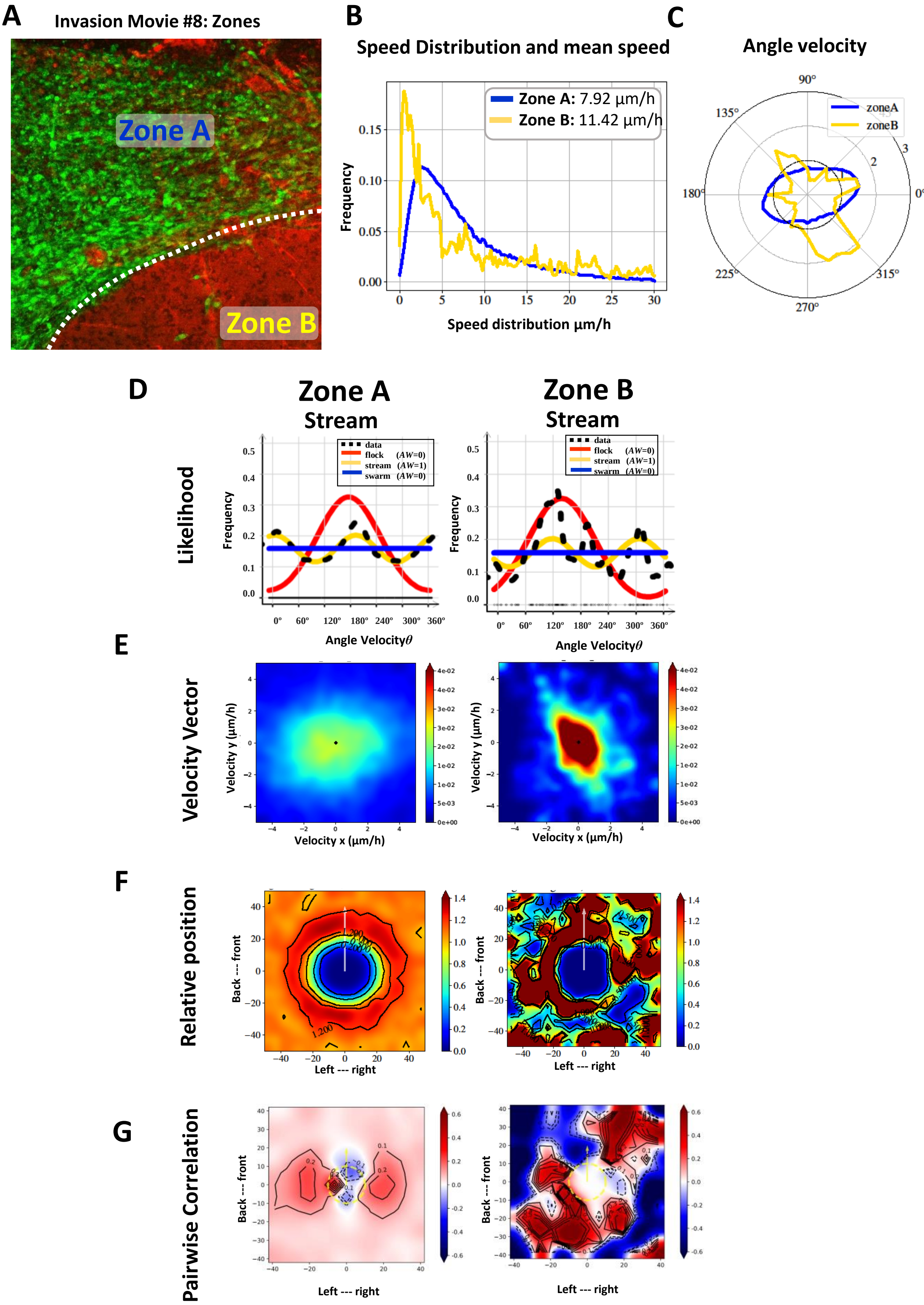
A
Invasion Movie #8: Zones
Zone A
Zone B
B
Speed Distribution and mean speed
Zone A: 7.92 µm/h
Zone B: 11.42 µm/h
Frequency
Speed distribution µm/h
C
Angle velocity
zoneA
zoneB
D
Zone A
Stream
Zone B
Stream
Likelihood
data
flock (AW=0)
stream (AW=1)
swarm (AW=0)
Frequency
Angle Velocityθ
E
Velocity Vector
Velocity y (µm/h)
Velocity x (µm/h)
F
Relative position
Back --- front
Left --- right
G
Pairwise Correlation
Back --- front
Left --- right

**Fig. S25: Invasion analysis using confocal time-lapse imaging of the tumor border: Movie #8. A)** Labeling of different regions of the tumor border for movie #7. Representative time-lapse confocal images subdivided into dynamic Zones A and B. **B)** Speed distribution ($\mu m/hr$) in Zone A (blue) and B (yellow). Inner panel shows mean speed for each zone. **C)** Angle Velocity distribution analysis ($\vartheta$) performed by zones. **D)** Likelihood analysis histograms classify dynamic motion patterns. Zone A: stream, Zone B: stream. Akaiki Weight; AW: 0 or AW:1. **E)** Heat map plot of the distribution of the velocity vectors in each zone. **F)** Histogram plot showing the relative position with nearby neighbors for each zone. *x* and *y* axes are in µm. Bars to the right of each figure indicate probability values in color. **G)** Histograms of pairwise correlations with nearby neighbors for each zone. *x* and *y* axes are in µm. Bars to the right of each figure indicate correlation values in colors.

Fig. S26

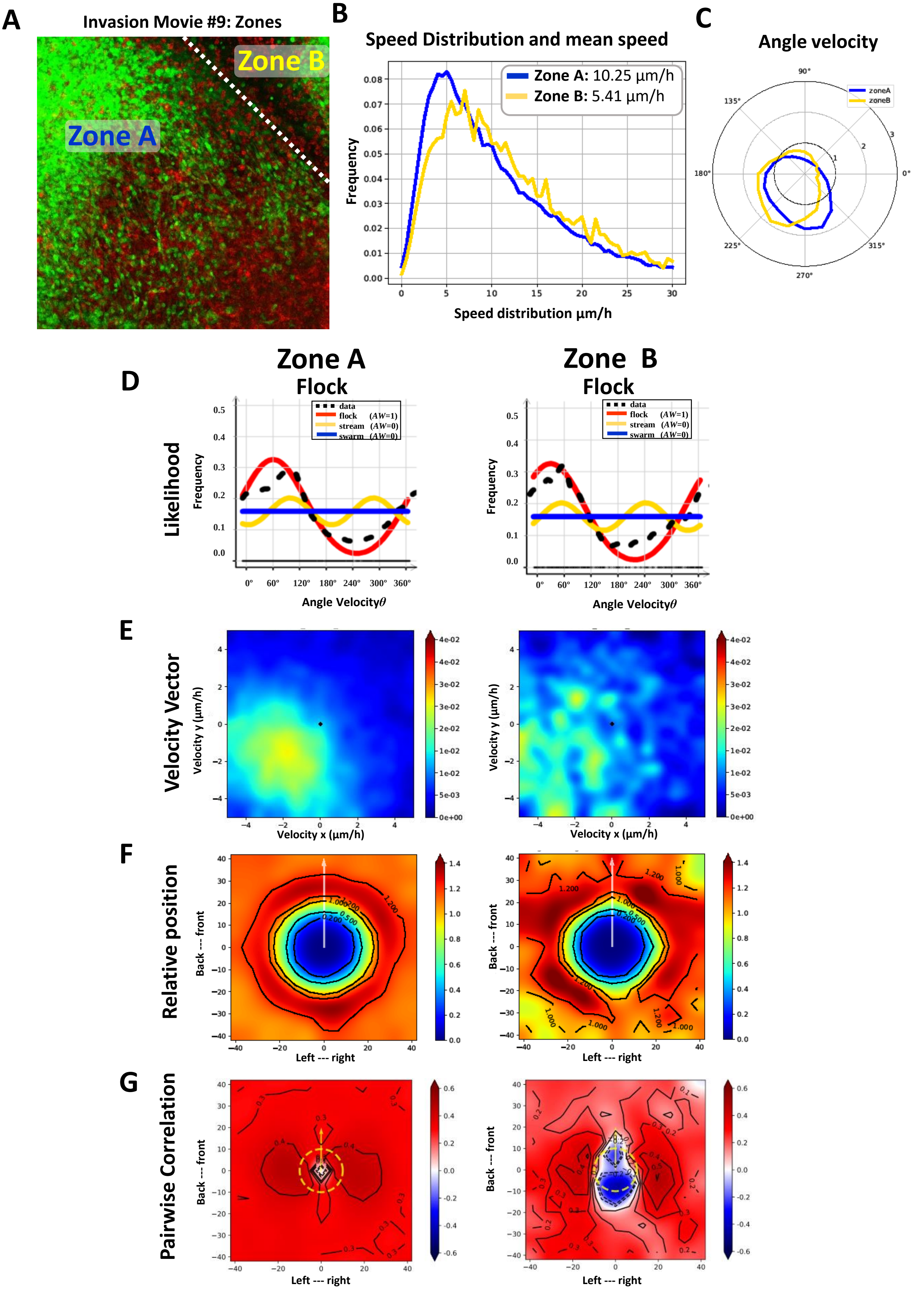
A
Invasion Movie #9: Zones
Zone A
Zone B
B
Speed Distribution and mean speed
Zone A: 10.25 µm/h
Zone B: 5.41 µm/h
Frequency
Speed distribution µm/h
C
Angle velocity
zoneA
zoneB
D
Zone A
Flock
Zone B
Flock
Likelihood
Frequency
data
flock (AW=1)
stream (AW=0)
swarm (AW=0)
Angle Velocityθ
E
Velocity Vector
Velocity y (µm/h)
Velocity x (µm/h)
F
Relative position
Back --- front
Left --- right
G
Pairwise Correlation
Back --- front
Left --- right

**Fig. S26: Invasion analysis using confocal time-lapse imaging on the tumor border: Movie #9. A)** Labeling of different regions of the tumor border for movie #8. Representative time-lapse confocal images subdivided into dynamic Zones A and B. **B)** Speed distribution ($\mu m/h$) in Zone A (blue) and B (yellow). Inner panel shows mean speed for each zone. **C)** Angle Velocity distribution analysis ($\vartheta$) performed by zones. **D)** Likelihood analysis histograms to classify dynamic motion patterns. Zone A: flock, Zone B: flock. AW: 0 or AW:1. **E)** Heat map plot of the distribution of the velocity vectors in each zone. **F)** Histogram plot showing the relative position with nearby neighbors for each zone. *x* and *y* axes are in µm. Bars to the right of each figure indicate probability values in color. **G)** Histograms of pairwise correlations with nearby neighbors for each zone. *x* and *y* axes are in µm. Bars to the right of each figure indicate correlation values in colors.

#### Fig. S27

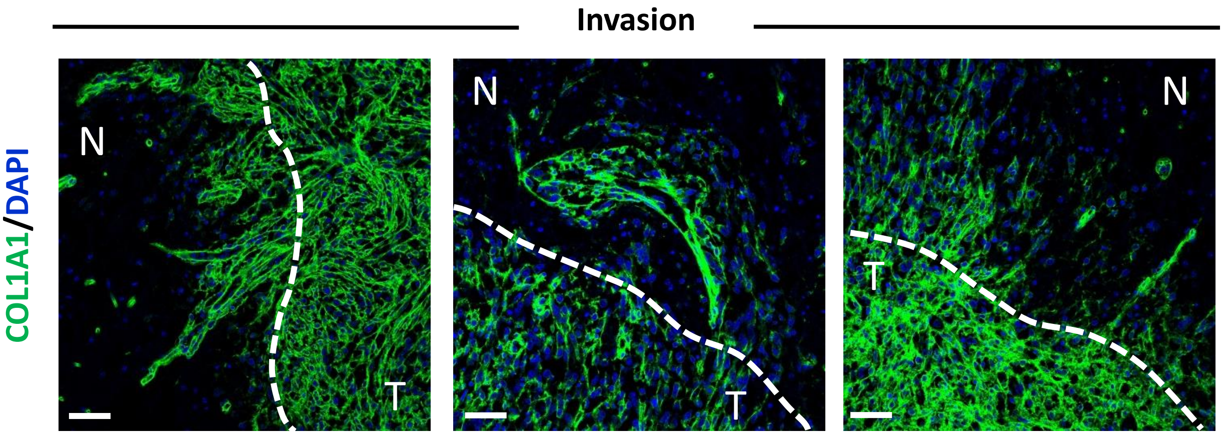


**Fig. S27. Role of COL1A1 fibers in oncostreams invasion of the normal brain.** Representative images of immunofluorescence analysis of COL1A1 expression in a genetically engineered mouse glioma, NPA (IDH1-WT). Images show aligned fibers of collagen1A1 along multicellular fascicles of glioma cells invading the normal brain. We propose that Collagen fibers function as scaffolds for collective tumoral cell migration. COL1A1 expression in green (Alexa 488) and nuclei in blue (DAPI). Images (20x, 1240 x 1240 pixels). White dotted line indicates tumor core border. Scale bar: 50 µm. T=Tumor. N=normal tissue.

**Fig. S28**

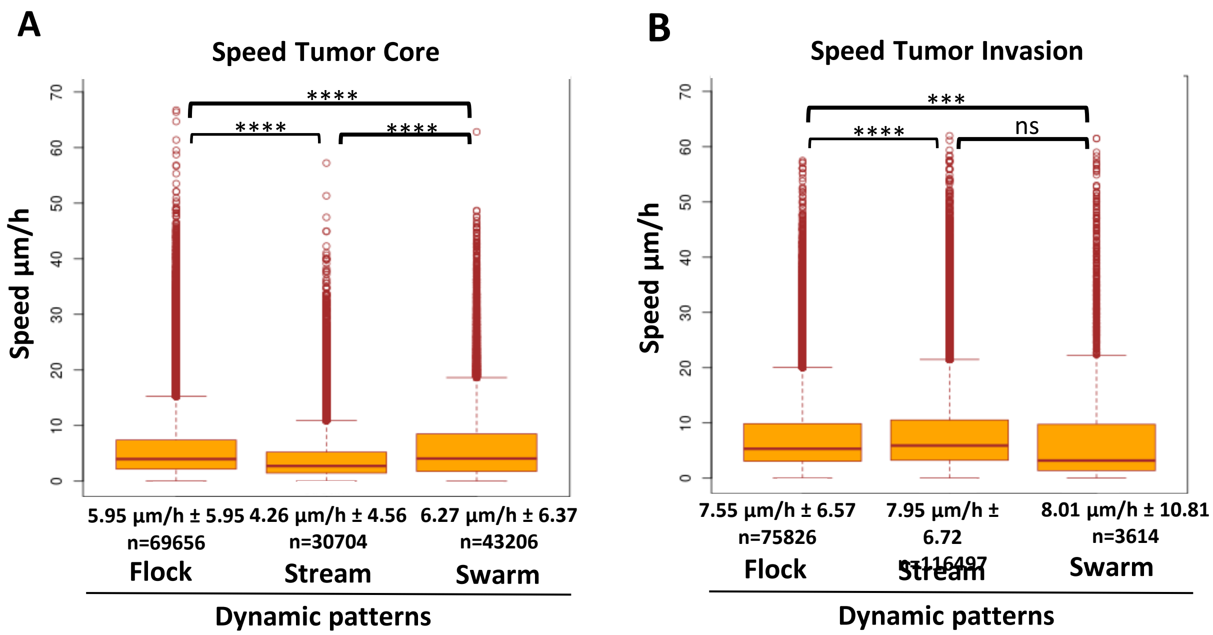


**Fig. S28: Speed analysis of cells in either flocks, streams, or swarms within the tumor core or the tumor border**. **A)** Box plots of cell speed (µm/hr) within flocks, streams, or swarms in the tumor core. Stream n=30704, Flock **n= 69656**, Swarm n=43206. Mean ±SEM are shown; One-way ANOVA, ***p<0.001, ***p<0.0001. **B)** Box plot of speed of cells (µm/hr) within flocks, streams, or swarms within the tumor border. Stream n=116497, Flock **n=75826**, Swarm n=3614. Mean ±SEM are shown; One-way ANOVA, ***p<0.001, ***p<0.0001.

**Fig. S29**

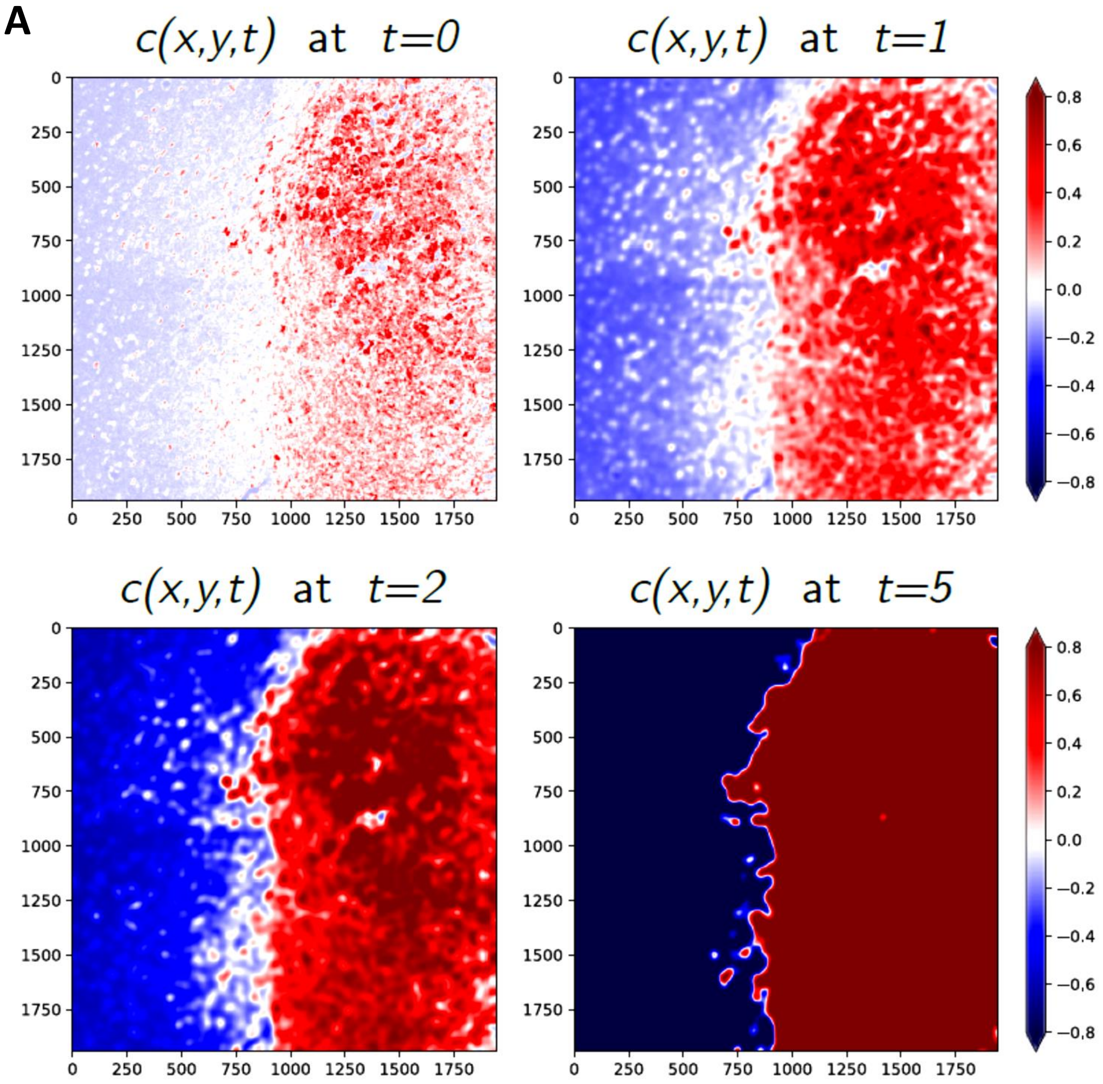


**Fig. S29: Analysis of the tumor borders. A)**Two-phase segmentation of an image using Allen-Cahn equation (3.1) described in Supplementary material and Methods. Each pixel has to converge to either +1 or -1 depending on its neighboring pixels. As a result, the image becomes segregated into two regions representing the inside and the outside of the tumor.

#### Fig. S30

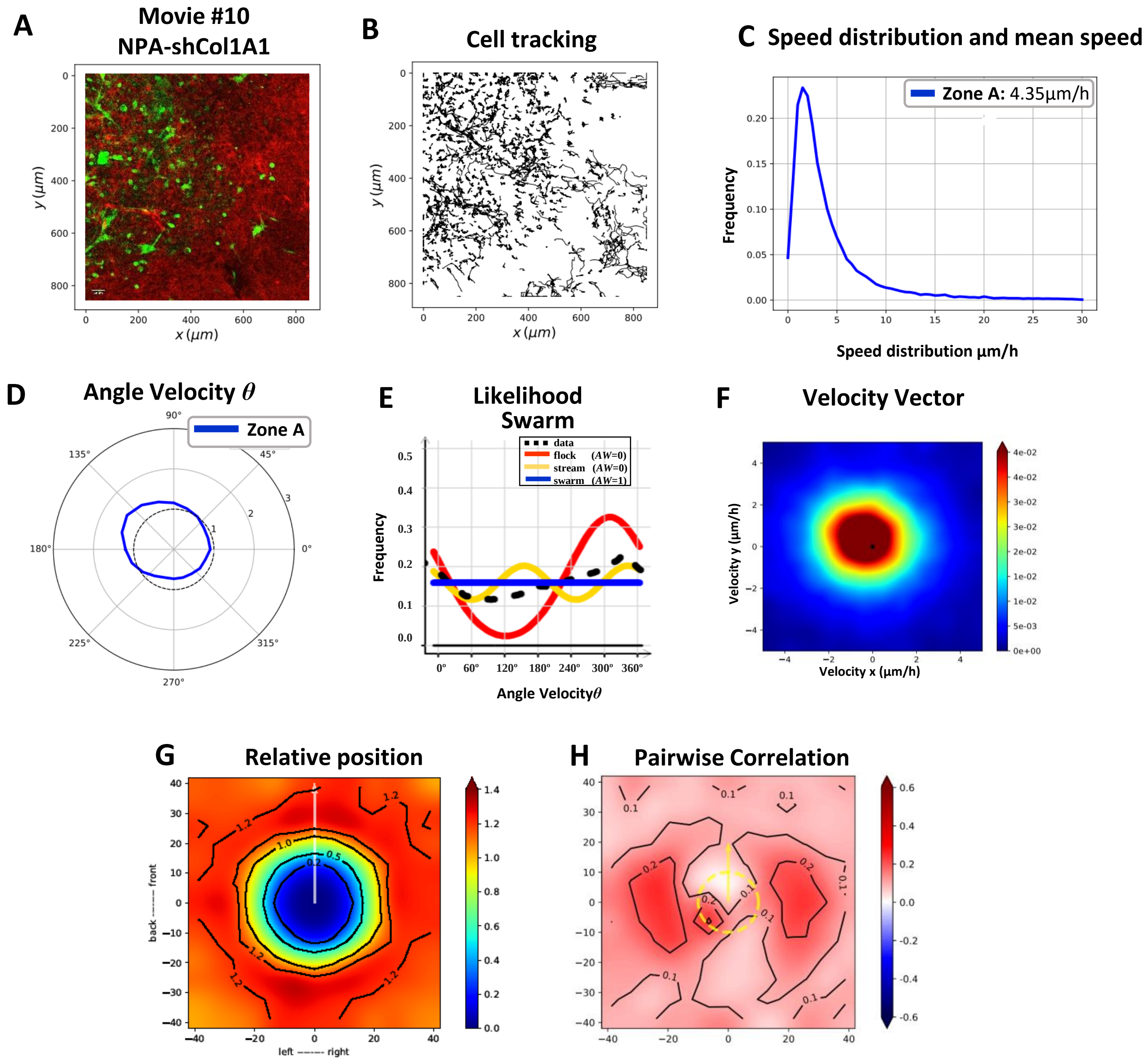


**Fig. S30: Migration analysis of an NPAshCol1A1 explant model using confocal time-lapse imaging; Movie # 10. A)** Representative time-lapse confocal images of the tumor border for movie #9. **B)** Tracking analysis of individual cell paths performed using the Track-Mate plugin from Image-J. **C)** Speed distribution and mean speed (*µm/h*). **D)** Angle Velocity distribution analysis (*ϑ*) performed. **E)** Likelihood analysis histogram to classify the dynamic motion pattern as a swarm. **F)** Heat map plot of the distribution of the velocity vectors. **G)** Histogram plot showing the relative position with nearby neighbors. *x* and *y* axes are in µm. Bars to the right of each figure indicate probability values in color. **H)** Histograms of pairwise correlations with nearby neighbors. *x* and *y* axes are in µm. Bars to the right of each figure indicate correlation values in colors.

**Fig. S31**

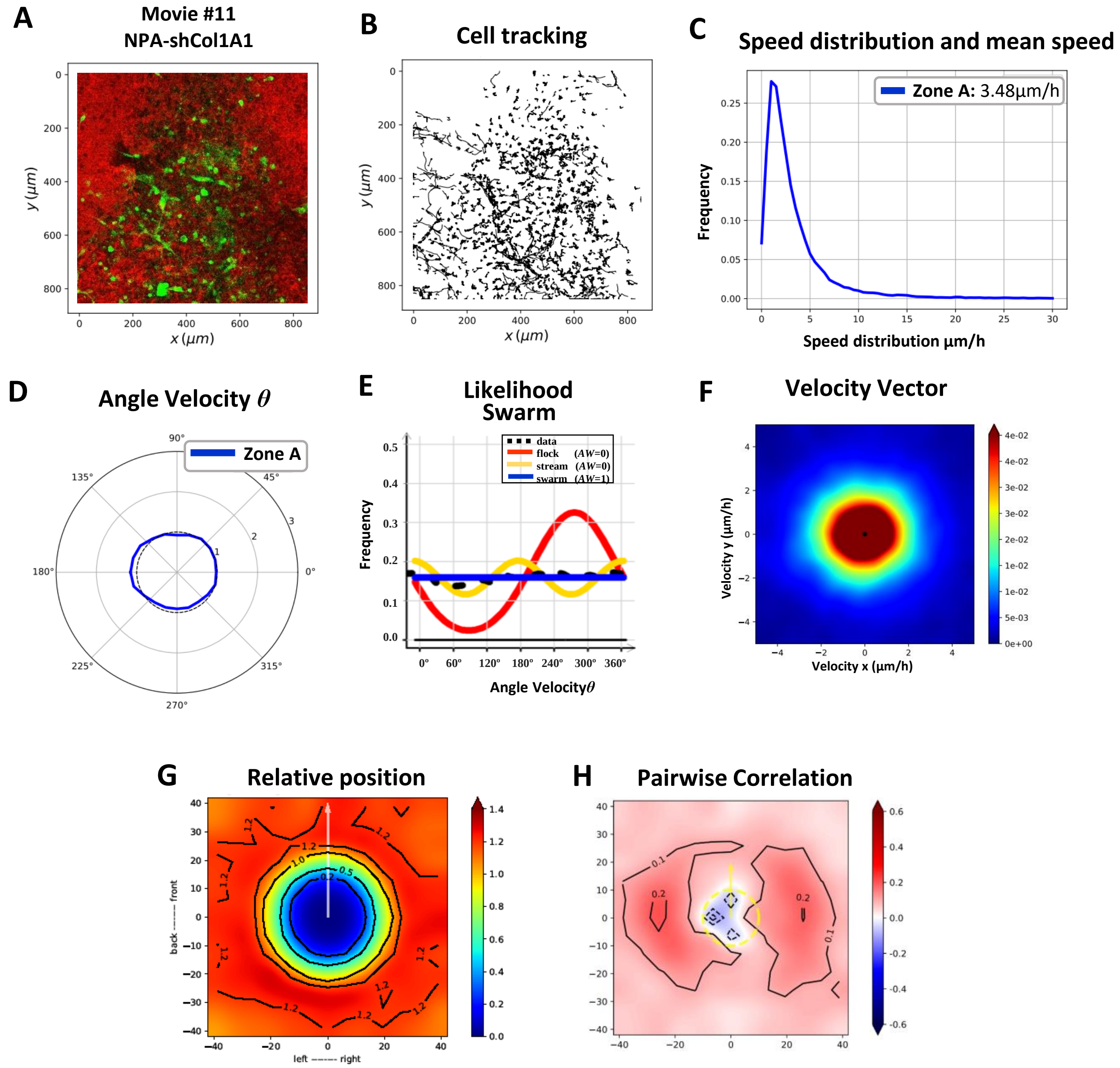


**Fig. S31: Migration analysis of an NPAshCol1A1 explant model using confocal time-lapse imaging; Movie # 11. A)** Representative time-lapse confocal images of the tumor border for movie #9. **B)** Tracking analysis of individual cell paths performed using the Track-Mate plugin from Image-J. **C)** Speed distribution and mean speed ($\mu m/h$). **D)** Angle Velocity distribution analysis ($\vartheta$) performed. **E)** Likelihood analysis histogram to classify the dynamic motion pattern as a swarm. **F)** Heat map plot of the distribution of the velocity vectors. **G)** Histogram plot showing the relative position with nearby neighbors. *x* and *y* axes are in µm. Bars to the right of each figure indicate probability values in color. **H)** Histograms of pairwise correlations with nearby neighbors, *x* and *y* axes are in µm. Bars to the right of each figure indicate correlation values in colors.

**Fig. S32**

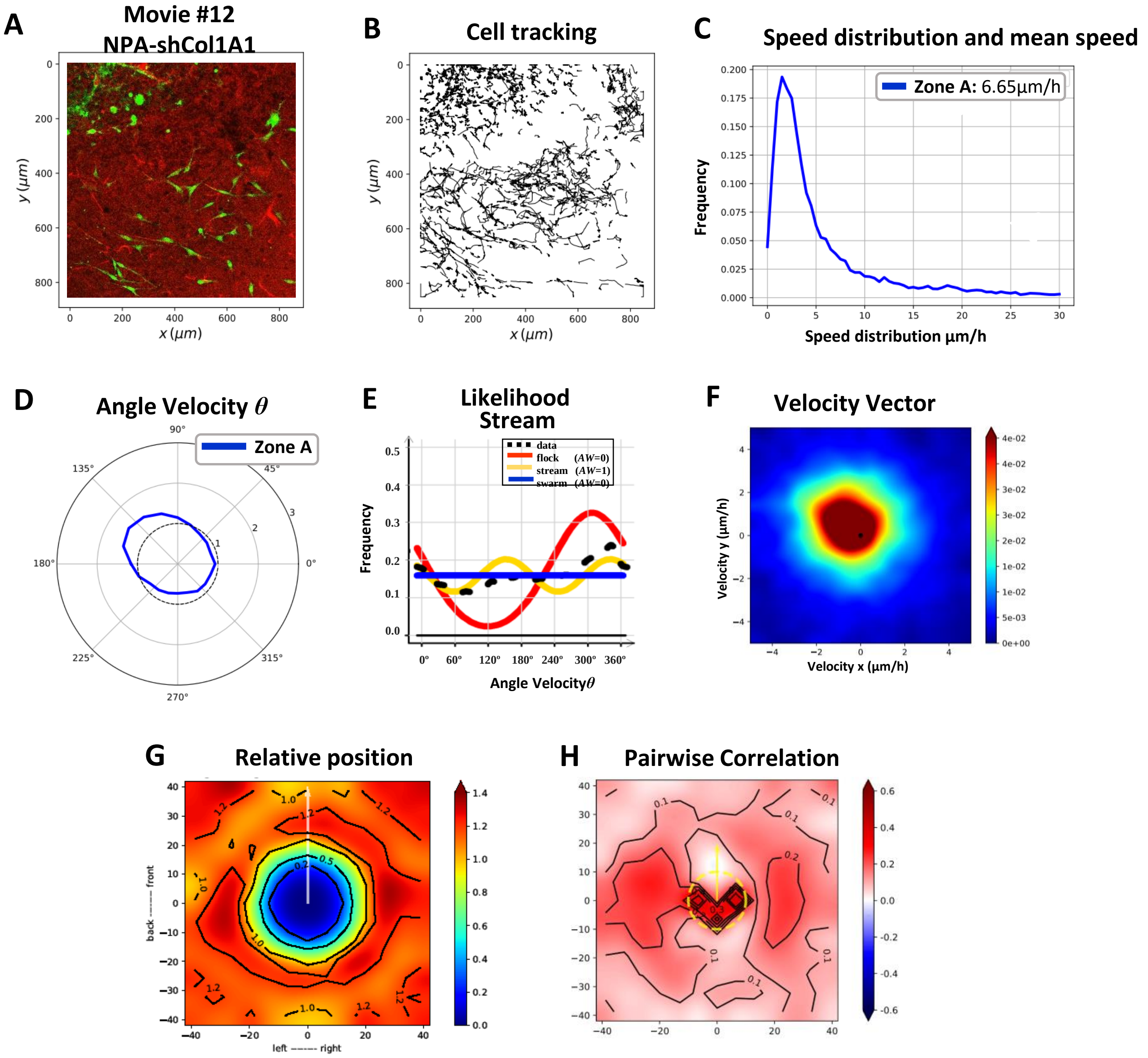


**Fig. S32: Migration analysis of an NPAshCol1A1 explant model using confocal time-lapse imaging; Movie # 12. A)** Representative time-lapse confocal images of the tumor border for movie #9. **B)** Tracking analysis of individual cell paths performed using the Track-Mate plugin from Image-J. **C)** Speed distribution and mean speed (*μm/h*). **D)** Angle Velocity distribution analysis (*ϑ*) performed. **E)** Likelihood analysis histogram to classify the dynamic motion pattern as a swarm. **F)** Heat map plot of the distribution of the velocity vectors. **G)** Histogram plot showing the relative position with nearby neighbors. *x* and *y* axes are in μm. Bars to the right of each figure indicate probability values in color. **H)** Histograms of pairwise correlations with nearby neighbors. *x* and *y* axes are in μm. Bars to the right of each figure indicate correlation values in colors.

### Fig. S33

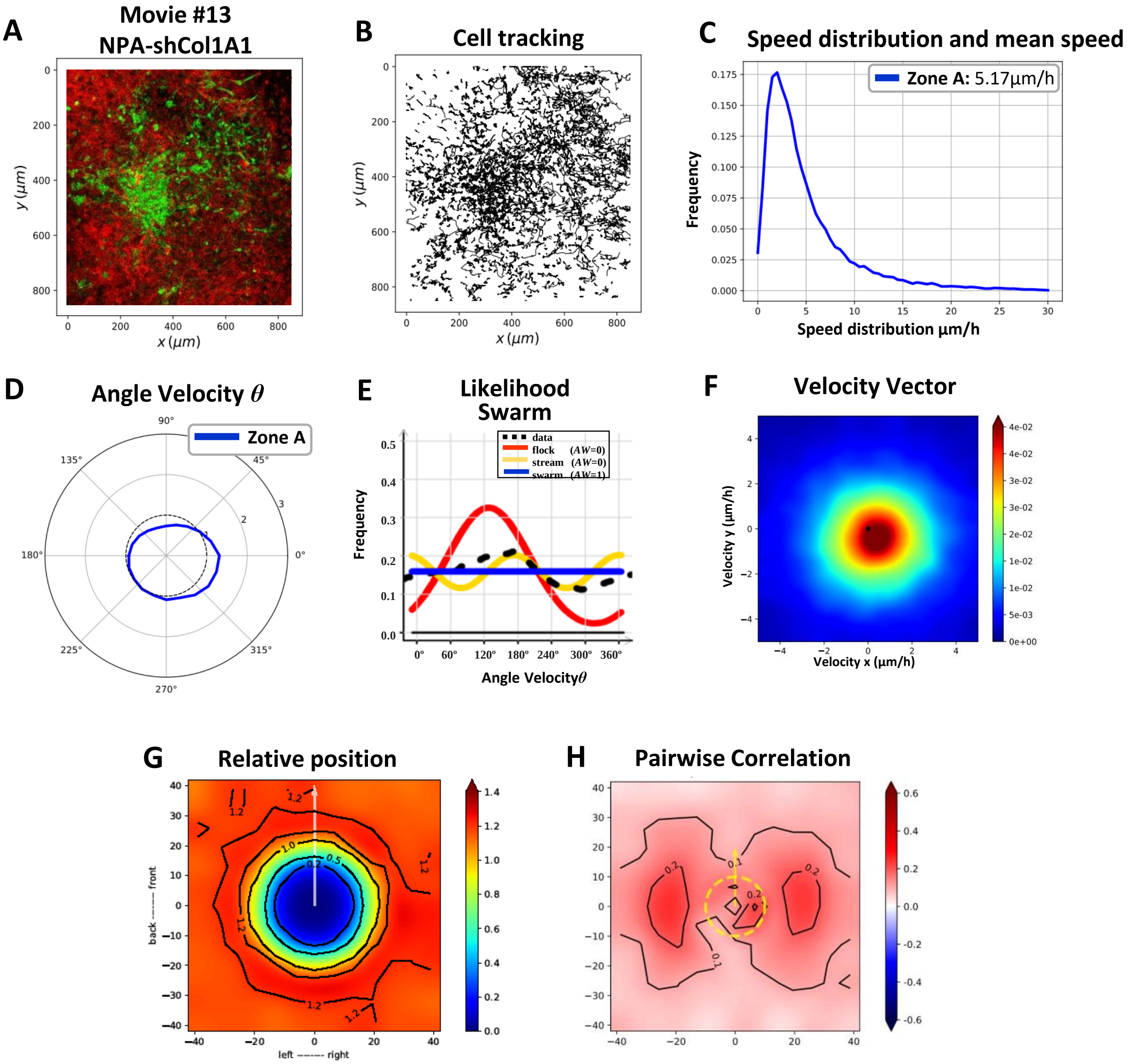


**Fig. S33: Migration analysis of NPAshCol1A1 explant model using confocal time-lapse imaging; Movie # 13 A)** Representative time-lapse confocal images of the tumor border for movie #9. **B)** Tracking analysis of individual cell paths performed using the Track-Mate plugin from Image-J. **C)** Speed distribution and mean speed (*µm/h*). **D)** Angle Velocity distribution analysis ($\vartheta$) performed. **E)** Likelihood analysis histogram to classify the dynamic motion pattern as a swarm. **F)** Heat map plot of the distribution of the velocity vectors. **G)** Histogram plot showing the relative position with nearby neighbors. *x* and *y* axes are in µm. Bars to the right of each figure indicate probability values in color. **H)** Histograms of pairwise correlations with nearby neighbors. *x* and *y* axes are in µm. Bars to the right of each figure indicate correlation values in colors.

**Fig. S34**

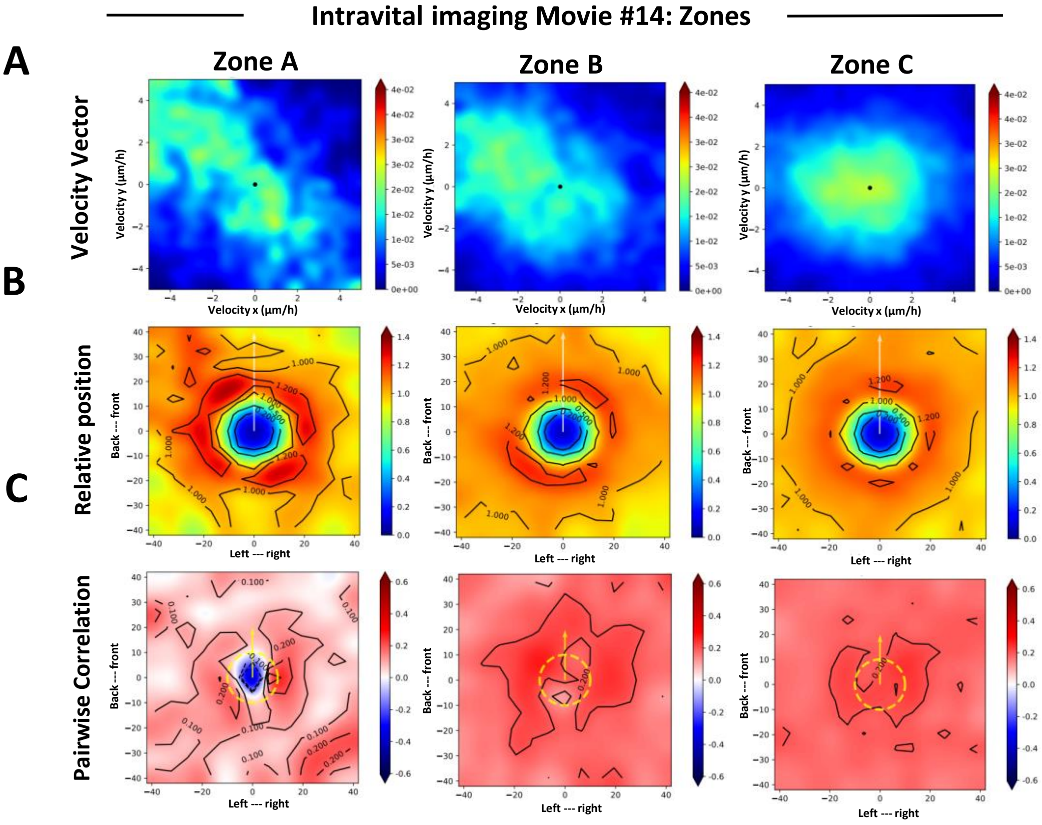


**Fig. S34. Migration analysis in intravital imaging Movie#14**

**A)** Heat map plot of the distribution of the velocity vectors in each zone. **B)** Histogram plot showing the relative position with nearby neighbors for each zone. x and y axes are in µm. Bars to the right of each figure indicate probability values in color. **C)** Histograms of pairwise correlations with nearby neighbors for each zone. x and y axes are in µm. Bars to the right of each figure indicate correlation values in colors.

Fig. S35

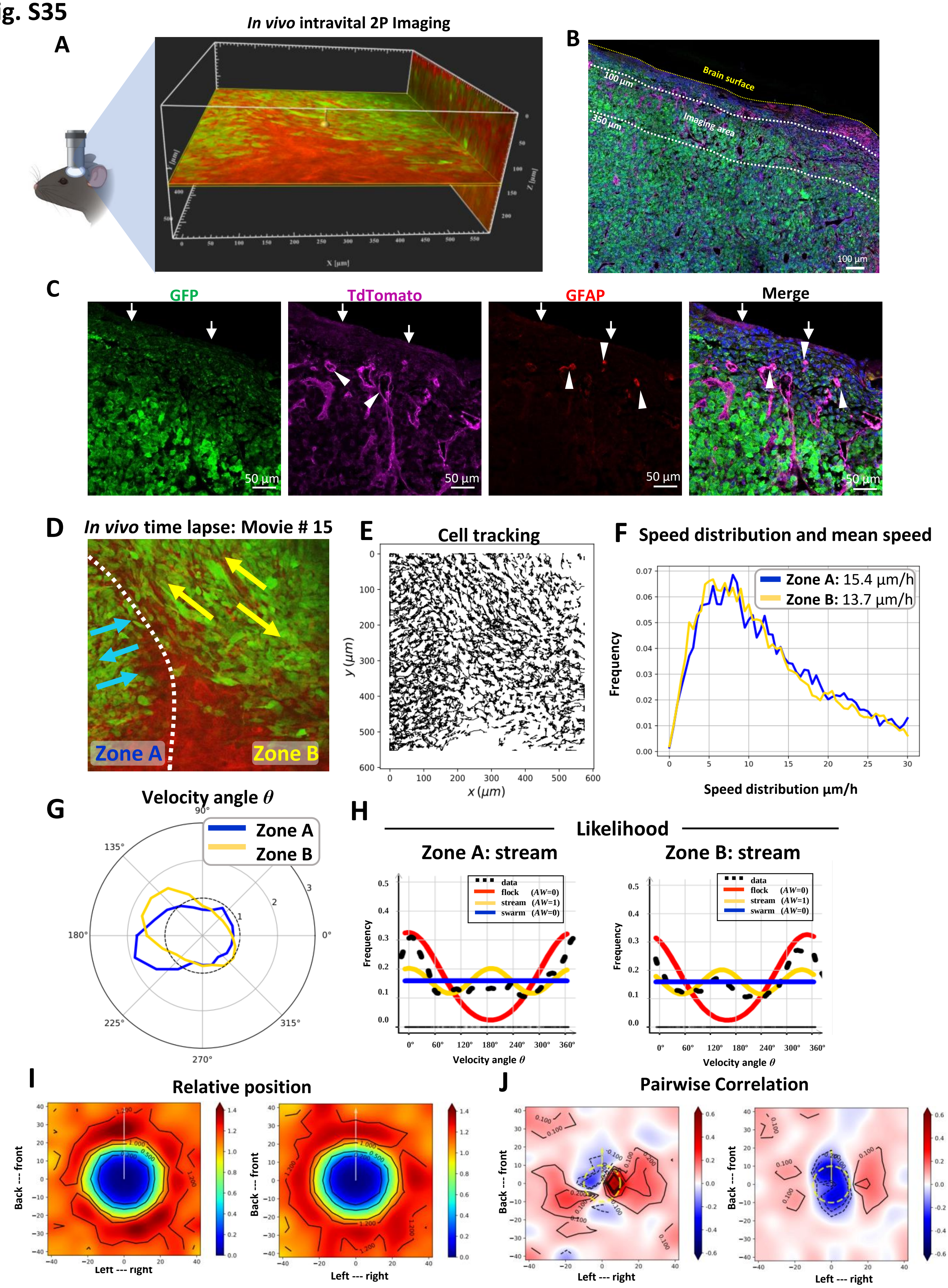
A
In vivo intravital 2P Imaging
X [µm]
Y [µm]
Z [µm]
B
Brain surface
100 µm
350 µm
Imaging area
100 µm
C
GFP
TdTomato
GFAP
Merge
50 µm
D
In vivo time lapse: Movie # 15
Zone A
Zone B
E
Cell tracking
y (µm)
x (µm)
F
Speed distribution and mean speed
Zone A: 15.4 µm/h
Zone B: 13.7 µm/h
Frequency
Speed distribution µm/h
G
Velocity angle θ
Zone A
Zone B
H
Likelihood
Zone A: stream
Zone B: stream
data
flock (AW=0)
stream (AW=1)
swarm (AW=0)
Frequency
Velocity angle θ
I
Relative position
Back --- front
Left --- right
J
Pairwise Correlation
Back --- front
Left --- right

**Fig. S35. Intravital two-photon imaging setup for time-lapse acquisition and immunofluorescence of coronal section of the imaging area: Movie #15**

**A)** shows an image generated using the Orthoslicer 3D function of the Imaris viewer software illustrating the plane used for time-lapse imaging. The image depicts the imaging plane and its distance from the brain's surface. For this experiment, we took 248 x-y frames from the brain's surface at a depth increment of 1 µm (voxel size=1) and resolution of 1024x1024 pixels. XYZ axes of the orthoslicer3D image are shown in white color (596x596x248 µm). The imaging plane was at 140 µm from the brain surface. Normal brain parenchyma Is shown in red and tumor cells in green. **B)** Following *in vivo* intravital imaging, the animal was perfused, and the brain was fixed and embedded in paraffin. Immunofluorescence was performed on 5 µm thick sections. The brain surface is marked in yellow, and the imaging area is marked in between white lines (100 to 350 µm in depth). Sections were stained with anti-GFP (green) to detect tumor cells, anti-tdTomato (pink) to detect brain parenchyma cells, and anti-GFAP (red) to detect astrocytes. Scale bar: 100 µm. **C)** From left to right: Tumor cells (GFP+) growing within the cortex. The brain parenchymal cells (anti-tdTomato+) are present in the brain surface, blood vessels, as well as in between tumor cells. GFAP+ cell processes can be found within the tumor. The merged channel shows the tumor growing mainly below the brain surface and within the brain parenchyma. The tumor cells are interacting with the brain parenchyma (blood vessels, astrocytes). Scale bar: 50 µm. **D)** Single representative time-lapse two-photon image of glioma cell invasion *in vivo* (Movie #15), showing Zones A, B. **E)** Individual cell paths trajectories of this *in vivo* time-lapse experiment. **F)** Histogram of speed distribution and mean speed (µm/hr) in Zones A, B. **G)** Angle Velocity distribution for each zone's in the *in vivo* time-lapse. The Angle Velocity of each cell is denoted θ. The plot shows the proportion of cells moving in the angle direction θ for each zone. **H)** Likelihood analysis of the dynamic patterns in each zone of Movie #15 by intravital imaging. Graph of frequency distribution *ρ flock* (red), *ρ stream* (yellow) and *ρ swarm* (blue). The estimation of the black line (data) uses a *non-parametric* estimation. AW: 0 or AW:1. **I)** Histogram plot showing the relative position with nearby neighbors for each zone; *x* and *y* axes are in µm. Bars to the right of each figure indicate probability values in color. **J)** Histograms of pairwise correlations with nearby neighbors for each zone; *x* and *y* axes are in µm. Bars to the right of each figure indicate correlation values in colors.

Fig. S36

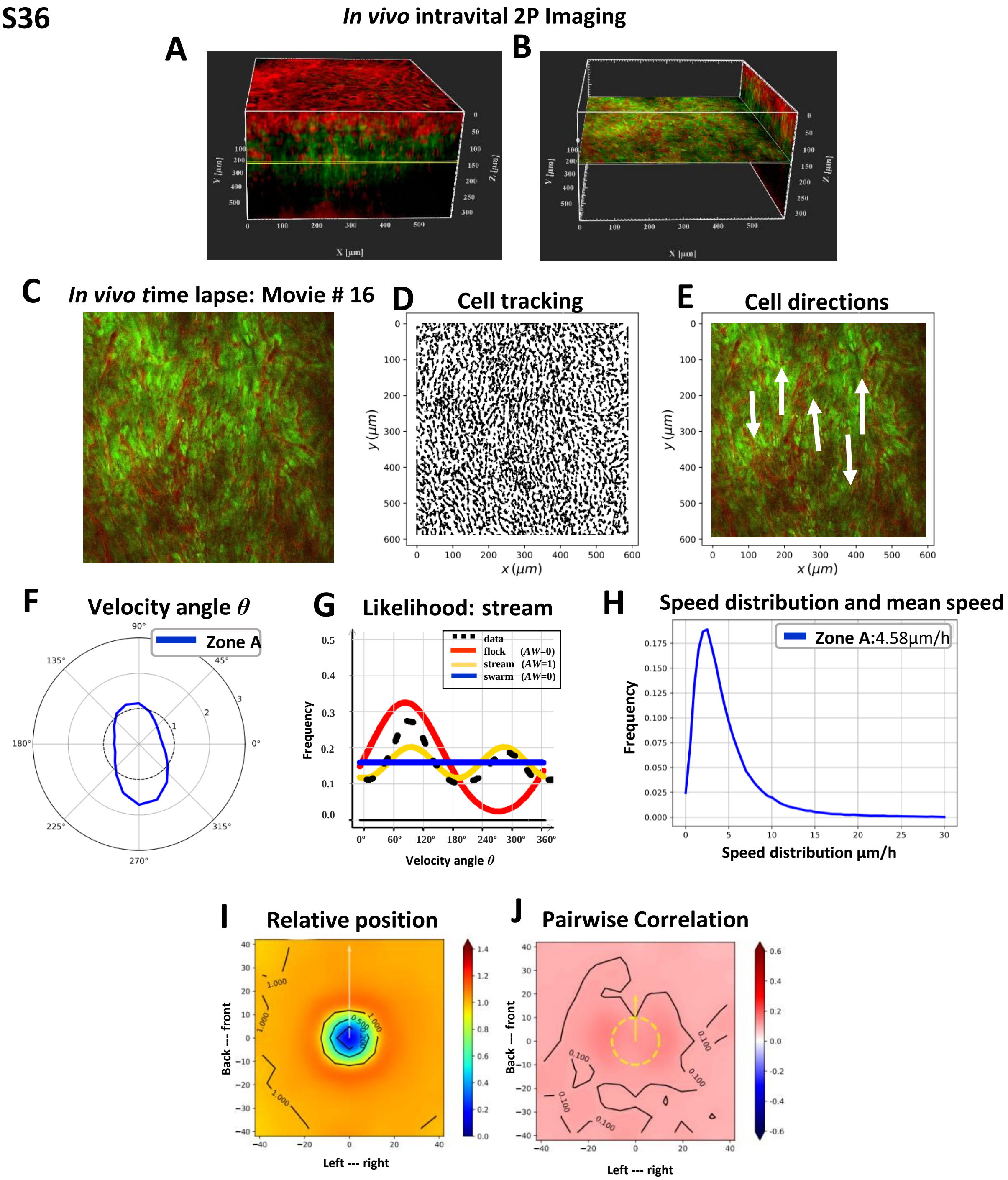

In vivo intravital 2P Imaging
A
B
X [µm]
Y [µm]
Z [µm]
C
In vivo time lapse: Movie # 16
D
Cell tracking
E
Cell directions
y (µm)
x (µm)
F
Velocity angle θ
Zone A
G
Likelihood: stream
data
flock (AW=0)
stream (AW=1)
swarm (AW=0)
Frequency
Velocity angle θ
H
Speed distribution and mean speed
Zone A:4.58µm/h
Frequency
Speed distribution µm/h
I
Relative position
Back --- front
Left --- right
J
Pairwise Correlation
Back --- front
Left --- right

**Fig. S36. Invasion analysis of NPA glioma dynamics within the tumor core using Intravital two-photon microscopy *in vivo:* Movie #16**

**A)** High-resolution 3D Z-stacks (spanning up to 330 µm depth from the brain's surface) were imported into the Imaris viewer to reconstruct the 3D image. To do so, 330 x-y frames from the brain's surface were taken at a depth increment of 1 µm (voxel size=1) and resolution of 1024x1024 pixels. The XYZ axes of the 3D image are shown in white color (each comprising an area of 596x596x330 µm), and the yellow line (at 145 µm depth) shows the exact imaging plane used to obtain the *in vivo* time-lapse data for movie #16. **B)** shows an image generated with the Orthoslicer3D from Imaris using the 3D reconstruction data shown in **(A)**; this illustrates the depth of the imaging plane and its distance from the brain surface. Normal brain parenchyma is shown in red, and tumor cells are shown in green. **C)** Single representative time-lapse two-photon image of glioma cells within the tumor core *in vivo* taken at a depth of 145 µm. **D)** Individual cell paths trajectories of this *in vivo* time-lapse experiment. **E)** Preferred directions of cells within the tumor core superimposed onto a representative time-lapse image. **F)** Angle Velocity distribution for the migration of cells growing within the tumor core for the *in vivo* time-lapse dataset. Angle Velocity for each cell is denoted θ. The plot shows the proportion of cells moving in the angle direction θ. **G)** The frequency distribution corresponding to Angle Velocity(θ) is shown by dotted black lines. Likelihood analysis of the dynamic patterns within the tumor core (Movie#16). Graph of frequency distribution is shown as *ρ flock* (red), *ρ stream* (yellow) and *ρ swarm* (blue). The estimation of the black line (data) uses *non-parametric* estimation. AW: 0 or AW:1. **H)** Histogram of speed distribution and mean speed in µm/hr for the full time-lapse movie considered as a single zone named Zone A. **I)** Histogram plot showing the relative position of nearest neighbors for Zone A; *x* and *y* axes are in µm. Bars to the right of each figure indicate probability values in color. **J)** Histograms of pairwise correlations with nearby neighbors for Zone A; *x* and *y* axes are in µm. Bars to the right of each figure indicate correlation values in colors.

Fig. S37

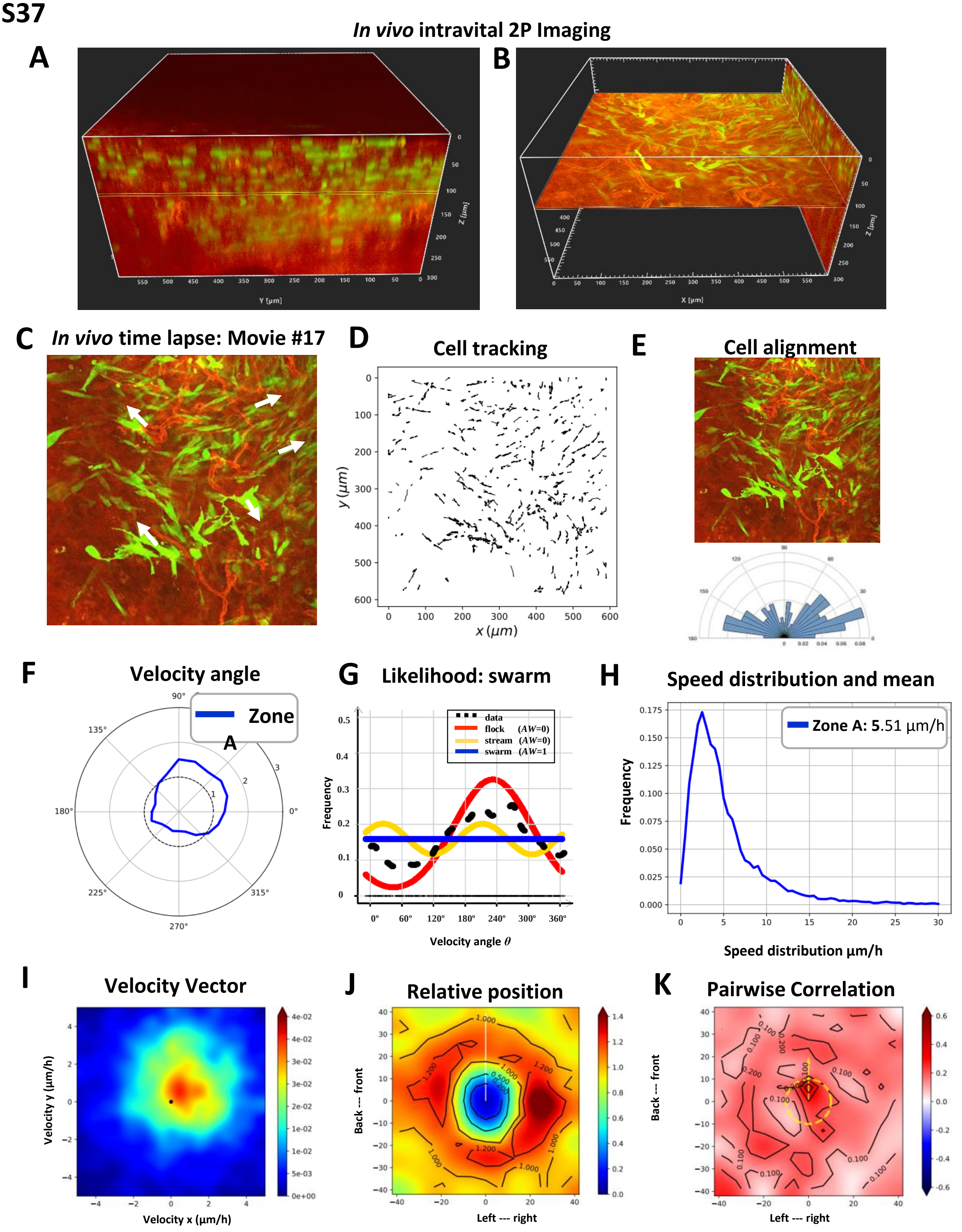
In vivo intravital 2P Imaging
A
B
Y [µm]
X [µm]
Z [µm]
C In vivo time lapse: Movie #17
D Cell tracking
y (µm)
x (µm)
E Cell alignment
F Velocity angle
Zone A
G Likelihood: swarm
data
flock (AW=0)
stream (AW=0)
swarm (AW=1
Frequency
Velocity angle θ
H Speed distribution and mean
Zone A: 5.51 µm/h
Frequency
Speed distribution µm/h
I Velocity Vector
Velocity y (µm/h)
Velocity x (µm/h)
J Relative position
Back --- front
Left --- right
K Pairwise Correlation
Back --- front
Left --- right

**Fig. S37. Invasion analysis of NPAshCol1A1 glioma dynamics using Intravital two-photon microscopy *in vivo:* Movie #17**

**A)** High-resolution 3D Z-stacks (spanning up to 300 µm depth from the brain's surface) were imported into the Imaris viewer to reconstruct the 3D image. To do so, 300 x-y frames from the brain's surface were taken at a depth increment of 1 µm (voxel size=1) and resolution of 1024x1024 pixels. The XYZ axes of the 3D image are shown in white color (each comprising an area of 596x596x300 µm), and the yellow line (at 110 µm depth) shows the exact imaging plane used to obtain the *in vivo* time-lapse data for movie #17. **B)** shows an image generated with the Orthoslicer3D from Imaris using the 3D reconstruction data shown in **(A)**; this illustrates the depth of the imaging plane and its distance from the brain surface. Normal brain parenchyma is shown in red, and tumor cells are shown in green. **C)** Single representative time-lapse two-photon image of NPAshCol1A1 glioma cells within the tumor core *in vivo* taken at a depth of 110 µm. Preferred directions of cells within tumor core superimposed onto a representative time-lapse image. **D)** Individual cell paths trajectories of this *in vivo* time-lapse experiment. **E)** Angle histogram plots show the random alignment of NPAshCol1A1 glioma cells lacking oncostreams *in vivo*. Angle histograms correspond to the representative time-lapse two-photon image on the top. **F)** Angle Velocity distribution for the migration of NPAshCol1A1 glioma cells growing within the tumor core for the *in vivo* time-lapse dataset. The Angle Velocity of each cell is denoted θ. The plot shows the proportion of cells moving in the angle direction θ. **G)** The frequency distribution corresponding to Angle Velocity(θ) is shown by dotted black lines. Likelihood analysis of the dynamic patterns within the tumor core (Movie#17). Graph of frequency distribution is shown as *ρ flock* (red), *ρ stream* (yellow) and *ρ swarm* (blue). The estimation of the black line (data) uses a *non-parametric* estimation. AW: 0 or AW:1. **H)** Histogram of speed distribution and mean speed in µm/hr for the full time-lapse movie considered as a single zone named Zone A. **I)** Heatmap plot of the distribution of velocity vectors within Zone A. **J)** Histogram plot showing the relative position of nearest neighbors for Zone A; *x* and *y* axes are in µm. Bars to the right of each figure indicate probability values in color. **K)** Histograms of pairwise correlations with nearby neighbors for Zone A; *x* and *y* axes are in µm. Bars to the right of each figure indicate correlation values in colors.
